# *Trem2* promotes anti-inflammatory responses in microglia and is suppressed under pro-inflammatory conditions

**DOI:** 10.1101/449884

**Authors:** Wenfei Liu, Orjona Taso, Rui Wang, Sevinc Bayram, Pablo Garcia-Reitboeck, Anna Mallach, William D. Andrews, Thomas M. Piers, Andrew C. Graham, Juan A. Botia, Jennifer M. Pocock, Damian M. Cummings, John Hardy, Frances A. Edwards, Dervis A. Salih

## Abstract

Genome-wide association studies have reported that, amongst other microglial genes, variants in *TREM2* can profoundly increase the incidence of developing Alzheimer’s disease (AD). We have investigated the role of TREM2 in primary microglial cultures from wild type mice by using siRNA to decrease *Trem2* expression, and in parallel from knock-in mice heterozygous or homozygous for the *Trem2* R47H AD risk variant. The prevailing phenotype of *Trem2* R47H knock-in mice was decreased expression levels of *Trem2* in microglia, which resulted in decreased density of microglia in the hippocampus. Overall, primary microglia with reduced *Trem2* expression, either by siRNA or from the R47H knock-in mice, displayed a similar phenotype. Comparison of the effects of decreased *Trem2* expression under conditions of LPS pro-inflammatory or IL-4 anti-inflammatory stimulation revealed the importance of *Trem2* in driving a number of the genes up-regulated in the anti-inflammatory phenotype. RNA-seq analysis showed that IL-4 induced the expression of a programme of genes including *Arg1* and *Ap1b1* in microglia, which showed an attenuated response to IL-4 when *Trem2* expression was decreased. Genes showing a similar expression profile to *Arg1* were enriched for STAT6 transcription factor recognition elements in their promoter, and *Trem2* knockdown decreased levels of the transcription factor STAT6. LPS-induced pro-inflammatory stimulation suppressed *Trem2* expression, thus preventing TREM2’s anti-inflammatory drive. Given that anti-inflammatory signaling is associated with tissue repair, understanding the signaling mechanisms downstream of *Trem2* in coordinating the pro- and anti-inflammatory balance of microglia, particularly mediating effects of the IL-4-regulated anti-inflammatory pathway, has important implications for fighting neurodegenerative disease.

**Graphical abstract:** 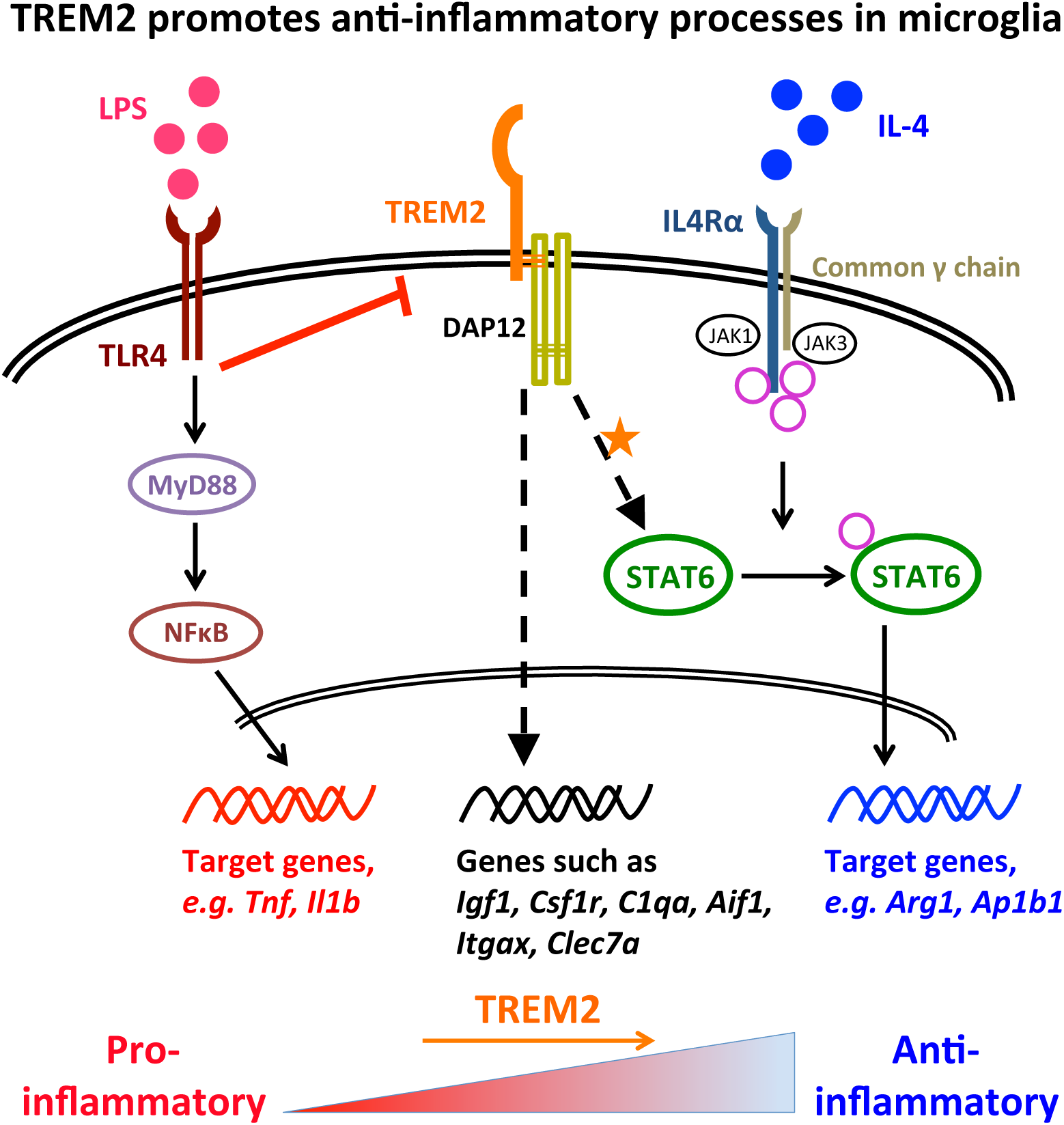

## Introduction

Genome-wide association studies (GWAS) continue to reveal variants in immune/microglia-related genes that significantly increase the risk of developing Alzheimer’s disease (AD). These genes now include *TREM2*, *CD33*, *ABI3, PLCG2* and *SPI1* (1–8). These studies particularly emphasize an important role of microglia in AD progression. However, the question of how microglial activation contributes to AD development is still far from resolved. Nor is it clear how to use these findings to leverage microglial pathways for drug discovery.

Triggering receptor expressed on myeloid cells 2 (TREM2) is a transmembrane protein containing a single immunoglobulin-superfamily (Ig-SF) domain. TREM2 expression was first observed in macrophages and dendritic cells, but not granulocytes or monocytes (9–11). It has been reported to be widely expressed in various cell types derived from the myeloid lineage, such as bone osteoclasts (12, 13), and lung and peritoneal macrophages (14). In the brain, *TREM2* is predominantly expressed by microglia (15–18). In 2013, two independent GWAS studies (2, 4) highlighted a rare variant rs75932628-T in exon 2 of the *TREM2* gene, causing an arginine-to-histidine substitution at amino acid position 47 (R47H), to be highly significantly associated with increased AD risk. The risk from this variant has a comparable effect size to the *APOEε4* allele (around 3-fold increased AD risk; 4, 19-22). Furthermore, the R47H variant was investigated in other neurodegenerative diseases and was also found to be associated with frontotemporal dementia, Parkinson’s disease, and amyotrophic lateral sclerosis (23, 24), suggesting a more widespread role for TREM2 in maintaining microglial function/survival throughout the brain. In addition to the R47H variant, further *TREM2* variants have been revealed to be risk factors of AD or other CNS diseases, such as R62H (8, 20), H157Y (25), Q33X, T66M and Y38C (26, 27). All of the *TREM2* genetic linkage studies with neurodegenerative diseases suggest an important contribution of impaired or decreased TREM2 function in regulating microglial functions contributing to disease progression.

Evidence from *in vitro* studies has accumulated to support the role of TREM2 in restricting inflammation and promoting phagocytosis. It was first reported in a study using primary microglia transduced with flag epitope tagged TREM2 that TREM2 stimulated microglial chemotaxis, and phagocytosis of fluorescent beads (17). The same study also showed that *Trem2* knockdown via a lentiviral strategy increased pro-inflammatory gene expression and decreased phagocytosis of labeled apoptotic neuronal membranes in primary microglia stressed with co-culture of apoptosis-induced primary neurons. TREM2 was also found to suppress LPS-induced pro-inflammatory markers up-regulated in primary alveolar macrophages *in vitro* (28), and the BV-2 microglial cell line (29). With particular relevance to AD, phagocytosis of amyloid beta (Aβ) was found to be impaired due to lack of or reduced mature TREM2 cell surface expression in studies with primary microglia from *Trem2* knock-out mice and mutant *Trem2* microglial cell lines (30, 31). Recently, it was further shown that apolipoproteins, lipids and Aβ itself acted as ligands of TREM2 to facilitate microglial phagocytosis and clearance of Aβ and apoptotic neurons (32–35). In general, these studies suggest that *Trem2* deficiency exacerbates microglial pro-inflammatory activation in response to various stimuli, and accompanies impaired phagocytosis. However, other cellular functions of TREM2, particularly anti-inflammatory or alternative activation, and the molecular mechanisms mediating the actions of TREM2, are not well understood.

In this study, we investigated microglia in mice carrying the *Trem2* R47H point mutation from the Jackson laboratories (*Trem2* R47H knock-in mice, KI mice), alongside wild type (WT) littermates, and then using primary cultures in parallel with acute knockdown of *Trem2* expression using RNAi. The primary effect reported in the R47H mice was a reduction of *Trem2* expression due to altered splicing, and this effect was also observed in independent *Trem2* R47H KI mice (36, 37). The changes in microglial number and activation in this mouse will thus be a combination of the partial knockdown of *Trem2* and any effect of the R47H mutation. Hence this is effectively an *in vivo* model of down regulation of TREM2 activity, through the combination of these two factors. We then used isolated primary mouse microglial cultures with acute siRNA-induced knockdown of *Trem2* expression and compared these to microglia carrying the R47H mutation, especially when polarized towards a pro-inflammatory or anti-inflammatory phenotype. We find that decreased levels of *Trem2* have a particularly strong effect on the IL-4 mediated anti-inflammatory gene expression profile, in part by acting through the STAT6 transcription factor.

## Results

### Decreased microglial density and down-regulated *Trem2* expression in the hippocampus of *Trem2* R47H knock-in mice

We aimed to investigate the effects of the *Trem2* R47H variant associated with AD in microglia and compare these to acute *Trem2* ‘loss-of-function’ using siRNA in microglia. We employed mice with knock-in of a R47H point mutation into the endogenous mouse *Trem2* gene (Jackson Laboratory, stock # 27918). The primary effect in these *Trem2* R47H KI mice was a substantial and chronic gene dose-related decrease in *Trem2* expression in hippocampal tissue from the mice (Fig. 1A). This has been shown to be due to a splice effect that seems to occur in mice but not in humans (36). Reduced expression of *Trem2* was also seen in an independently generated *Trem2* R47H KI model (37). We can thus consider this R47H mouse line as a useful model for *Trem2* down-regulation *in vivo* to compare to our findings with RNAi-mediated knockdown of *Trem2* presented below. We quantified microglial density in different layers of the hippocampal CA1 region at 4 months of age, using immunohistochemistry with an antibody against IBA1, and found that homozygous *Trem2* R47H KI mice displayed a significant decrease in the number of microglia compared to WT littermates, which was consistent in all four CA1 layers (two-way ANOVA: main effect of genotype p<0.01; Fig. 1B and 1C). Interestingly, we also observed a significant decrease of CD68 positive microglia in the homozygous *Trem2* R47H mice (two-way ANOVA: main effect of genotype p<0.0001). This change was not primarily due to the total microglia number change because the proportion of CD68 positive microglia also showed a substantial decrease (two-way ANOVA: main effect of genotype p<0.001; Fig. 1B and 1C). This result suggests that, in addition to slightly reduced microglial density in the hippocampus of homozygous *Trem2* R47H mice, phagocytosis would also be reduced in the surviving microglia.

**FIGURE 1.**
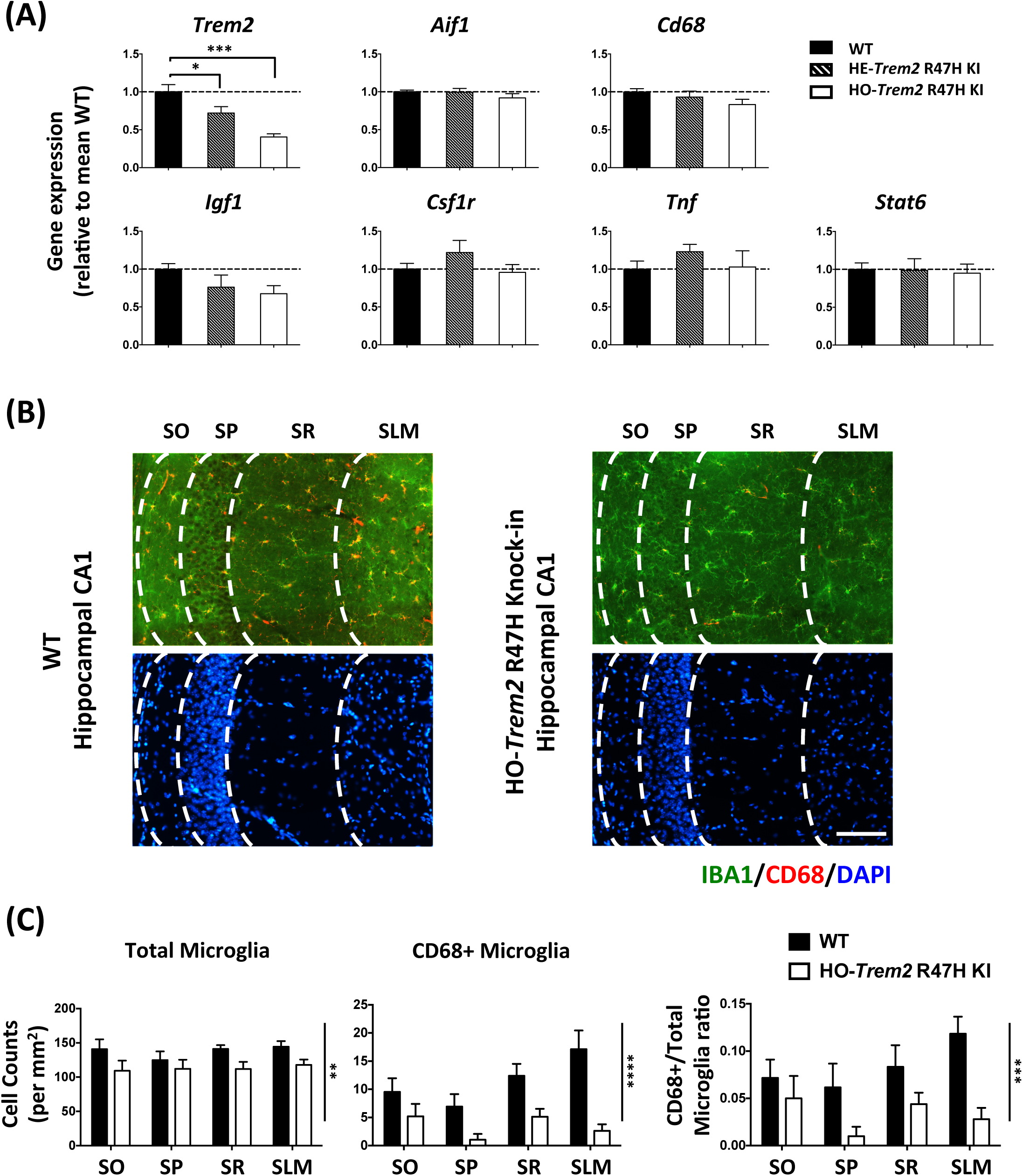
*Trem2* R47H KI mice (4 months old) showed significant down-regulation of *Trem2* expression, and decreased microglia density and CD68 levels in the hippocampus. (A) Gene expression in hippocampal homogenates from homozygous (HO) and heterozygous (HE) *Trem2* R47H KI mice relative to *Rps28* and then expressed relative to WT within the batch. N=6-7 mice per group. Data shown as mean ± SEM. One-way ANOVA showed a significant main effect of genotype for *Trem2* expression only, and so Sidak’s *post hoc* tests were used to test pairwise significance between the three genotypes; * p<0.05, ** p<0.01, *** p<0.001. (B) Representative images showing AIF1/IBA1 and CD68 immuno-staining of microglia with DAPI in the hippocampal CA1 regions from WT and HO-*Trem2* R47H KI mice. The layers from left are SO (*stratum oriens*), SP (*stratum pyramidale*), SR (*stratum radiatum)*, and SLM (*stratum lacunosum-moleculare*). Scale bar: 100 µm. (C) Left panel: total microglia density in all four CA1 layers was decreased in HO-*Trem2* R47H KI mice. Middle panel: density of CD68 positive microglia decreased in HO-*Trem2* R47H KI mice. Right panel: proportion of CD68 positive microglia was significantly lower in HO-*Trem2* R47H KI mice. N=5-6 mice per group. Data shown as mean ± SEM. Two-way ANOVA with significant main effect of *Trem2* genotype indicated with vertical lines, no significant interactions were seen between hippocampal layer and *Trem2* genotype; * p<0.05, ** p<0.01, *** p<0.001, **** p<0.0001.

We next assessed the microglial activation state in the hippocampus of R47H *Trem2* KI mice at 4 months of age by investigating the expression of a battery of microglial genes covering pro- and anti-inflammatory functions, and other microglial processes (Fig. 1A). Overall, at 4 months of age, a number of genes did not show any significant difference in the hippocampus of the *Trem2* R47H KI mice compared to WT littermates (Fig. 1A). It is important to note that in the hippocampal tissue, the gene expression in microglia is greatly diluted by the much greater contribution of other cell types and so small changes in gene expression will be much more difficult to detect than in enriched microglia. In summary, the *Trem2* R47H KI mice showed that reduced *Trem2* expression resulted in predominantly a phenotype of a slight but significant decrease in microglial numbers and also lower CD68-positive microglia in the CNS environment of healthy young adult mice. This may prime the mouse for neurodegeneration later in life with ageing or other environmental factors.

### Primary microglial cultures with acute *Trem2* knockdown and reduced *Trem2* expression in *Trem2* R47H microglia

To investigate the functional potential of microglia with reduced *Trem2* expression using siRNA or in microglia carrying the *Trem2* R47H variant, we established an *in vitro* model where primary microglia were kept in mixed glial culture conditions to develop for around 21 days before isolation (Fig. 2). The microglia isolated from the *Trem2* R47H KI mice show decreased *Trem2* expression throughout the life of the mouse, without the need for transfection. The purity of our primary microglial culture 24 hr after isolation was validated via immunocytochemistry against IBA1 (97.2±0.3% IBA1-positive cells; Fig. 3A and Fig. S1). In order to validate the potential of the microglial culture to be polarized in the direction of either the pro- or anti-inflammatory phenotypes, LPS or IL-4 was applied respectively, and expression of typical pro-inflammatory (*Tnf* and *Il1b*) or anti-inflammatory (*Arg1* and *Tgfb1*) genes were analyzed via RT-qPCR (Fig. 3B). In particular, IL-4 treatment induced a dramatic time-dependent up-regulation of the anti-inflammatory gene *Arg1* (Fig. 3B). *Arg1* expression was undetectable in our non-stimulated microglia, but reached levels well above detection threshold at 24 hr after IL-4 treatment, and by 48 hr expression levels had further doubled. There is no substantial change in the proportions of microglia versus other cell types during 72 hr of culture for WT compared to homozygous *Trem2* R47H KI microglia, under basal conditions, and also in response to LPS and IL-4 treatment (Fig. S1), although there is some cell death (Fig. S2).

**FIGURE 2.**
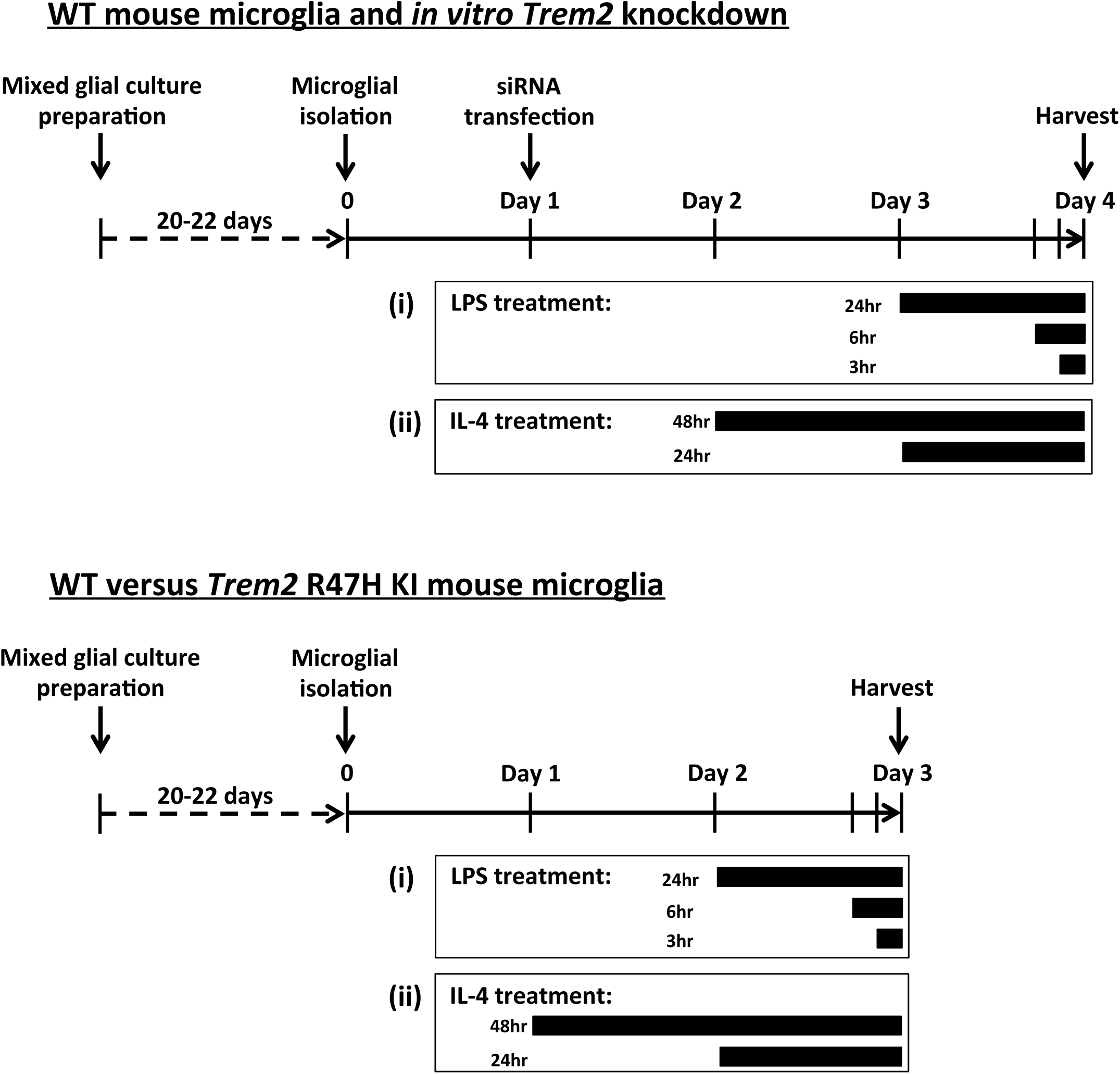
The *in vitro* primary microglial models. Schedule of primary microglia culture from WT or *Trem2* R47H KI mice, and treatment with *Trem2* siRNA, control non-targeting siRNA, LPS or IL-4.

**FIGURE 3.**
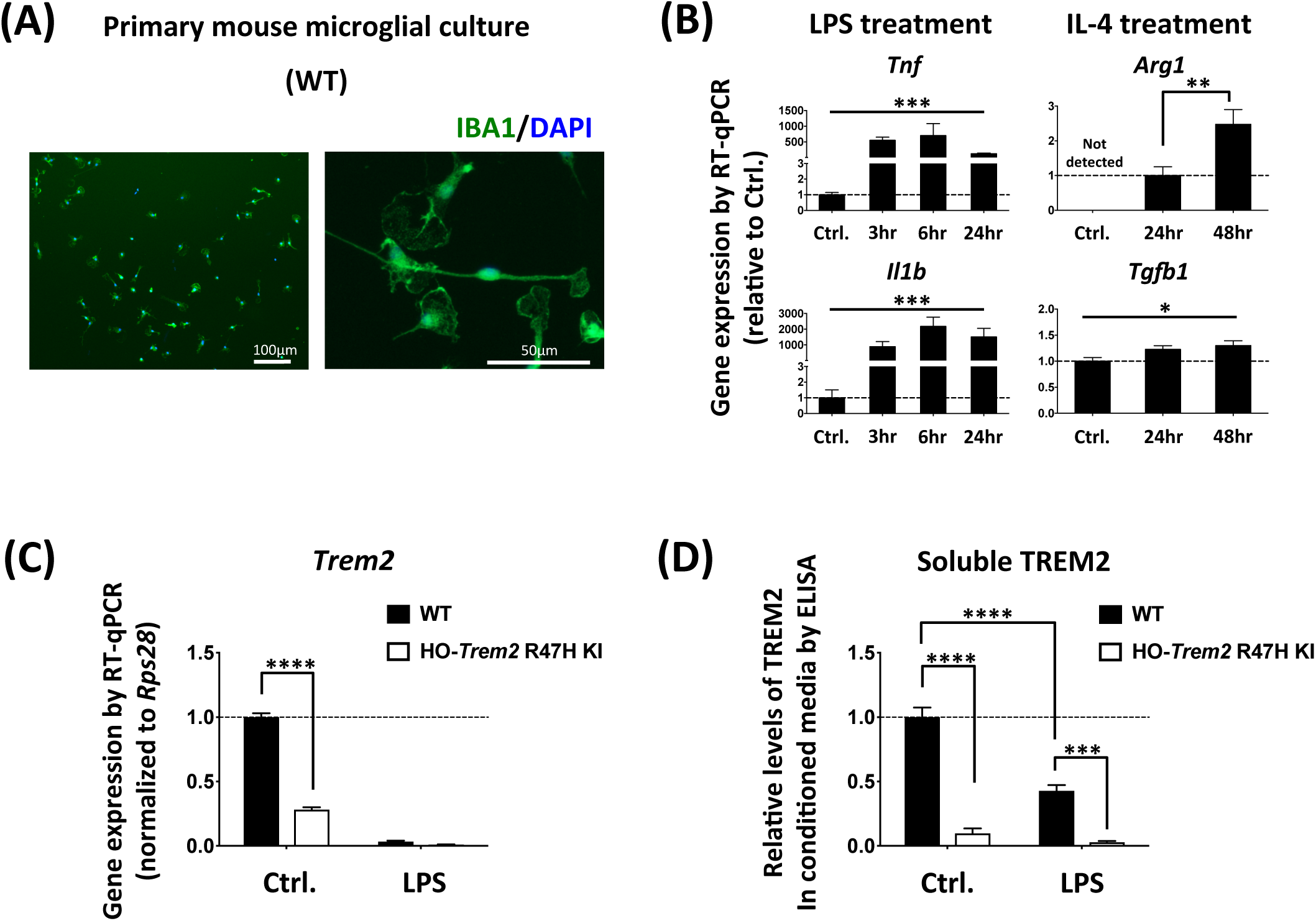
LPS suppression of *Trem2* in primary microglia. (A) Left panel: primary microglia stained with an antibody against IBA1 and DAPI. Right panel: a higher-magnification image showing the morphology of microglia *in vitro*. (B) Pro- and anti-inflammatory gene expression changes in primary microglia with either LPS or IL-4 treatment. N=3-6 independent experiments. Data shown as mean ± SEM. One-way ANOVA showed a significant effect of treatment for *Tnf*, *Il1b* and *Tgfb1*. *Arg1* was not detected under control conditions, and so Student’s *t*-test was used to test pairwise significance between 24 and 48 hr of IL-4 treatment; * p<0.05, ** p<0.01, *** p<0.001. (C) Expression of *Trem2*. N=6 mice per genotype. Data shown as mean ± SEM. Two-way ANOVA with significant main effect of both LPS-treatment and genotype, and a significant interaction between treatment and genotype, so Sidak’s *post hoc* tests were performed to test pairwise significance between the genotypes within a treatment group, marked above individual groups. The interaction reflects a lack of effect of genotype in the LPS treated microglia. *** p<0.001. (D) ELISA analysis of soluble TREM2 levels in the microglial supernatants. Soluble TREM2 levels were normalized to the expression levels of three housekeeping genes, *H3f3b*, *Arf1* and *Rps28* assessed by RT-qPCR. N=3-4 independent microglial preparations. Data shown as mean ± SEM. Two-way ANOVA with significant main effect of both LPS-treatment and genotype, and a significant interaction between treatment and genotype (p=0.0002). Thus Sidak’s *post hoc* tests were performed to test pairwise significance between the four groups, marked above individual groups. *** p<0.001, **** p<0.0001.

Given the reduced number of microglia in the hippocampus of *Trem2* R47H KI mice (Fig. 1), we assessed apoptosis in primary microglia from *Trem2* R47H KI versus WT mice by assessing Annexin V binding using FACS analysis (Fig. S2). As expected, we saw increased cell death in *Trem2* R47H KI microglia (14.6% Annexin V-positive, propidium iodide-negative) compared to microglia from WT littermates (7.3% Annexin V-positive, propidium iodide-negative). The increase in cell death is likely due to reduced expression of *Trem2,* consistent with previous findings with *Trem2* knock-out and knock-down (38–40).

To develop a parallel approach to reduce *Trem2* expression, three different siRNA sequences were tested in parallel to a non-targeting control siRNA (Fig. S3). The most effective siRNA (sequence 2) gave around 80% knockdown of *Trem2* mRNA levels compared to the non-targeting siRNA control. Thus we chose this sequence for the majority of RNAi experiments in primary microglia, and confirmed that key findings were dependent on the levels of *Trem2* expression using siRNA sequence 3, which produced 50-60% knockdown of *Trem2* (Fig. S3).

### Acute *Trem2* knockdown in primary microglia is effective and impairs phagocytosis, validating the preparation

To assess the effect of *Trem2* knockdown by siRNA in this system, we initially tested the effects on phagocytosis based on previous studies showing impaired phagocytosis with *Trem2* manipulation (17, 30, 41). We found that microglia with *Trem2* knockdown showed a significant reduction (48.1±9.1%) of phagocytosis of *E. coli*, conjugated to pHrodo (a pH-sensitive fluorescent dye), compared to the non-targeting siRNA treated microglia (Fig. S4), confirming previous reports and further validating this cell system.

### Reduced *Trem2* causes transcriptional changes in primary microglia

To understand the genetic pathways regulated by TREM2, we next investigated the subsequent gene expression changes in microglia caused by *Trem2*-knockdown under non-stimulated conditions. We then found that *Trem2* knockdown resulted in a significant down-regulation of a number of microglial genes at 72 hr after siRNA transfection including *C1qa, Cd68, Csf1r, Igf1, Pik3cg, Spi1, Tnf and Tgfb1* (Fig. S5). Overall, acute *Trem2* knockdown produced a significant change in the gene expression profile of microglia under non-stimulated conditions, which might result in a shift of the microglial functional phenotype.

### LPS stimulation dramatically suppresses *Trem2* gene expression in primary microglia

An important outstanding question is how *Trem2* is regulated, so to investigate this, microglia were stimulated with LPS to induce classical pro-inflammatory responses. We first found that, in both the wild type and *Trem2* R47H KI cells, LPS treatment strongly down-regulated *Trem2* expression (Fig. 3C and Fig. S6), which was consistent with the previous findings in a variety of cell preparations (16, 42). In addition, the soluble TREM2 protein level in the conditioned medium in response to LPS was decreased for the microglia from the wild type littermates (Fig. 3D), and the level of TREM2 protein in the medium conditioned by cells from the homozygous *Trem2* R47H KI mice was also decreased, supporting the transcriptional findings. This suggests that pro-inflammatory conditions inhibit microglial *Trem2* expression, which might result in a state of temporary or chronic ‘TREM2-deficiency’ in microglia.

The expression levels of *Tnf* and *Il1b,* two typical pro-inflammatory markers, were greatly increased in our LPS-treated microglia from as early as 3 hr after LPS application. As expected, considering so little *Trem2* remained under these conditions, knockdown of *Trem2* by siRNA had little effect on LPS-induced gene expression with up-regulation of *Tnf* or *Il1b* being unchanged at the transcriptional level with *Trem2* knockdown compared to control cultures (Fig. S6). Consistent with the transcriptional results, the protein level of TNF-alpha released from the microglial cells showed that LPS strongly up-regulated TNF-alpha, but there was no significant change caused by *Trem2* knockdown (Fig. S6). Similar results were seen in microglia from the homozygous *Trem2* R47H KI mice with no substantial change to *Tnf* expression but a decrease in Il1b expression (Fig. S7). We next investigated the effects of this pro-inflammatory treatment on a variety of genes (Fig. S6, S7 and S8). In general, the LPS-dependent effects on gene expression studied here were apparently independent of *Trem2* expression whether knocked-down by siRNA or using the *Trem2* R47H KI microglia. The gene expression profiles were similar whether we normalized to the ubiquitous reference gene *Rps28* or three housekeeping genes (Fig. S8), accordingly all reference genes were stable between genotypes under the conditions tested (Fig. S9 and S10). However, considering the decreased expression of *Trem2* under pro-inflammatory conditions, it is not surprising that further reducing *Trem2* had little additional effect.

### *Trem2* is involved in microglial anti-inflammatory responses with IL-4

Next we studied the effect of acute *Trem2* knockdown on the alternative anti-inflammatory activation of microglia in response to IL-4 treatment. We first found that microglial *Trem2* expression upon IL-4 treatment was maintained at a similar level to that under basal conditions (Fig. 4A and 5A). We next investigated two anti-inflammatory markers *Arg1* and *Tgfb1* (Fig. 4B). Upon IL-4 treatment, *Arg1* expression showed a time-dependent up-regulation, however, this up-regulation was considerably decreased in the *Trem2* knockdown group (two-way ANOVA: interaction between main effects of *Trem2* knockdown and IL-4 treatment time, p=0.05; Sidak’s multiple comparisons: significant difference between the *Trem2* knockdown group and the control at 48 hr post IL-4 treatment, p<0.001). The *Tgfb1* expression also showed an increase with IL-4 stimulation, however, in contrast to *Arg1* the effect of *Trem2* knockdown and IL-4 showed no interaction. Our findings were similar in the microglia from the *Trem2* R47H KI mice, with *Arg1* showing a strong induction by IL-4 treatment and the down-regulation of *Trem2* in the microglia from *Trem2* R47H KI mice significantly preventing this change in a gene dose-dependent manner (Fig. 5B, one-way ANOVA: p=0.002, Sidak’s multiple comparisons: p=0.08 for WT versus heterozygous, and p=0.01 for WT versus homozygous). The other anti-inflammatory marker *Tgfb1* was significantly decreased in homozygous *Trem2* R47H microglia independent of IL-4-treatment, and the fold-change in *Tgfb1* up-regulation in response to treatment was similar for WT and homozygous R47H cells. These findings suggest that separate mechanisms regulate basal levels of some genes like *Tgfb1* (which may be *Trem2*-dependent at least in part), compared to mechanisms that regulate levels following induction (which can be *Trem2*-independent).

**FIGURE 4.**
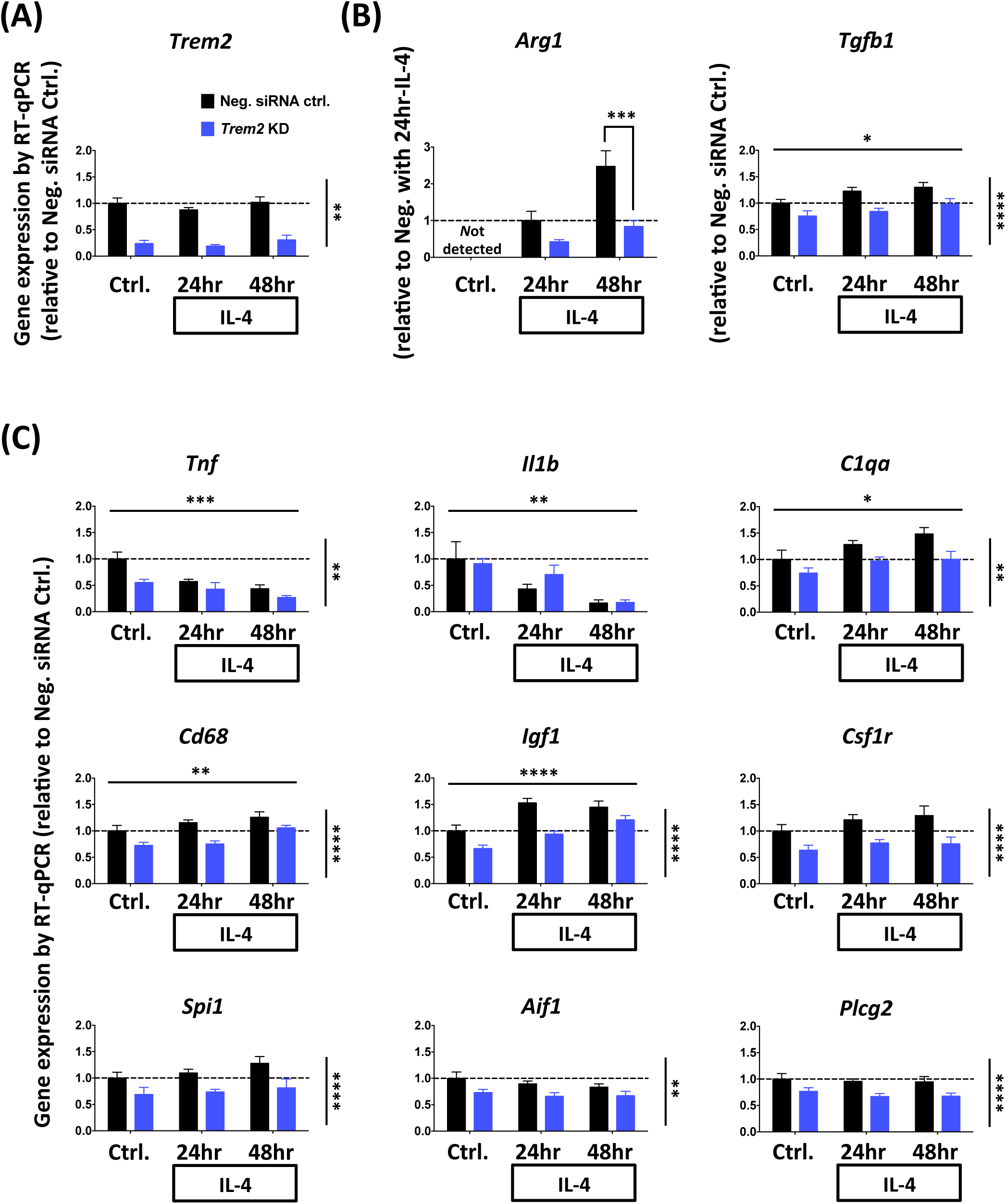
*Trem2* knockdown impairs the IL-4-induced anti-inflammatory response of primary microglia. (A) *Trem2* gene expression was not significantly influenced by IL-4 stimulation. (B) *Arg1* and *Tgfb1* expression, as markers of the anti-inflammatory response, showed significant up-regulation with time after IL-4 application. Particularly, *Trem2* knockdown greatly decreased IL-4-induced *Arg1* expression compared to negative controls. Gene expression levels were normalized to *Rps28* and calculated as fold change relative to the negative control without IL-4 treatment in each individual culture preparation. Two-way ANOVA with significant main effect of IL-4 incubation time and *Trem2* knockdown indicated as horizontal and vertical lines respectively. A significant interaction between IL-4 treatment length and *Trem2* knockdown was seen only in *Arg1* expression (panel B), and so Sidak’s *post hoc* tests were performed to test pairwise significance between the negative siRNA control and *Trem2* knockdown at each time-point. (C) Expression of the pro-inflammatory genes (*Tnf* and *Il1b)* and other microglial genes. Gene expression levels were normalized to *Rps28* and calculated as fold change relative to the negative control without IL-4 treatment in each individual culture preparation. N=7-9 independent experiments. Data shown as mean ± SEM. Two-way ANOVA; significant main effects of IL-4 treatment time and *Trem2*-knockdown indicated by horizontal and vertical lines respectively, no significant interactions were seen between IL-4 treatment and *Trem2* knockdown; * p<0.05, ** p<0.01, *** p<0.001, **** p<0.0001.

**FIGURE 5.**
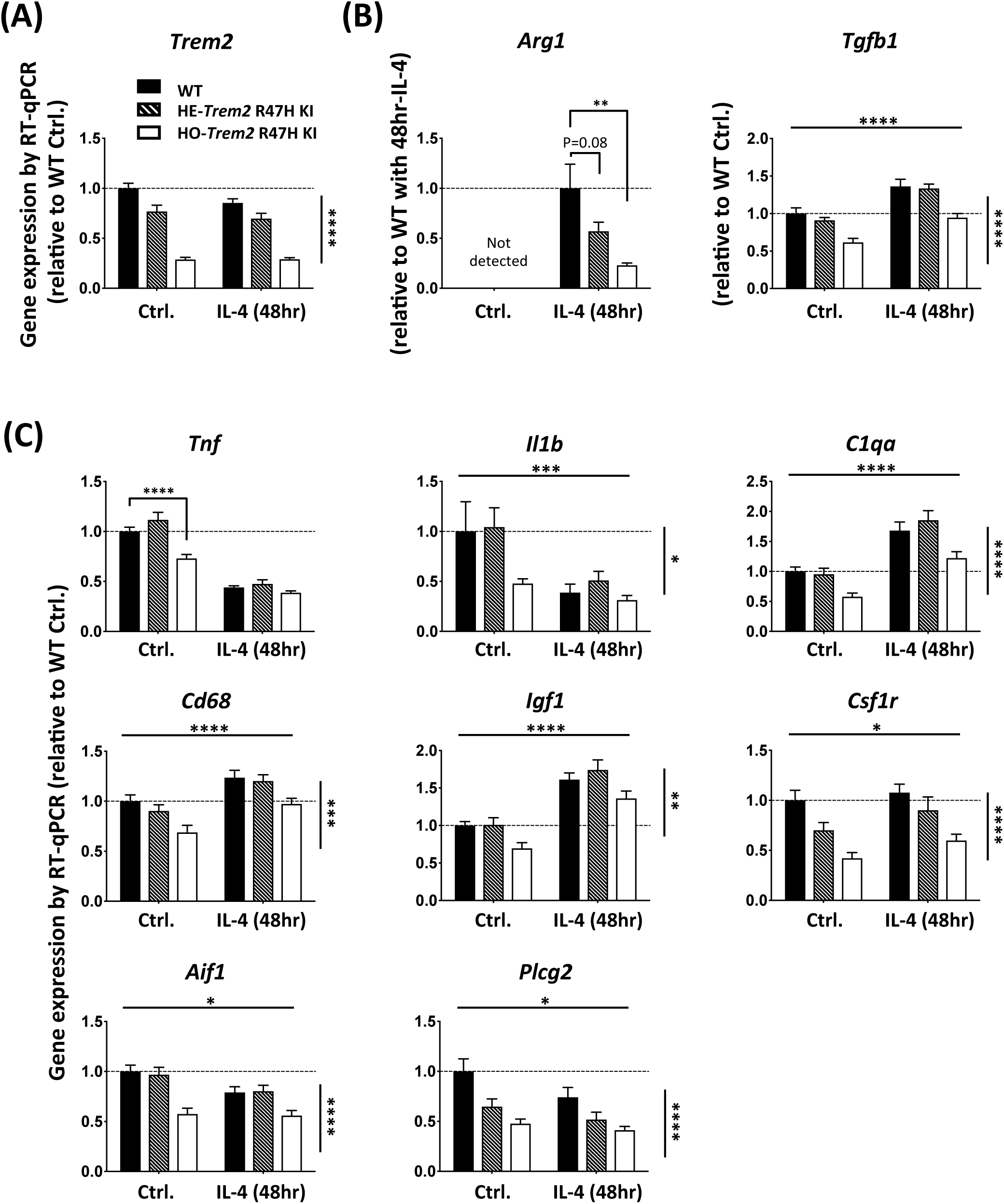
Primary microglia from *Trem2* R47H KI mice show decreased *Trem2* expression, and an impaired IL-4-induced anti-inflammatory response. (A) Expression of *Trem2*. (B) Expression of anti-inflammatory markers, *Arg1* and *Tgf1b*. (C) Expression of other microglial genes. Gene expression was normalized to *Rps28* and calculated as fold change relative to the WT control without IL-4 treatment. N=8-15 mice per genotype. Data shown as mean ± SEM. Two-way ANOVA (for all genes except *Arg1*). Significant main effect of IL-4-treatment and genotype indicated as horizontal and vertical lines respectively, no significant interactions were seen between IL-4-treatment and *Trem2* knockdown. Sidak’s *post hoc* tests were performed to test pairwise significance between the three genotypes within a treatment group. *Arg1* was analyzed by one-way ANOVA (p=0.002) followed by Sidak’s multiple comparisons to test pairwise significance between the three genotypes within a treatment group. * p<0.05, ** p<0.01, *** p<0.001, **** p<0.0001.

We went on to compare the effect of the anti-inflammatory stimulus on the gene expression levels of a variety of genes (Fig. 4C). As expected, the anti-inflammatory IL-4 treatment decreased expression of pro-inflammatory marker genes *Tnf* and *Il1b* (two-way ANOVA: main effect for IL-4 treatment, p<0.001 for *Tnf,* and p<0.01 for *Il1b*). Expression of *C1qa*, *Cd68* and *Igf1* were all found to be significantly up-regulated with IL-4 treatment, which is similar to what we saw in transgenic mice in correlation with amyloid plaque load (43, 44) (www.mouseac.org). Expression of *Csf1r*, *Spi1*, *Aif1* and *Plcg2* were unaffected by IL-4 treatment. Interestingly, the expression of all of these genes was significantly lower in cells with acute *Trem2* knockdown, and was not dependent on IL-4 stimulation (no significant interaction of knockdown versus IL-4). Similarly to the *Trem2* knockdown experiments, genes for which expression was increased in response to IL-4, such as *Igf1, Cd68 and C1qa,* also showed decreased expression in the homozygous *Trem2* R47H microglia, regardless of IL-4-treatment (Fig. 5C; two-way ANOVA: main effect of IL-4-treatment, and main effect of *Trem2* genotype, but no interaction for *Igf1*, *Cd68* and *C1qa*).

Collectively, these data suggest that the effect of TREM2 on the microglial gene expression profile pushes microglia towards an anti-inflammatory phenotype. TREM2 promotes an anti-inflammatory response via two mechanisms: i) an IL-4 independent mechanism supporting the expression of common genes, such as *Igf1*, *Cd68* and *C1qa*, which are induced by IL-4 in control cells and suppressed by *Trem2* reduction, but there is lack of interaction between the effects of *Trem2* and IL-4 on these genes, and so here TREM2 and IL-4 likely act via independent parallel pathways to converge onto similar endpoints; ii) an IL-4-dependent mechanism where the gene expression response to IL-4 depends on TREM2, genes such as *Arg1*, where there is a significant interaction between the effects of *Trem2* and IL-4. Interestingly, the effects of IL-4 stimulation on gene expression mimic the changes seen in association with plaque development in transgenic mice (44), where *Trem2* levels are also strongly increased, whereas the pro-inflammatory effects of LPS have the opposite effect.

### Gene expression profiles regulated by TREM2 and IL-4

To discover novel gene expression profiles regulated by TREM2 and IL-4, we performed RNA-seq using cultured primary microglia from homozygous *Trem2* R47H KI mice versus WT littermates under basal conditions and in response to IL-4 treatment. As a means of validation of the RNA-seq data, the expression profile of all the test and reference genes assayed by RT-qPCR above were similar to the gene expression obtained by RNA-seq under basal conditions and in response to IL-4 (Fig. S10 and S11, compare to Fig. 4 and 5). Differential expression analysis showed decreased expression of *Trem2* in the microglia from *Trem2* R47H KI mice as expected, and also reduced expression of a number of genes including *Abi3* (*ABI family member 3*), *Cd72* (*CD72 antigen*), *Ch25h* (*cholesterol 25-hydroxylase*), *Apoc2* (*apolipoprotein C II*), *Oas1b* (*oligoadenylate synthetase 1b*) and *Neurl1a* (*neuralized E3 ubiquitin protein ligase 1A*) (311 genes with decreased expression, DESeq2 FDR<0.05; Fig. 6A; Table S1). Genes showing increased expression in the *Trem2* R47H KI microglia included *App (amyloid precursor* protein), *Hspa1b* (*heat-shock 70 kDa protein 1B*), and *Ccnd2* (*cyclin D2*) (408 genes with increased expression, DESeq2 FDR<0.05; Fig. 6A; Table S2). The genes differentially expressed in response to reduced *Trem2* expression in the *Trem2* R47H KI microglia showed a significant overlap with a network of genes expressed by amyloid-responsive microglia from bulk RNA-seq analysis of mouse hippocampus, which showed up to a 9.2-fold increase in *Trem2* expression, providing evidence that altering *Trem2* levels impacts expression of similar downstream genes in this method of culturing microglia (Fisher’s Exact test p=3·5e-47) (44). The genes differentially expressed due to reduced *Trem2* expression in the *Trem2* R47H KI microglia were significantly enriched for biological annotations associated with immune cell cytoskeletal processes, actin organization, and cell chemotaxis/migration (Fig. 6B).

**FIGURE 6.**
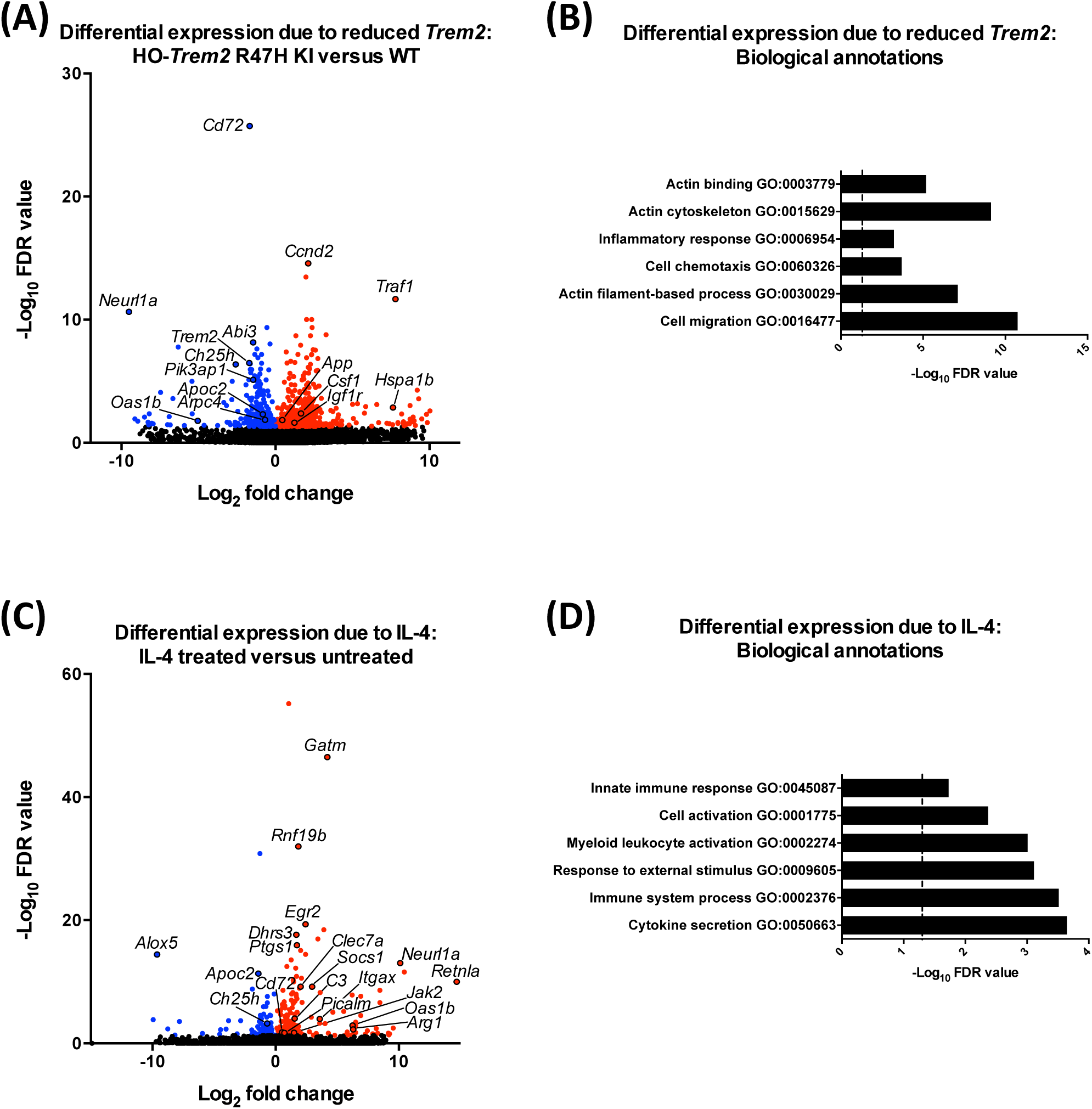
Gene expression profile in primary microglia from *Trem2* R47H KI and WT mice with IL-4 treatment by RNA-seq. (A) Volcano plot showing differentially expressed genes for microglia from HO-*Trem2* R47H KI versus WT mice by DESeq2 (FDR<0.05). (B) Biological annotations associated with differentially expressed genes for microglia from HO-*Trem2* R47H KI versus WT mice (FDR<0.05) with enrichment p-values corrected for multiple testing. Vertical dashed line signifies corrected enrichment p=0.05. (C) Volcano plot showing differentially expressed genes for microglia treated with IL-4 versus untreated by DESeq2 (FDR<0.05). (D) Biological annotations associated with differentially expressed genes for microglia treated with IL-4 versus untreated (FDR<0.05) with enrichment p-values corrected for multiple testing. Vertical dashed line signifies corrected enrichment p=0.05. N=3 mice per genotype per condition.

We next investigated differential expression in the cultured primary microglia in response to IL-4 treatment, and found many genes induced by IL-4, including *Arg1* (described above), *Retnla* (*resistin-like alpha*), *Clec7a* (*C-type lectin domain family 7, member A*), *Itgax*/CD11C (*integrin alpha X*), *Jak2* (*Janus kinase 2*), *Picalm* (*phosphatidylinositol binding clathrin assembly protein*), *Oas1b*, *Neurl1a*, and *Gatm* (*glycine amidinotransferase*) (184 genes with increased expression, DESeq2 FDR<0.05; Fig. 6C; Table S3). We also found a number of genes showed decreased expression in response to IL-4, including *Ch25h*, *Apoc2*, and *Alox5* (*arachidonate 5-lipoxygenase*) (83 genes with decreased expression, DESeq2 FDR<0.05; Fig. 6C; Table S4). As a means of validation, we saw a significant overlap of genes regulated by IL-4 in the cultured primary microglia, compared with genes regulated by IL-4 in microglia in the mouse spinal cord *in vivo* (Fisher’s Exact test p=1·8e-12) (45), confirming that this method of culturing microglia *in vitro* can model a significant component of the *in vivo* response. In addition, the genes responding to IL-4 treatment of the microglia were significantly enriched for biological annotations associated with immune cell activation and cytokine secretion (Fig. 6D).

To investigate further the response of the microglia to reduced *Trem2* expression and IL-4 treatment we performed weighted gene co-expression network analyses (WGCNA) (46), with an optimization for constructing more biologically meaningful co-expression networks (47). We identified a network associated with reduced *Trem2* expression that contained *Trem2*, the adaptor *Tyrobp,* and the microglial transcription factors *Spi1*/PU.1 and *Stat6* (network correlation with *Trem2* genotype, Pearson’s product-moment correlation=-0·73, p=0.007; Fig. 7A). Hub genes are those with the highest connectivity within a co-expression network, and are postulated to drive the response of the entire network. The hub genes in the genetic network associated with *Trem2* status were *Nckap1l* (*NCK associated protein 1 like*), *Cd53* (*leukocyte surface antigen 53*), *Adam8* (*a disintegrin and metallopeptidase domain 8*) and *Fxyd5* (*FXYD domain-containing ion transport regulator 5*). The genetic network containing *Trem2* was significantly enriched for the gene expression module expressed by hippocampal microglial (p=4.70e-7), and also significantly enriched for biological annotations associated with immune cell cytoskeletal processes and actin organization as above (Fig. S12). We then identified a network associated with IL-4 treatment that contained *Arg1*, *Retnla*, *Clec7a*, *Picalm* and *Gatm* (network correlation with IL-4 treatment, Pearson’s product-moment correlation=0.98, p=9e-9; Fig. 7B). The hub genes in the genetic network associated with IL-4 treatment were *Ap2m1* (*adaptor-related protein complex 2, mu 1 subunit*), *Gatm*, *Ptgs1*/COX1 (*prostaglandin-endoperoxide synthase 1*), *Dhrs3* (*dehydrogenase/reductase SDR family member 3*) and *Rnf19b* (*ring finger protein 19B*). The genetic network associated with IL-4 treatment was significantly enriched for biological annotations associated with vesicular and endosomal functions (Fig. S12).

**FIGURE 7.**
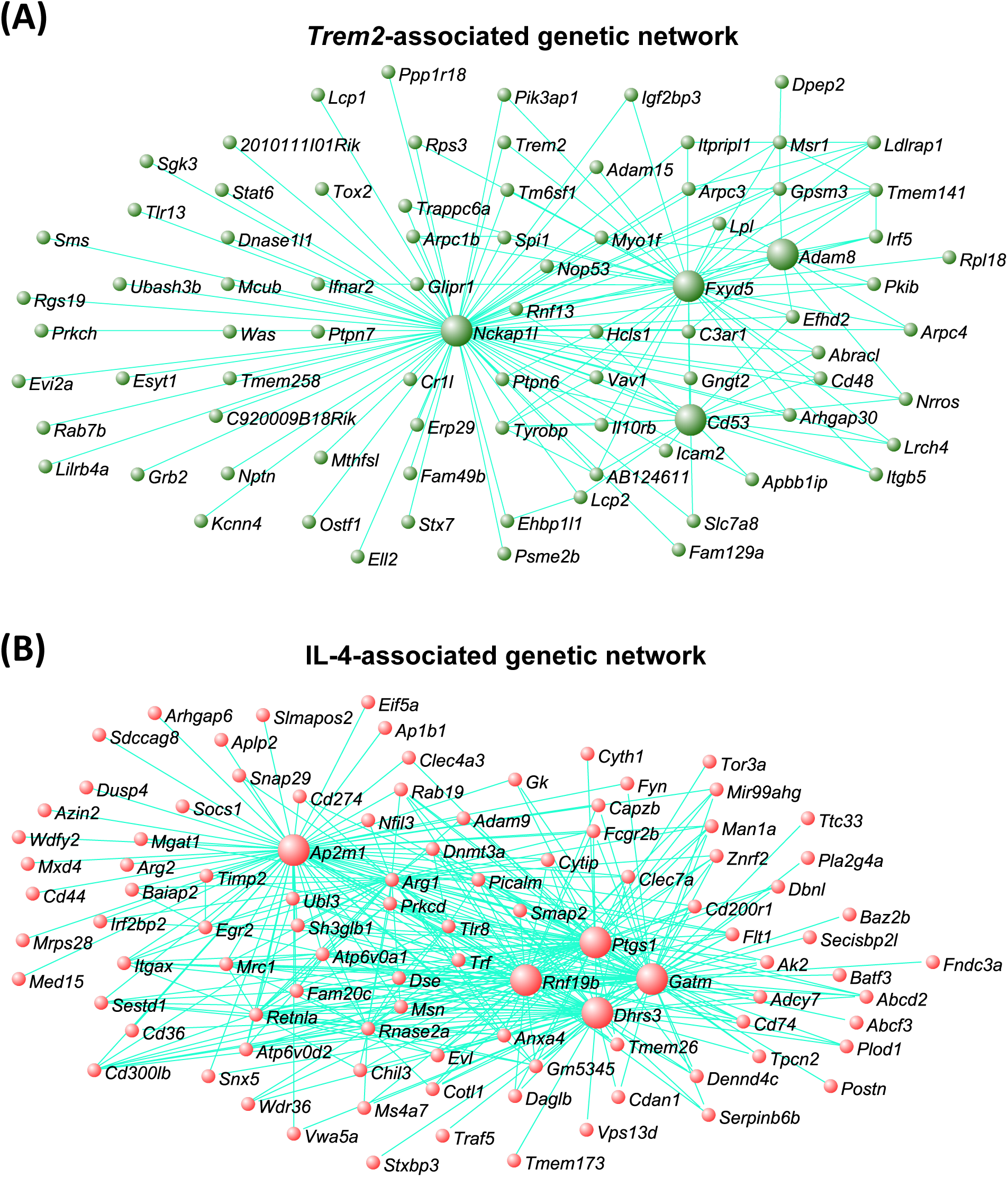
Co-expression genetic networks associated with *Trem2* expression and IL-4-treatment. (A) Co-expression network associated with *Trem2* expression, using RNA-seq derived gene expression from primary microglia of *Trem2* R47H KI and WT mice. Network contains key genes mediating microglial functions, and includes *Trem2*, *Spi1*/PU.1, *Tyrobp* and *Stat6*. Larger green spheres represent ‘hub’ genes, those showing the greatest number of connections to other genes in the network, which are likely to play important roles in driving microglial functions alongside *Trem2*. (B) Co-expression network associated with IL-4-treatment, using RNA-seq derived gene expression from primary microglia treated with IL-4 and untreated. Network includes key genes involved in anti-inflammatory processes, such *Arg1*, *Retnla* and *Chil3*. Larger red spheres represent ‘hub’ genes, which are likely to play important roles in driving the microglial response to IL-4. N=3 mice per genotype per condition.

We next aimed to identify genes that behave similarly to our canonical anti-inflammatory marker *Arg1*, in that their induction or expression change by IL-4 is dependent on *Trem2,* and so we selected the cluster of genes showing the highest connectivity to *Arg1* from the co-expression genetic network associated with IL-4 (Topological overlap measure, TOM>0.4; Fig. 8A; Table S5). The genes part of the signature showing strong connectivity to *Arg1* included *Ap1b1 (adaptor protein complex AP-1, beta 1 subunit*, also known as beta 1 adaptin), *Dusp4* (*dual specificity phosphatase 4*), *Itgax*/CD11C, and *Rnf19b*. To understand how this signature of genes associated with *Arg1* was regulated, we mined for transcription factor recognition elements in the promoters of genes correlating with *Arg1* expression using i-*cis*Target (48). We found an enrichment of a number of recognition elements for different transcription factors, including the canonical microglial transcription factor SPI1/PU.1, and also STAT6, in the promoters of genes behaving like *Arg1* (Fig. 8B), This suggests that TREM2 and IL-4 regulate an overlapping signature of genes that are controlled by transcription factors including STAT6 and SPI1.

**FIGURE 8.**
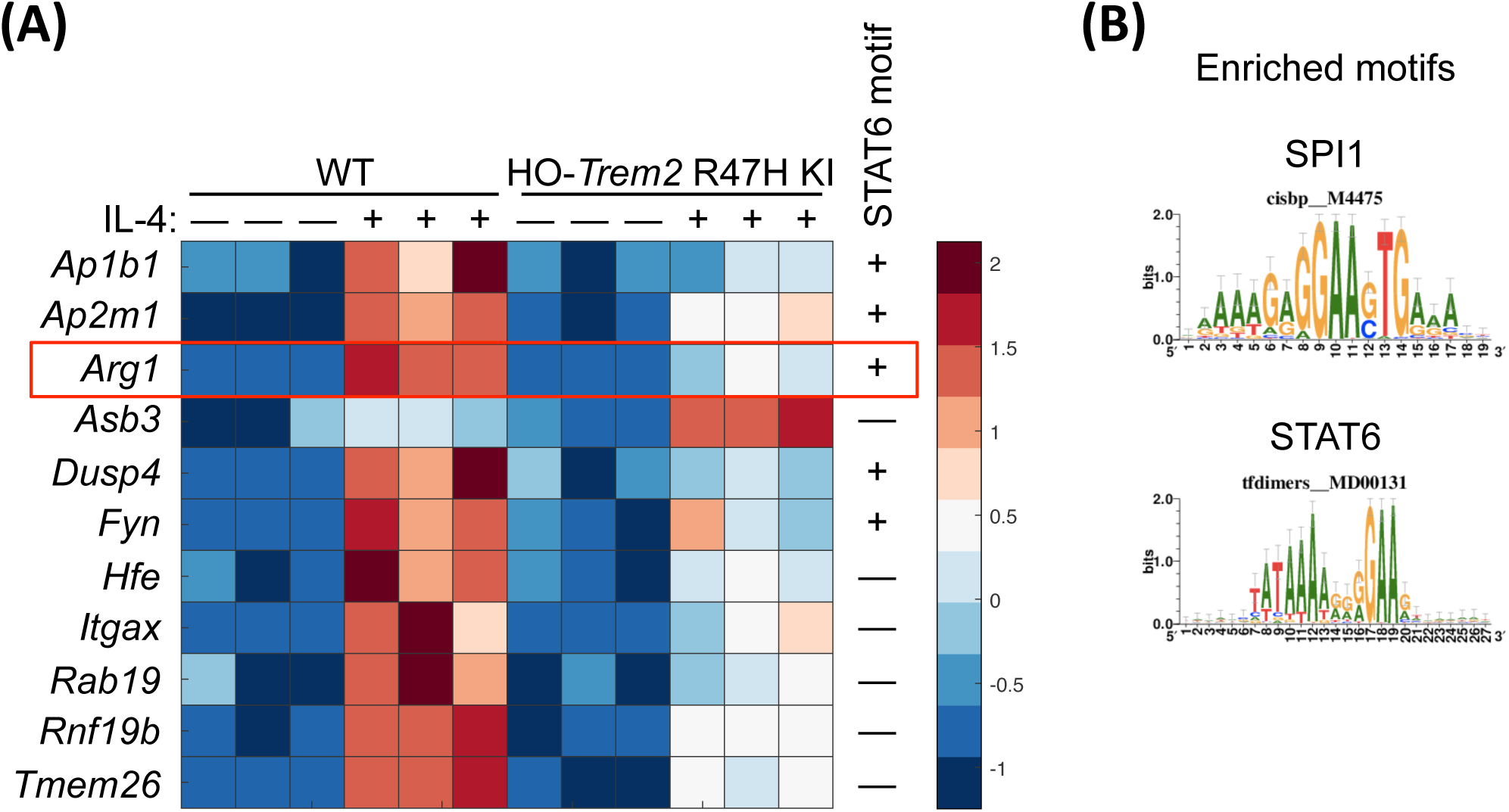
Cluster of genes showing the highest connectivity to *Arg1* from the genetic network associated with IL-4. (A) Heat map showing the genes with the highest connectivity to *Arg1* from the co-expression network associated with IL-4 (TOM>0.4). Selection of genes presented with expression tested by two-way ANOVA, all genes showing a significant main effect of IL-4-treatment and genotype, and a significant interaction. The expression of these genes is represented in primary microglia from HO-*Trem2* R47H KI versus WT mice treated with IL-4 or untreated. Colours represent the z-score of the expression level for each gene (red is high expression and blue is low expression). (B) The promoters of genes that show the highest connectivity to *Arg1* (TOM>0.4) contain an enrichment of consensus binding sites for a number of transcription factors including SPI1/PU.1 and STAT6 determined using i-*cis*Target. Consensus matrices for SPI1/PU.1 and STAT6 are shown. N=3 mice per genotype per condition.

### TREM2 is involved in maintaining normal STAT6 levels in microglia

As up-regulation of genes such as *Arg1* and *Ap1b1* caused by IL-4 stimulation interacted with and was largely suppressed by reduced *Trem2* expression, we next investigated whether there was an intersection between the IL-4 and TREM2 pathways. We found an enrichment of STAT6 transcription factor recognition elements in the promoters of genes behaving like *Arg1* and regulated by IL-4, and previous studies have shown that phosphorylation of STAT6 is a key step downstream in the IL-4 cascade that leads to *Arg1* expression (49–51). We thus performed western blotting to test whether total or phosphorylated levels of STAT6 were changed in primary microglia with acute *Trem2* knockdown with or without IL-4 stimulation (Fig. 9A and 9B). We found a significant IL-4-induced increase in STAT6 levels but, independent of IL-4, knockdown of *Trem2* caused a substantial decrease of the STAT6 total protein levels in the microglia (two-way ANOVA: main effect of IL-4-treatment, p<0.05; main effect of *Trem2* knockdown, p<0.0001; no interaction). IL-4 caused robust phosphorylation of STAT6, with phosphorylated STAT6 not detectable in the absence of IL-4 but clearly measurable 48 hr after IL-4 treatment. *Trem2* knockdown also decreased the total levels of phosphorylated STAT6 but this could be explained by a decrease in the overall levels of STAT6 rather than an effect on phosphorylation *per se*. The expression level of the *Stat6* gene was also decreased with reduced *Trem2* expression (Fig. 9C), and so confirms the decreased levels of the translated protein. These results suggest that TREM2 deficiency leads to a smaller STAT6 pool in microglia, resulting in decreased amount of phosphorylated STAT6 upon IL-4 stimulation. This is consistent with the decrease in IL-4-induced *Arg1* and *Ap1b1* expression caused by reduced STAT6-dependent microglial gene transcription in response to decreased *Trem2* expression.

**FIGURE 9.**
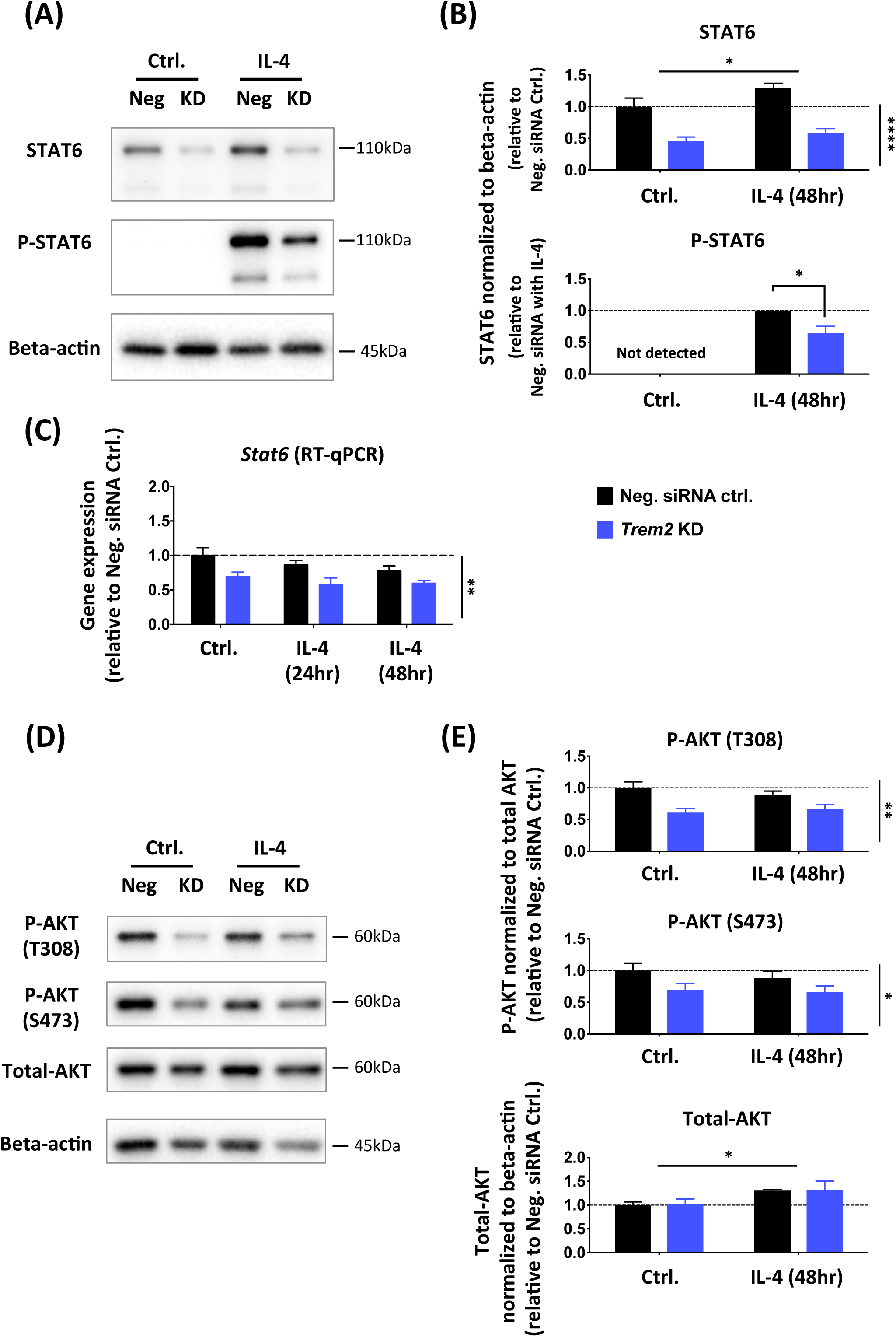
*Trem2* knockdown resulted in significantly decreased total STAT6 levels and AKT phosphorylation in primary microglia *in vitro*. (A-B) Total and phosphorylated STAT6 protein levels in primary microglia treated with IL-4 and *Trem2* siRNA were analyzed with western blot. N=4 independent experiments. (C) *Stat6* gene expression levels in primary microglia, assessed by RT-qPCR (N=3 independent experiments). (D-E) Total and phosphorylated (at Thr308 and Ser473) AKT protein levels analyzed by western blot. N=4 independent experiments. Protein levels were normalized to beta-actin and gene expression was normalized to *Rps28*. Fold change was calculated relative to the negative control without IL-4 treatment in each individual culture preparation. Note that phosphorylated STAT6 was not detectable under control conditions and so under IL-4 conditions the effect of *Trem2* knockdown was calculated as fold change relative to control cells. Data shown as mean ± SEM. With the exception of phosphorylated STAT6, data were analyzed by two-way ANOVA; significant main effects of IL-4 treatment and *Trem2*-knockdown indicated by horizontal and vertical lines respectively, no significant interactions were seen between IL-4 treatment and *Trem2* knockdown. Phosphorylated STAT6 was analyzed by a one-sample *t*-test. * p<0.05, ** p<0.01, *** p<0.001, **** p<0.0001.

TREM2 signaling is also mediated by the PI3K/AKT pathway (40, 52), and so we also investigated the effects of *Trem2* knockdown on AKT signaling in primary microglia (Fig. 9D and 9E). Different from the decrease of total STAT6 protein levels, the total AKT protein level was not significantly affected by *Trem2* knockdown, although there was a slight increase in AKT levels in response to IL-4 treatment. We then tested activation of AKT, and found that AKT phosphorylation at both Thr308 and Ser473 relative to total AKT was significantly reduced in *Trem2* knockdown microglia (two-way ANOVA: main effect of *Trem2* knockdown, p<0.01 for phosphoT308-AKT and p<0.05 for phosphoS473-AKT), and this was independent of IL-4 stimulation (no interaction between knockdown and IL-4). This is not surprising considering that PI3K-AKT signaling is downstream of TREM2, but the unchanged total AKT level with knockdown suggests that the STAT6 change is not due to general changes in the health and metabolic status of microglia in our cultures.

## Discussion

### *Trem2* R47H knock-in mice can largely be considered a model of *in vivo Trem2* down-regulation

Of the many genes that have been revealed in GWAS to increase the risk of AD, variants of *TREM2* have high odds ratios. The most common of these is the R47H mutation. We thus aimed to investigate the effect of this mutation in *Trem2* R47H KI mice. However in the present study we find that the primary phenotype of these mice both in primary microglial cultures and in brain tissue, is a substantial gene-dose-dependent decrease in the expression of *Trem2* compared to WT mice. This effect has been described by Cheng-Hathaway *et al.* (37) and Xiang *et al.* (36) studying the same mouse line used here from the Jackson Laboratory, and an independent *Trem2* R47H KI mouse line. This does not occur in human tissue with this mutation but appears to be a mouse specific effect related to altered splicing. Hence, we can largely consider these mice as a model of *Trem2* down-regulation, validated by our siRNA data in primary microglia. By investigating microglial density in fixed brains from the *Trem2* R47H mice, we saw a decrease in the density of microglia *in vivo* in the hippocampus, and that the remaining microglia had dramatically reduced CD68 protein levels. This raises the question as to whether any changes that we found in *Trem2* R47H microglia are directly due to the mutation itself or downstream of its effect on decreasing *Trem2* expression, and so we studied in parallel the effect of *Trem2* knockdown using siRNA in primary microglia and compared this to the *Trem2* R47H KI microglia phenotype.

### siRNA-induced knockdown of *Trem2* largely mimics the *Trem2* R47H phenotype in non-stimulated conditions

Knockdown of *Trem2* in the primary microglia transfected with siRNA showed similar effects to those seen in microglia from the *Trem2* R47H KI mice. In both preparations *Csf1r* expression was decreased. Pharmacological blockade of CSF1 receptors has been shown to impair microglial survival and can completely remove microglia from the mouse brain (53–55). These findings together with the decreased density of microglia in the hippocampus of *Trem2* R47H mice and increased apoptosis of the microglia cultured from these mice, suggests that the decrease in *Csf1r* expression in the microglia with decreased *Trem2* expression may contribute to the survival of microglia. The knockdown of *Trem2* decreasing survival and altering the metabolism of microglia is consistent with previous studies (35, 38–40, 56, 57). Interestingly no difference in *Csf1r* expression was seen in the remaining microglia in the brain tissue from the young adult KI mice, possibly suggesting different populations of microglia, with loss of those more vulnerable to the effects of *Trem2* knockdown than others, at least in whole tissue. Differences in gene expression profiles of different microglial populations have been recently described using single cell RNA-seq showing some microglial populations are selectively dependent on *Trem2* (58–62).

*Igf1* expression was also decreased when *Trem2* expression was knocked-down or if the *Trem2* R47H variant was carried by the primary microglial cultures. *Igf1* is abundantly expressed by microglia during postnatal development, ageing, or following brain injuries and plays a vital role in promoting neuronal survival (59, 63–66). While this may not affect microglial survival as the IGF1 receptor is found largely on neurons, it could contribute to neurodegeneration, being an important factor in neuronal survival and lifespan across many species (67). Our findings, in line with previous studies (59, 68–71), show that *Igf1* expression is inhibited by pro-inflammatory stimuli and enhanced in anti-inflammatory environments. Thus, the *Igf1* expression decrease caused by *Trem2* deficiency may suggest a microglial phenotype shift towards pro-inflammatory activation and away from non-inflammatory, and a decrease of the pro-survival role of IGF1 on neurons.

### *Trem2* supports an anti-inflammatory programme of gene expression by IL-4 stimulation in part by promoting levels of the transcription factor STAT6

In this study we have further investigated the role of TREM2 and effects on *Trem2* expression in microglia polarized to anti-inflammatory or pro-inflammatory phenotypes. Anti-inflammatory processes are associated with tissue repair, such as those mediated by IL-4, working through microglia and macrophages, and have been shown to provide benefits in models representing the early stages of neurodegenerative conditions, such as AD and amyotrophic lateral sclerosis, and other disorders related to cell death (45, 72, 73). Our results suggest that there is cross-talk between the TREM2 and anti-inflammatory IL-4 signaling pathways, in that TREM2 knockdown decreases levels of the STAT6 transcription factor, which is downstream of IL-4 (74). As the IL-4-induced up-regulation of a number of genes including *Arg1* is critically dependent on phosphorylation of STAT6 (75), this suggests that TREM2 signaling functions in parallel to support IL-4-induced STAT6-gene transcription by supporting overall levels of STAT6. Decreased *Trem2* expression in both primary microglia models resulted in impaired induction of a programme of anti-inflammatory genes induced by IL-4, including genes such as *Arg1*, *Ap1b1* and *Dusp4*.

ARG1 is an important anti-inflammatory marker induced upon IL-4 stimulation. ARG1 in myeloid cells mainly functions as an enzyme to hydrolyze arginine to produce ornithine as part of the urea cycle, which both promotes tissue repair through generation of polyamines and collagens, and also reduces nitric oxide production from inducible nitric oxide synthase through decreasing intracellular arginine availability (76–78). ARG1-positive microglia have been shown to modify Aβ accumulation (79), and IL-33 ameliorated AD-associated phenotypes by promoting anti-inflammatory functions, which included induction of *Arg1* (80). Other genes induced by IL-4 included *Ap1bp1*, *Dusp4* and *Itgax*. AP1B1 is part of the clathrin-associated adaptor protein complex AP-1 and is involved in endocytosis and intracellular trafficking (81). Further work is required to understand the function of AP1B1 in microglia, although work with a microglial cell line suggests that Aβ binding to scavenger receptors induces oxidative stress, which then may impair AP1B1-dependent internalisation of the Aβ (82). DUSP4 is a phosphatase for a number of mitogen-activated protein kinase (MAPK) family members, also known as MAPK phosphatase 2 (MKP-2). DUSP4 has not been studied well in microglia, but *Dusp4* null mice show impaired T cell activation (83). Finally, *Itgax*/CD11C is an integrin or cell surface antigen, and is upregulated in a number of recent RNA-seq studies in response to amyloid plaques (58, 59, 61, 84). ITGAX-positive microglia are a source of *Igf1* expression (59). Our findings in cultured primary microglia suggest an attenuated anti-inflammatory phenotype with reduced expression of *Trem2* (Fig. 10).

**FIGURE 10.**
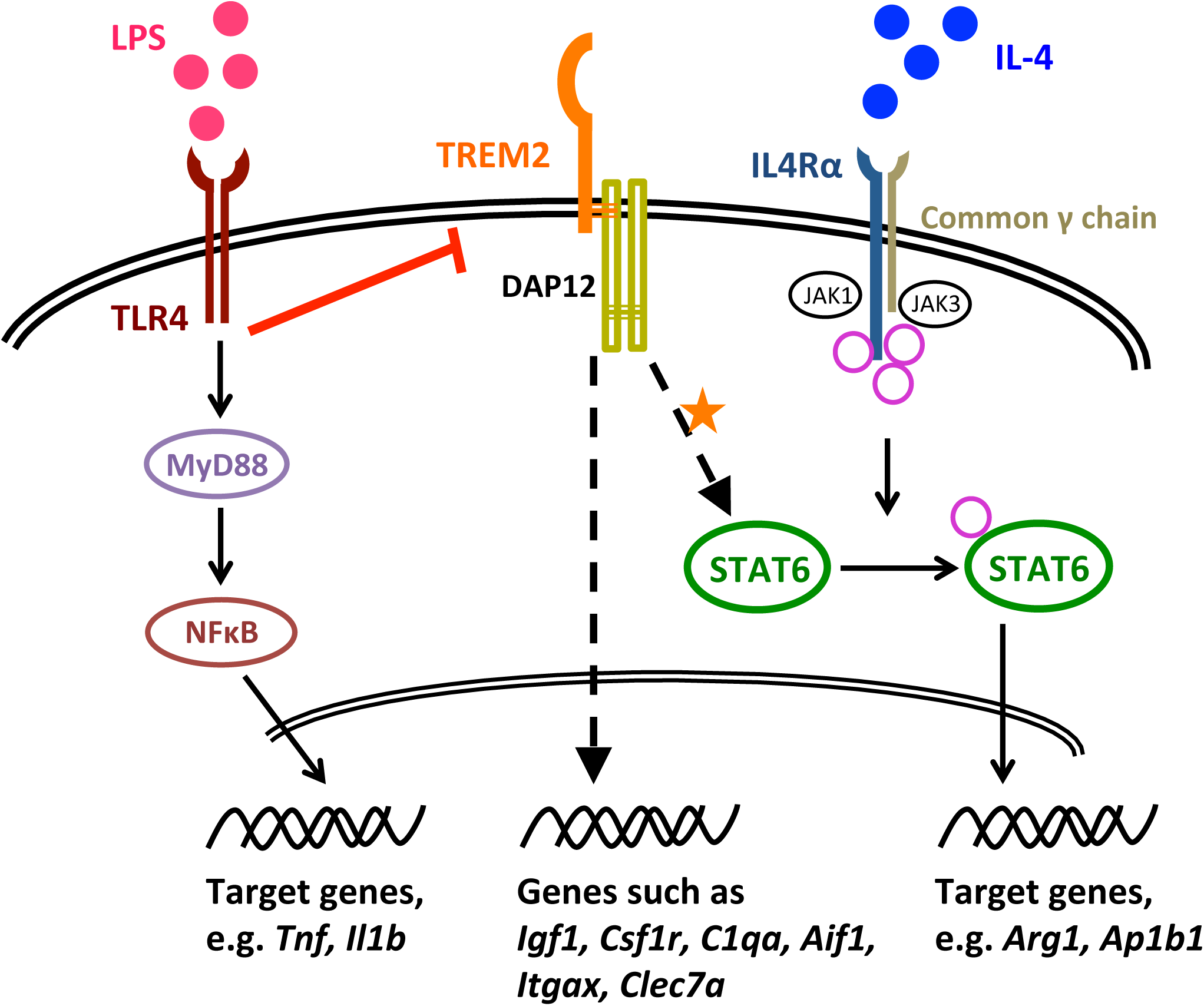
Proposed model of the involvement of TREM2 in the IL-4-induced anti-inflammatory response. LPS induces pro-inflammatory activation of microglia, which is accompanied by down-regulated *Trem2* expression, and so microglia with reduced TREM2 activity generally show normal activation in response to the pro-inflammatory LPS stimulus. Under non-stimulated conditions, a number of microglial genes are down-regulated when TREM2 activity is decreased, such as *Csf1r* expression, which may result in the seen reduced survival of microglia. However, IL-4 induces an anti-inflammatory phenotype of microglia. As our data suggest, TREM2 signaling is involved in maintenance of microglial STAT6 levels, which in the IL-4 pathway is phosphorylated and translocates to the nucleus and functions as a key transcription factor for IL-4-induced gene expression changes, such as for *Arg1* and *Ap1b1*. Thus, *Trem2* deficiency results in decreased STAT6 levels in microglia, which leads to an impairment of IL-4-induced signaling. TREM2 also regulates the levels of a number of genes that are up-regulated by IL-4 (suppressed by LPS), and considered a component of the transcriptional response to AD-associated pathology, such as *Igf1*, *C1qa*, *Itgax* and *Clec7a*.

In hippocampal homogenates from young adult *Trem2* R47H mice, we did not see differences in expression of the genes we studied, including *Stat6* and *Igf1*, despite a slight but significant reduction in the number of microglia. Lack of gene expression changes *in vivo* may be due to a supportive environment from healthy neurons and astrocytes, providing regulating input to the microglia in young mice. Other studies have not revealed obvious phenotypes in young *Trem2* R47H expressing mice, or even *Trem2* knockout mice, unless they are aged or bred to amyloid mouse models (35, 61, 85–87). The effects seen in the primary cultured microglia probably reflect a different mean activation state from microglia in the brain environment, although there remains a significant overlap with microglia expression profiles *in vivo* that are dependent on *Trem2* levels and IL-4 simulation. Thus the pathway interactions described here between *Trem2* expression and other genes, in primary microglial cultures, nevertheless reflects effects that *Trem2* can have under particular conditions *in vivo*. This also applies to the relationship between *Trem2* expression and the IL-4-induction of *Arg1* expression and an anti-inflammatory gene expression programme putatively via STAT6.

### Pro-inflammatory stimuli prevent the effects of TREM2 by inhibiting its expression, similar to the effect seen in microglia from *Trem2* R47H KI mice

With LPS stimulation, primary microglia from WT mice exhibited a robust pro-inflammatory response, as measured by strong up-regulation of *Tnf* and *Il1b* expression (Fig. 10). In contrast to IL-4 stimulation, which had little effect on *Trem2* expression, LPS largely suppressed *Trem2*. In the presence of LPS, decrease in expression of *Trem2* began as early as 3 hr after LPS application and reached more than an 85% decrease by 24 hr. This finding is consistent with previous studies showing that LPS/IFNγ stimulation down-regulated *Trem2* expression via TLR4 receptors in cell lines and primary cultures of microglia/macrophages and also in mouse brains (14, 16, 42, 88, 89). Moreover, Owens and colleagues (42) showed that treatment with pro-inflammatory mediators engaging other Toll-like receptors (TLRs) also suppressed *Trem2* expression in the BV-2 cell line. This suggests that the *Trem2* down-regulation is not specific to LPS-TLR4 signaling but perhaps a common event occurring in microglia induced by various classical pro-inflammatory stimuli. In conditions of *Trem2* knockdown with siRNA, similar LPS-induced up-regulation of *Tnf* and *Il1b* expression resulted. It was, however, interesting to note that, in microglia originating from *Trem2* R47H KI mice with a similar level of *Trem2* down-regulation, there was about a 39% decrease in the activation of *Il1b* compared to WT mice possibly suggesting that the *Trem2* R47H mutation underlies this effect. There are other possible explanations for this difference however, as the down-regulation of *Trem2* in these mice is present throughout life and so these microglia may have a different state when isolated than those in which *Trem2* is knocked down acutely.

Taken together, pro-inflammatory mediators, either locally produced, or infiltrating into the CNS from the periphery following blood-brain barrier breakdown, could induce down-regulation of microglial *Trem2* expression and subsequently induce a microglial state with low TREM2-signaling (Fig. 10). This might reflect a mechanism for microglia to perform specific functional requirements under pro-inflammatory conditions. Although, accumulated pro-inflammation and oxidative stress in the brain, which is often observed in aged brains and aged microglia (90–94), could cause sustained suppression of *Trem2* expression and thus an impairment of TREM2-dependent microglial functions. This might be one of the major mechanisms underlying various neurodegenerative diseases with advanced age by suppression of wild type TREM2 in many sporadic cases, considering that *TREM2* variants associated with loss-of-function are extremely rare in frequency in humans (2).

### Functions of TREM2 in Alzheimer’s disease and the effect of the R47H mutation

Recent important studies have also shown that impaired/absent *Trem2* function attenuates the microglial response to amyloid pathology by regulating microglial number around plaques, regulating microglial activity, and restricting neuritic damage (37, 61, 84–87, 95–99). However anti-inflammatory signaling was not studied. An important issue about microglial activation *in vivo* has to be carefully considered. In contrast to populations of microglia polarized with purified LPS or IL-4 stimuli *in vitro*, microglia *in vivo* probably never receive such simple stimuli to govern their activation state or at least not across the whole population. Instead, they are modulated by mixed cues including various regulatory signals from the local environment (*e.g.* from neurons and astrocytes). There is increasing evidence that reactive microglia in diseased brains demonstrate a mixed phenotype with expression of pro- or anti-inflammatory and homeostatic markers, particularly showing heterogeneity at the single-cell level, rather than completely reproducing the phenotype obtained from homogenous *in vitro* polarization studies (58, 60, 61, 100–103). Indeed, a series of recent studies identified transcriptional signatures in response to AD-associated pathology, neurodegeneration and apoptotic neurons, which were in part dependent on *Trem2* and included *Igf1*, *Csf1, Itgax, Clec7a and Arg1*, at least in mice (58, 59, 61, 84, 86). Therefore, we are cautious in directly applying the *in vitro* phenotype to interpretation of *in vivo* microglial activation. However, we propose that pro- and anti-inflammatory stimuli still represent an important part of the driving factors in the spectrum of microglial activation, and understanding their effects separately may help to differentiate possible mechanisms for moderating function in disease.

Overall, our data shows that decreased expression of *Trem2* has acute and chronic effects on a subset of non-inflammatory functions in microglial cells that overlap with IL-4 anti-inflammatory signaling (Fig. 10). Interestingly, when we tested expression of genes that show substantial changes in transgenic mice associated with AD (43, 44) a number of aspects of gene expression were mimicked by anti-inflammatory IL-4 stimulation, particularly the increased expression of genes associated with the neurodegenerative signature, including *Igf1, Itgax, Clec7a and Arg1* (58, 59, 61, 84, 86). Instead, the pro-inflammatory effects of LPS generally had the opposite effect, leading to almost complete down-regulation of *Trem2* expression. This suggests that the very strong microglial response to plaque deposition that we have previously reported in transgenic mouse models of AD (43, 44), particularly the up to 9-fold increase in *Trem2* expression, preferentially involves engagement of anti-inflammatory pathways that may contribute to the resistance of mice with a heavy amyloid load from developing the full AD pathology of Tau tangles and neurodegeneration, discussed further in Ref (44, 104). Further studies are necessary to investigate whether the IL-4 signaling pathway cross-talk with TREM2 is involved in a protective microglial phenotype in response to amyloid pathology in AD models.

Generally, *Trem2* seems to enhance an anti-inflammatory programme of genes that may protect against disease progression and moreover may increase the number of protective microglia. Pro-inflammatory stimuli inhibit this effect by down-regulating the expression of *Trem2*. Although in this study, the down-regulation of *Trem2* expression in the *Trem2* R47H KI mice which is not reflected in humans with this mutation, means that we can conclude little about the effect of the R47H mutation itself, this is nevertheless a model for loss of function of *Trem2* and validates our findings on the effects of siRNA-induced knockdown. Our findings are thus compatible with increased expression of *TREM2* in Alzheimer’s disease having a protective function slowing disease progression, and hence loss of function due to the presence of the *Trem2* R47H mutation increasing the chance of AD reaching the stage of diagnosis (44, 104–107). Further work into the molecular pathways regulated by TREM2, particularly the cross-talk with anti-inflammatory pathways, may provide important insights for therapeutic approaches.

## Materials and Methods

### Mice

All experiments were performed in agreement with the Animals (Scientific Procedures) Act 1986, with local ethical approval. *Trem2* R47H knock-in (R47H KI) mice were obtained from the Jackson Laboratory (C57BL/6J-*Trem2^em1Adiuj^*/J; stock number: 027918, RRID: IMSR_JAX:027918) (36), and maintained on a C57BL/6J background. All mice were bred and maintained at UCL on a 12 hr light/12 hr dark cycle with *ad libitum* supply of food and water. Genomic DNA was extracted for genotyping using the ‘HotSHOT’ lysis method. Alkaline lysis reagent (25 mM NaOH, 0.2 mM EDTA [pH 12]) was added to tissue samples prior to heating to 95°C for 30 min. The sample was then cooled to 4°C before the addition of neutralization buffer (40 mM Tris-HCl [pH 5]). Mice were genotyped using a quantitative PCR reaction utilizing the following primers and probes (Thermo Fisher Scientific):

Forward primer 5’ ACT TAT GAC GCC TTG AAG CA 3’,
Reverse primer 5’ CTT CTT CAG GAA GGC CAG CA 3’,
Wild type probe 5’ VIC-AGA CGC AAG GCC TGG TG -MGBNFQ 3’,
Mutant probe 5’ 6-FAM-AGA CAC AAA GCA TGG TGT CG -MGBNFQ 3’

TaqMan genotyping master mix was used according to the manufacturer’s instructions.

### Primary microglial culture

Primary microglial cultures were generated from cerebral cortices and hippocampi of 1-3-day-old newborns from WT C57BL/6J or *Trem2* R47H KI mouse strains, following an adapted protocol of Schildge *et al.* (108) and Saura *et al.* (109). For each independent preparation of WT primary microglial culture, brains of 4-6 pups from one litter of neonatal mice were dissected and pooled. For the *Trem2* KI mice, primary cultures from 2-4 pups from the same litter were prepared simultaneously but cultured separately, and genotyping was performed after culture preparation. On average, around 3,000,000∼4,000,000 mixed glial cells from the combined cortex and hippocampus per pup were obtained. Specifically, after decapitation, the brain was quickly extracted and transferred to ice-cold HBSS. The cortices and hippocampi from each brain were then isolated by removing all other brain regions, meninges and blood vessels. Following mechanical and enzymatic (HBSS with 0.25% trypsin) dissociation, cells were centrifuged, collected and seeded at a density of 120,000 cells/cm^2^ in poly-D-lysine coated flasks/wells with a glial culture medium (high glucose DMEM with 10% heat-inactivated fetal bovine serum, 100 U/ml penicillin and 100 µg/ml streptomycin). Cells were cultured at 37°C in humidified 5% CO_2_ and the medium was changed every 3 days. After 19-23 days, once the mixed glial cells achieved confluence, a mild trypsinization method (incubation with DMEM with 0.05% trypsin and 0.2 mM EDTA for 30-45 min) was applied to purify the primary microglia by removing the astrocytic cell layer. Conditioned medium from the mixed glia was collected before trypsinization. Isolated primary microglial cells were collected and seeded in poly-D-lysine coated plates at a density of 40,000-100,000 cells/cm^2^ using the collected conditioned medium, except for the cultures used for ELISA, which were cultured in fresh medium. Pure primary microglia were cultured under the same conditions as the mixed glial culture, and experiments were started at 24 hr after isolation.

### Adult mouse brain tissue extraction

Brains were removed from the skull on ice following decapitation, and cut into two hemispheres. The hippocampus and cortex were dissected from one hemisphere and then snap frozen on dry ice and stored at −80°C until total RNA was isolated by homogenisation of tissue with a polytron. The second hemisphere was fixed in 4% paraformaldehyde overnight at 4°C and then cryoprotected and stored in 30% sucrose in phosphate buffered saline (PBS) with 0.02% sodium azide.

### Hippocampal histology

The fixed hemisphere was serially sectioned at 30 µm transverse to the long axis of the hippocampus using a frozen microtome (Leica, Germany). For assessing microglial density and activity, free floating sections were permeabilized in 0.3% Triton-X in PBS three times, blocked (3% goat serum in 0.3% Triton X-100 in PBS) for 1 hr at room temperature and incubated with primary antibodies at 1:500 dilution in blocking solution (IBA1 rabbit antibody, #234003, Synaptic Systems, RRID: AB_10641962; CD68 rat antibody, #MCA1957, Bio Rad, RRID: AB_322219), overnight at 4°C. Sections were washed, then incubated for 2 hr at room temperature with corresponding Alexa-conjugated secondary antibodies diluted to 1:800 in blocking solution (goat anti-rabbit Alexa 488 and goat anti-rat Alexa 594 antibodies, Jackson ImmunoResearch). Sections were washed, nuclei counterstained with 4’,6-diamidine-2-phenylindole (DAPI), then mounted with Fluoromount-G medium (Southern Biotech). Entire hippocampus was imaged in each section using an EVOS FL Auto microscope (Life Technologies) with a x20 objective, by area defined serial scanning and a motorised stage. For AIF1 and CD68 staining, cells were counted in the CA1: *stratum radiatum* in subfields of size 91,100 µm^2^, *stratum lacunosum-moleculare* (subfield 68,300 µm^2^), *stratum pyramidale* and *stratum oriens* (subfields 76,900 µm^2^). Microglial cells were only counted if their DAPI-stained nucleus could be clearly seen surrounded by AIF1 staining with at least two processes protruding from this, and more than 50% of the cell body being present in the subfield. CD68 positive cells were characterized by the cell body containing greater than 25% fluorescence. At least three serial hippocampal sections approximately 720 µm apart were counted for each mouse. Experimenter was blind to the genotype of the samples during quantification and analysis.

### BV-2 cell culture

BV-2 cells (maximum passage number<20), were cultured in medium containing DMEM with 10% heat-inactivated fetal bovine serum, 2 mM L-glutamine, 100 U/ml penicillin, 100 µg/ml streptomycin, and 1% mycoplasma removal agent (AbD Serotec), and passaged every 2-4 days (avoiding confluence of more than 90%). Prior to experiments, BV-2 cells were collected with Accutase (A6964, Sigma), and seeded on 6-well plates at a density of around 10,000 cells/cm^2^ for testing siRNA sequences.

### siRNA transfection

The following three siRNAs (Ambion) that target mouse *Trem2* were tested (with a non-targeting siRNA as a control, Ambion, Catalog No. 4457287):

siRNA No. 1 sequences: sense 5’-CCCUCUAGAUGACCAAGAUtt-3’, antisense 5’-AUCUUGGUCAUCUAGAGGGtc-3’;
siRNA No. 2 sequences: sense 5’-GCGUUCUCCUGAGCAAGUUtt-3’, antisense 5’-AACUUGCUCAGGAGAACGCag-3’;
siRNA No. 3 sequences: sense 5’-GCACCUCCAGGAAUCAAGAtt-3’, antisense 5’-UCUUGAUUCCUGGAGGUGCtg-3’)

Nuclease-free water was used to make 20 µM siRNA solutions. The 3 *Trem2* siRNAs were tested in BV-2 cells and the siRNA No. 2 and No. 3 were selected for *Trem2* knockdown in primary microglia.

Primary mouse microglia were transfected with either *Trem2* siRNA or the non-targeting negative control siRNA using the transfection reagent Lipofectamine RNAiMax (Invitrogen), following the manufacturer’s instructions. The amount of siRNA to volume of Lipofectamine per well was 1 pmol siRNA to 0.3 µl Lipofectamine. Cells were harvested 72±2 hr after transfection.

### LPS or IL-4 treatment

For the microglial activation studies, primary microglia were treated with 1 µg/ml lipopolysaccharides (LPS; L4391, Sigma) at 3, 6, or 24 hr before harvest, or 20 ng/ml Interleukin-4 (IL-4; 214-14, PeproTech) at 24 or 48 hr before harvest. Time points for the addition of LPS or IL-4 treatment are given in Fig. 2.

### Cell survival

Purified primary microglia were cultured on 24-well plates at a density of around 50,000 cells/cm^2^ (100,000 cells/well), from WT or *Trem2* R47H KI sibling mice. After 72 hr, microglia were washed with PBS (without Ca^2+/^Mg^2+^), and dissociated with trypsin (0.25%). Cells were washed twice with ice cold Annexin V binding buffer containing: 10 mM HEPES/NaOH [pH 7.4], 150 mM NaCl, 5 mM KCl, 5 mM MgCl_2_ and 1.8 mM CaCl_2_. Cells were resuspended in Annexin V-FITC (diluted 1:100, cat no: 560931; BD), for 30 min on ice in the dark. Cells were then stained with 0.83 µg/ml propidium iodide (Sigma), just prior to running on a CytoFLEX S flow cytometer (Beckman Coulter, UK), counting at least 1,000 cells per sample. Excitation at 488 nm and detection within a range of 500-535 nm for Annexin V-FITC, and excitation at 561 nm and detection within 610-620 nm for propidium iodide. Following identification of single microglial cells with the expected size and granularity by forward (FSC-H) versus side (SSC-H) scatter of light, microglia with positive staining for annexin V and/or PI were were plotted and analyzed using CytExpert software. Unstained cells, and cells treated with 50 nM staurosporine for 24 hr were used as negative and positive controls respectively to allow correct gating.

### Phagocytosis assay

The phagocytosis assay using the pHrodo green *E. coli* bioparticles conjugate (P35366, Invitrogen) was adapted from the Invitrogen protocol (30, 41). In brief, purified primary microglia were cultured on 24-well plates at a density of around 50,000 cells/cm^2^ (100,000 cells/well) and transfected with either *Trem2* siRNA or the negative control siRNA. After 72 hr, microglia were treated with 50 µg pHrodo-conjugated *E. coli* per well for 1 hr at 37°C. Cells pre-incubated with 10 µmol cytochalasin-D for 30 min prior to the phagocytosis assay were used as a negative control. Cells were re-suspended with PBS after the assay, and then washed and collected in ice-cold fluorescence-activated cell sorting (FACS) buffer (PBS without Ca^2+/^Mg^2+^, with 0.5% BSA, 0.05% sodium azide and 2 mM EDTA). Collected microglia were assessed (10,000 cells counted per sample) by flow cytometry (FACSCalibur running CellQuest Pro; Becton Dickinson, UK) with excitation at 488 nm and emission within a range of 500-535 nm. Following identification of single microglial cells with the expected size and granularity by forward (FSC-H) versus side (SSC-H) scatter of light, microglia with different amounts of phagocytosed fluorescent *E. coli* and thus with different fluorescent intensities were plotted and analyzed using Flowing software (developed by Perttu Terho, Turku Centre for Biotechnology, Finland; www.flowingsoftware.com).

### Primary microglia lysis for gene expression and protein analysis

For gene expression analysis, primary microglia were washed with ice-cold PBS three times and then lysed in ice-cold QIAzol RNA lysis reagent (Qiagen). Lysed cells were snap frozen on dry ice and stored at −80°C.

To analyze cellular protein, primary microglia were washed with ice-cold PBS three times and then lysed in 2X Laemmli dye. Lysed cells were boiled at 95°C in a heating block for 5 min and stored at −20°C.

For ELISA analyses, supernatants from primary microglial culture were collected, centrifuged to remove cell debris and stored at −20°C.

### RNA purification and cDNA preparation

Prior to RNA purification, the primary microglia samples were homogenized using a 1 ml syringe and a G21 needle, or hippocampal samples were homogenized using a polytron. Total RNA was isolated using miRNAeasy columns (Qiagen) following manufacturer’s instructions. The concentration of RNA was assessed with a NanoDrop Spectrophotometer (Thermo Scientific), with the A_260_/A_280_ ratio typically around 2. Randomly selected RNA samples from different experiments (*Trem2* knockdown and non-targeting siRNA, or from *Trem2* R47H mice and WT littermates; n = 6 per group under control conditions as well as treatment with LPS and IL-4), were tested by capillary electrophoresis using an Experion (Bio Rad) and Tapestation (Agilent) to confirm the quality and concentration of the total RNA (Fig. S9 and S10). Typical RNA Integrity values (RIN) around 9.0 from 10.0 were obtained regardless of siRNA, genotype of mice or treatment type.

The same amount of RNA from each sample was first treated using DNase I (Amplification Grade, Invitrogen) plus RNaseOUT (Invitrogen). The reverse transcription reaction was then performed using the High Capacity cDNA Reverse Transcription Kit with RNase Inhibitor (Applied Biosystems) following the manufacturer’s instructions, in parallel with a negative control lacking the reverse transcriptase.

### Primer design and test

Primers were designed to span at least two exons and tested for specificity against the entire mouse transcriptome using Primer-BLAST (NCBI) and supplied by Eurofins MWG Operon. All primers were tested for specificity by performing a standard PCR reaction and resolving the products on a 3% agarose gel with ethidium bromide, followed by a quantitative RT-qPCR reaction to obtain the linearity range for primers, calculate primer efficiency and test primer specificity using a “melt-curve” analysis. All primer sequences are listed in Table S6.

### RT-qPCR

The cDNA samples were tested in triplicate with a reverse-transcriptase (RT) lacking control in a 96-well plate using the CFX96 system (BioRad), with each 20 µl reaction containing the cDNA dilution, 0.25 µM of forward and reverse primers, and SYBR Green PCR master mix (Bio-Rad). Cycling conditions were: 95°C-3 min, 40 cycles of [95°C-10 sec, 58°C-30 sec and 72°C-30 sec], and then 72°C-5 min. A melt curve was generated by heating from 60 to 90°C with 0.5°C increments and 5 sec dwell time. All RT-qPCR reactions were checked for a single peak with the melt-curve analysis reflecting a single PCR product. Expression levels of the housekeeping genes *Arf1*, *H3f3b* and *Rps28* were shown to be stable between the different experimental samples when higher molecular weight RNA was measured (Fig. S9 and S10). The raw cycle threshold (CT) values of target genes (mean of the triplicates for each sample) were normalized to *Rps28* CT values for each sample using the delta CT method (43), and were similar when normalized to the geometric mean of all three housekeeping genes *Arf1*, *H3f3b* and *Rps28*.

### RNA-seq transcriptome work

The quality and concentration of the total RNA was assessed using capillary electrophoresis of each sample from *Trem2* R47H KI and WT primary microglia cultured under basal conditions and in response to IL-4. The RNA-seq library preparation and sequencing was performed by Eurofins Genomics (Ebersberg, Germany). Libraries were created using commercially available kits according to the manufacturer’s instructions, where first strand cDNA synthesis was poly(A)-primed, and then fragmented (SMART-Seq v4 Ultra Low Input RNA Kit for Sequencing, Takara). After fragmentation the library was prepared with the SMARTer ThruPLEX DNA-Seq Kit (Takara). During PCR amplification 10-12 cycles were used. After PCR amplification, the resulting fragments were purified, pooled, quantified and used for cluster generation. For sequencing, pooled libraries were loaded on the cBot (Illumina) and cluster generation was performed according the manufacturer’s instructions. Paired-end sequencing using 100 bp read length (multiplex 12 samples per lane -28M reads) was performed on a HiSeq2500 (HiSeq Control Software 2.2.58) using HiSeq Flow Cell v4 and TruSeq SBS Kit v4. For processing of raw data RTA version 1.18.64 and bcl2fastq v2.20.0.422 was used to generate FASTQ files. Adaptors and low quality base pairs were removed from FASTQ files using Trim Galore (Babraham Bioinformatics). Transcripts were quantified with Salmon (110), using gene annotation from ENSEMBL GRCm38. Salmon was used because it incorporates GC correction and accounts for fragment positional bias. To obtain gene level quantification from the transcripts, and correct for average transcript length and library size, expressed as transcripts per million (TPM), the tximport R package was used (111). TPM values were log2 transformed, and genes were considered expressed when log2 TPM values displayed a mean >1·5 for a given gene, resulting in a total of 11,119 genes expressed.

Weighted gene co-expression network analysis (WGCNA) was performed (46), with an optimization for constructing more biologically meaningful co-expression networks (47). R code and tutorial available from: https://github.com/juanbot/CoExpNets). Genes with variable expression patterns (coefficient of variation >5% considering genotype and IL-4 treatment) from log2 TPM values were selected for network analyses resulting in 10,463 genes for network analyses. The module of genes with high correlation of the module eigengene with *Trem2* genotype or IL-4 treatment was selected for analysis (*Trem2* genotype module, Pearson’s product-moment correlation=-0·73, p=0.007; IL-4 treatment module, Pearson’s product-moment correlation=0.98, p=9e-9). TOM connectivity values were used to plot the network diagrams (TOM>0·461 for *Trem2*-associated module, and TOM>0.488 for IL-4-associated module, to plot approximately the top 50% genes with the highest connections per module). Hub genes were considered to be the top 5% most connected genes within the plotted module. Genetic networks were visualized from the TOM matrix using VisANT 5.0 (112).

### ELISA of TREM2 and TNF-alpha by primary microglia

The TREM2 levels in the primary microglial supernatants were quantified using MaxiSORP 96 well plates (Nunc) coated with 1 µg/ml of a rat anti-mouse/human TREM2 monoclonal antibody (clone 237920, R&D Systems, RRID: AB_2208679) overnight at 4°C. The plates were washed with PBS/0.1% Tween-20 (PBST), then blocked with 1% BSA in PBST for 45 min at room temperature (RT), followed by 3 washed with PBST. 100 µl of supernatant samples and standards (recombinant mouse TREM2-FC, 1729-T2; R&D Systems; 0-20 ng/ml in BSA-PBST) were then incubated for 2 hr at RT, followed by 3 washes with PBST. For the detection, plates were incubated for 1.5 hr at RT with biotinylated sheep anti-mouse TREM2 antibody (0.1 µg/ml, BAF1729, R&D Systems, RRID: AB_356109) diluted in BSA-PBST. Plates were subsequently washed 4 times with PBST, followed by an incubation with streptavidin-HRP (0.2 µg/ml, Invitrogen) diluted in PBST for 45 min at RT. Plates were washed 3 times with PBST followed by the addition of a chromogenic substrate solution (tetramethylbenzidine, TMB) in the dark. The reaction was terminated with the addition of stop solution (0.16 M H_2_SO_4_) and the absorbance was read at 450 nm (Tecan Genios).

The TNF-alpha levels in the primary microglial supernatants were quantified using Quantikine^®^ Mouse TNF-alpha ELISA kit (R&D Systems, MTA000B), following the manufacturer’s instructions. In general, equal volumes of cell supernatants, together with the provided buffer, were loaded in duplicate to the ELISA microplate and incubated for 2 hr at RT. Following five washes, wells were then incubated with horseradish peroxidase conjugated mouse TNF-alpha antibody for another 2 hr at RT. The microplate was washed again and incubated with a substrate solution prepared with chromogen (TMB) and hydrogen peroxide for 30 min before addition of a stop solution with hydrochloric acid. Then the color intensity was measured at 450 nm using a plate reader (EL800, BioTek) and the TNF-alpha concentrations in microglial supernatants were calculated from the standard curve. To account for the differences in the cell number between different wells of microglia, cell lysates were collected in parallel with the supernatants from the same wells and then tested for levels of a housekeeping protein beta-actin (Ab8227, Abcam, RRID: AB_2305186) by western blot, which was used for normalization of TNF-alpha levels.

### Western blotting

Cellular protein samples from primary microglia were loaded onto 15% polyacrylamide gels and resolved by sodium dodecyl sulfate polyacrylamide gel electrophoresis. Proteins were transferred to a 0.45 µm nitrocellulose membrane (BioRad) by wet electro-transfer overnight.

Membranes were washed in Tris-buffered saline (TBS; 30 mM NaCl and 30 mM Tris [pH 7.4]) for 10 min, and then blocked in TBS with 0.1% Tween-20 (TBST) and 5% non-fat milk for 1 hr at room temperature. Membranes were then probed with primary antibody (STAT6 antibody, 5397S, Cell Signaling Technology, RRID: AB_11220421; p-STAT6 (Y641) antibody, 56554S, Cell Signaling Technology; AKT (pan) antibody, 4691T, Cell Signaling Technology, RRID: AB_915783; p-AKT (T308) antibody, 2965S, Cell Signaling Technology, RRID: AB_2255933; p-AKT (S473) antibody, 4060T, Cell Signaling Technology, RRID: AB_2315049; beta-actin antibody, ab8227, Abcam, RRID: AB_2305186) diluted in blocking buffer overnight at 4°C. Following three washes with TBST, membranes were then incubated with a horseradish peroxidase conjugated secondary antibody that was diluted 1:10,000 in blocking buffer for 1 hr at room temperature. Membranes were finally washed 3 more times, and the horseradish peroxidase signals revealed with enhanced chemiluminescence detection (ECL, BioRad). Image acquisition and densitometric analysis was performed using ImageLab (v5.2, BioRad).

### Immunocytochemistry

Primary microglia plated on poly-D-lysine coated coverslips were fixed with 2% PFA for 15 min at room temperature, followed by five washes with PBS. Prior to immunocytochemical staining, coverslips were washed in PBS for 10 min and blocked for 1 hr at room temperature with 5% goat serum diluted in PBS with 0.125% Triton-X 100. Coverslips were then incubated overnight at 4°C with AIF1/IBA1 antibody (diluted 1:1,000; 019-19741, Wako, RRID: AB_839504) and GFAP antibody (diluted 1:1,000; G3893, Sigma, RRID: AB_477010), in the blocking solution. After primary antibody incubation, coverslips were washed three times with PBS and then incubated for 2 hr at room temperature in the anti-rabbit secondary antibody dilution in blocking solution. After three washes with PBS and staining with DAPI, coverslips were mounted on SuperFrost Plus slides (Fisher) with Fluoromount G (Scientific Biotech).

### Imaging and analysis

Coverslips were imaged using an EVOS^®^ FL Auto Cell Imaging System (Life Technologies) under a 20X objective. Two coverslips per genotype and treatment were used and at least six fields of view (600 µm x 600 µm) per coverslip were imaged. AIF1/IBA1-positive, GFAP-positive cells and DAPI were counted with Adobe PhotoShop CS6 software and the microglial purity was calculated as a percentage of the AIF1/IBA1-positive cell counts versus the DAPI counts in the same field of view.

### Statistics

Data are shown as mean ± SEM. The sample size (N) represents the number of independent cell preparations or mice. All statistics were performed using Prism v7 (Graphpad). For the primary microglial gene expression changes with *Trem2* knockdown versus the non-targeting siRNA control, single gene expression level was normalized as a fold change of the negative control within each independent microglial culture preparation, and then a one-sample Student’s *t*-test with Bonferroni correction was conducted for statistical analysis. For the phagocytosis experiment, a paired Student’s *t*-test was used to compare the *Trem2* knockdown group and the non-targeting siRNA control group (samples from the same microglial culture preparation were paired). For the LPS or IL-4 treatment experiments, to assess the impact of *Trem2* knockdown and LPS/IL-4 treatment on the gene expression, data were normalized to the negative control without LPS/IL-4 treatment within each individual microglial preparation and then analyzed with two-way ANOVA followed by Sidak’s multiple comparison tests when appropriate. The *Trem2* R47H KI expression data in whole hippocampal homogenates were tested by one-way ANOVA with Sidak’s multiple comparisons tests when appropriate. The *Trem2* R47H KI immunohistochemistry data were analyzed using two-way ANOVA with Sidak’s multiple comparisons tests when appropriate. For the primary microglia cultured from the *Trem2* R47H KI mice, data were analyzed with two-way ANOVA separately for LPS or IL-4 treatment using the same control samples, and then Sidak’s multiple comparisons tests were performed when appropriate. Differences were considered significant if p<0.05.

RNA-seq read counts were analyzed for differential expression using the DESeq2 package (113). DESeq2 uses raw read counts, applies normalization, and estimates dispersion. Genes were filtered to remove those with low levels of expression (minimum of 12 counts across all 12 samples). We tested for *Trem2* genotype and IL-4-treatment effects using the likelihood-ratio test, and state differentially expressed genes as FDR<0.05 (113). Biological annotations using Gene Ontology, REACTOME (114), and KEGG (115) databases were identified using gProfileR2 (116), using genes with FDR<0.05 or present in specific co-expression modules, using all detected genes in our experiment as the background. Comparisons between differentially expressed genes, or genes present in co-expression modules, were performed with Fisher’s exact test, after determining common genes that could be detected between experiments.

## Acknowledgements

Thanks to Dr Lion Shahab for statistical advice. WL funded by a UCL PhD Scholarship. DAS, DMC and FAE were funded by Alzheimer’s Research UK (ARUK) [grant numbers: ARUK-SRF2013-7 to DAS, ARUK-PG2019B-018 to FAE, and an ARUK pump priming grant via the UCL network]. FAE and DMC were also funded by the UK Dementia Research Institute (DRI). PG-R was supported by a Clinical Research fellowship from Alzheimer’s Research UK. AM was supported by the Biotechnology and Biological Sciences Research Council [grant number BB/M009513/1]. TMP was supported by Eisai, working with the Eisai:UCL Therapeutic Innovation Group (TIG) and was also supported by funding to JMP and JH from the Innovative Medicines Initiative 2 Joint Undertaking under grant agreement 115976. The Joint Undertaking receives support from the European Union’s Horizon 2020 research and innovation programme and the European Federation of Pharmaceutical Industries and Associations (EFPIA). JH is also supported by the Dolby Foundation. DAS and JH are members of the UK DRI, which receives its funding from the DRI Ltd, funded by the UK Medical Research Council, Alzheimer’s Society and ARUK.

## References

1. Huang, K.L., Marcora, E., Pimenova, A.A., Di Narzo, A.F., Kapoor, M., Jin, S.C., Harari, O., Bertelsen, S., Fairfax, B.P., Czajkowski, J. et al. (2017) A common haplotype lowers PU.1 expression in myeloid cells and delays onset of Alzheimer’s disease. Nat Neurosci, 20, 1052–1061.

2. Guerreiro, R., Wojtas, A., Bras, J., Carrasquillo, M., Rogaeva, E., Majounie, E., Cruchaga, C., Sassi, C., Kauwe, J.S., Younkin, S. et al. (2013) TREM2 variants in Alzheimer’s disease. N Engl J Med, 368, 117–127.

3. Malik, M., Simpson, J.F., Parikh, I., Wilfred, B.R., Fardo, D.W., Nelson, P.T. and Estus, S. (2013) CD33 Alzheimer’s risk-altering polymorphism, CD33 expression, and exon 2 splicing. J Neurosci, 33, 13320–13325.

4. Jonsson, T., Stefansson, H., Steinberg, S., Jonsdottir, I., Jonsson, P.V., Snaedal, J., Bjornsson, S., Huttenlocher, J., Levey, A.I., Lah, J.J. et al. (2013) Variant of TREM2 associated with the risk of Alzheimer’s disease. N Engl J Med, 368, 107–116.

5. Griciuc, A., Serrano-Pozo, A., Parrado, A.R., Lesinski, A.N., Asselin, C.N., Mullin, K., Hooli, B., Choi, S.H., Hyman, B.T. and Tanzi, R.E. (2013) Alzheimer’s disease risk gene CD33 inhibits microglial uptake of amyloid beta. Neuron, 78, 631–643.

6. Naj, A.C., Jun, G., Beecham, G.W., Wang, L.S., Vardarajan, B.N., Buros, J., Gallins, P.J., Buxbaum, J.D., Jarvik, G.P., Crane, P.K. et al. (2011) Common variants at MS4A4/MS4A6E, CD2AP, CD33 and EPHA1 are associated with late-onset Alzheimer’s disease. Nat Genet, 43, 436–441.

7. Hollingworth, P., Harold, D., Sims, R., Gerrish, A., Lambert, J.C., Carrasquillo, M.M., Abraham, R., Hamshere, M.L., Pahwa, J.S., Moskvina, V. et al. (2011) Common variants at ABCA7, MS4A6A/MS4A4E, EPHA1, CD33 and CD2AP are associated with Alzheimer’s disease. Nat Genet, 43, 429–435.

8. Sims, R., van der Lee, S.J., Naj, A.C., Bellenguez, C., Badarinarayan, N., Jakobsdottir, J., Kunkle, B.W., Boland, A., Raybould, R., Bis, J.C. et al. (2017) Rare coding variants in PLCG2, ABI3, and TREM2 implicate microglial-mediated innate immunity in Alzheimer’s disease. Nat Genet, 49, 1373–1384.

9. Bouchon, A., Dietrich, J. and Colonna, M. (2000) Cutting edge: inflammatory responses can be triggered by TREM-1, a novel receptor expressed on neutrophils and monocytes. J Immunol, 164, 4991–4995.

10. Bouchon, A., Hernandez-Munain, C., Cella, M. and Colonna, M. (2001) A DAP12-mediated pathway regulates expression of CC chemokine receptor 7 and maturation of human dendritic cells. J Exp Med, 194, 1111–1122.

11. Daws, M.R., Lanier, L.L., Seaman, W.E. and Ryan, J.C. (2001) Cloning and characterization of a novel mouse myeloid DAP12-associated receptor family. Eur J Immunol, 31, 783–791.

12. Humphrey, M.B., Daws, M.R., Spusta, S.C., Niemi, E.C., Torchia, J.A., Lanier, L.L., Seaman, W.E. and Nakamura, M.C. (2006) TREM2, a DAP12-associated receptor, regulates osteoclast differentiation and function. J Bone Miner Res, 21, 237–245.

13. Paloneva, J., Mandelin, J., Kiialainen, A., Bohling, T., Prudlo, J., Hakola, P., Haltia, M., Konttinen, Y.T. and Peltonen, L. (2003) DAP12/TREM2 deficiency results in impaired osteoclast differentiation and osteoporotic features. J Exp Med, 198, 669–675.

14. Turnbull, I.R., Gilfillan, S., Cella, M., Aoshi, T., Miller, M., Piccio, L., Hernandez, M. and Colonna, M. (2006) Cutting edge: TREM-2 attenuates macrophage activation. J Immunol, 177, 3520–3524.

15. Paloneva, J., Manninen, T., Christman, G., Hovanes, K., Mandelin, J., Adolfsson, R., Bianchin, M., Bird, T., Miranda, R., Salmaggi, A. et al. (2002) Mutations in two genes encoding different subunits of a receptor signaling complex result in an identical disease phenotype. Am J Hum Genet, 71, 656–662.

16. Schmid, C.D., Sautkulis, L.N., Danielson, P.E., Cooper, J., Hasel, K.W., Hilbush, B.S., Sutcliffe, J.G. and Carson, M.J. (2002) Heterogeneous expression of the triggering receptor expressed on myeloid cells-2 on adult murine microglia. J Neurochem, 83, 1309–1320.

17. Takahashi, K., Rochford, C.D. and Neumann, H. (2005) Clearance of apoptotic neurons without inflammation by microglial triggering receptor expressed on myeloid cells-2. J Exp Med, 201, 647–657.

18. Wang, Y., Ulland, T.K., Ulrich, J.D., Song, W., Tzaferis, J.A., Hole, J.T., Yuan, P., Mahan, T.E., Shi, Y., Gilfillan, S. et al. (2016) TREM2-mediated early microglial response limits diffusion and toxicity of amyloid plaques. J Exp Med, 213, 667–675.

19. Benitez, B.A., Cruchaga, C. and United States-Spain Parkinson’s Disease Research, G. (2013) TREM2 and neurodegenerative disease. N Engl J Med, 369, 1567–1568.

20. Jin, S.C., Benitez, B.A., Karch, C.M., Cooper, B., Skorupa, T., Carrell, D., Norton, J.B., Hsu, S., Harari, O., Cai, Y. et al. (2014) Coding variants in TREM2 increase risk for Alzheimer’s disease. Hum Mol Genet, 23, 5838–5846.

21. Ruiz, A., Dols-Icardo, O., Bullido, M.J., Pastor, P., Rodriguez-Rodriguez, E., Lopez de Munain, A., de Pancorbo, M.M., Perez-Tur, J., Alvarez, V., Antonell, A., et al. (2014) Assessing the role of the TREM2 p.R47H variant as a risk factor for Alzheimer’s disease and frontotemporal dementia. Neurobiol Aging, 35, 444 e441-444.

22. Murray, C.E., King, A., Troakes, C., Hodges, A. and Lashley, T. (2019) APOE epsilon4 is also required in TREM2 R47H variant carriers for Alzheimer’s disease to develop. Neuropathology and applied neurobiology, 45, 183–186.

23. Cady, J., Koval, E.D., Benitez, B.A., Zaidman, C., Jockel-Balsarotti, J., Allred, P., Baloh, R.H., Ravits, J., Simpson, E., Appel, S.H. et al. (2014) TREM2 variant p.R47H as a risk factor for sporadic amyotrophic lateral sclerosis. JAMA Neurol, 71, 449–453.

24. Rayaprolu, S., Mullen, B., Baker, M., Lynch, T., Finger, E., Seeley, W.W., Hatanpaa, K.J., Lomen-Hoerth, C., Kertesz, A., Bigio, E.H. et al. (2013) TREM2 in neurodegeneration: evidence for association of the p.R47H variant with frontotemporal dementia and Parkinson’s disease. Mol Neurodegener, 8, 19.

25. Jiang, T., Tan, L., Chen, Q., Tan, M.S., Zhou, J.S., Zhu, X.C., Lu, H., Wang, H.F., Zhang, Y.D. and Yu, J.T. (2016) A rare coding variant in TREM2 increases risk for Alzheimer’s disease in Han Chinese. Neurobiol Aging, 42, 217 e211–213.

26. Guerreiro, R., Bilgic, B., Guven, G., Bras, J., Rohrer, J., Lohmann, E., Hanagasi, H., Gurvit, H. and Emre, M. (2013) Novel compound heterozygous mutation in TREM2 found in a Turkish frontotemporal dementia-like family. Neurobiol Aging, 34, 2890 e2891–2895.

27. Guerreiro, R.J., Lohmann, E., Bras, J.M., Gibbs, J.R., Rohrer, J.D., Gurunlian, N., Dursun, B., Bilgic, B., Hanagasi, H., Gurvit, H. et al. (2013) Using exome sequencing to reveal mutations in TREM2 presenting as a frontotemporal dementia-like syndrome without bone involvement. JAMA Neurol, 70, 78–84.

28. Gao, X., Dong, Y., Liu, Z. and Niu, B. (2013) Silencing of triggering receptor expressed on myeloid cells-2 enhances the inflammatory responses of alveolar macrophages to lipopolysaccharide. Mol Med Rep, 7, 921–926.

29. Zhong, L., Chen, X.F., Zhang, Z.L., Wang, Z., Shi, X.Z., Xu, K., Zhang, Y.W., Xu, H. and Bu, G. (2015) DAP12 Stabilizes the C-terminal Fragment of the Triggering Receptor Expressed on Myeloid Cells-2 (TREM2) and Protects against LPS-induced Pro-inflammatory Response. J Biol Chem, 290, 15866–15877.

30. Kleinberger, G., Yamanishi, Y., Suarez-Calvet, M., Czirr, E., Lohmann, E., Cuyvers, E., Struyfs, H., Pettkus, N., Wenninger-Weinzierl, A., Mazaheri, F. et al. (2014) TREM2 mutations implicated in neurodegeneration impair cell surface transport and phagocytosis. Sci Transl Med, 6, 243ra286.

31. Xiang, X., Werner, G., Bohrmann, B., Liesz, A., Mazaheri, F., Capell, A., Feederle, R., Knuesel, I., Kleinberger, G. and Haass, C. (2016) TREM2 deficiency reduces the efficacy of immunotherapeutic amyloid clearance. EMBO Mol Med, 8, 992–1004.

32. Atagi, Y., Liu, C.C., Painter, M.M., Chen, X.F., Verbeeck, C., Zheng, H., Li, X., Rademakers, R., Kang, S.S., Xu, H. et al. (2015) Apolipoprotein E Is a Ligand for Triggering Receptor Expressed on Myeloid Cells 2 (TREM2). J Biol Chem, 290, 26043–26050.

33. Yeh, F.L., Wang, Y., Tom, I., Gonzalez, L.C. and Sheng, M. (2016) TREM2 Binds to Apolipoproteins, Including APOE and CLU/APOJ, and Thereby Facilitates Uptake of Amyloid-Beta by Microglia. Neuron, 91, 328–340.

34. Zhao, Y., Wu, X., Li, X., Jiang, L.L., Gui, X., Liu, Y., Sun, Y., Zhu, B., Pina-Crespo, J.C., Zhang, M. et al. (2018) TREM2 Is a Receptor for beta-Amyloid that Mediates Microglial Function. Neuron, 97, 1023–1031 e1027.

35. Nugent, A.A., Lin, K., van Lengerich, B., Lianoglou, S., Przybyla, L., Davis, S.S., Llapashtica, C., Wang, J., Kim, D.J., Xia, D. et al. (2020) TREM2 Regulates Microglial Cholesterol Metabolism upon Chronic Phagocytic Challenge. Neuron, 105, 837–854 e839.

36. Xiang, X., Piers, T.M., Wefers, B., Zhu, K., Mallach, A., Brunner, B., Kleinberger, G., Song, W., Colonna, M., Herms, J. et al. (2018) The Trem2 R47H Alzheimer’s risk variant impairs splicing and reduces Trem2 mRNA and protein in mice but not in humans. Mol Neurodegener, 13, 49.

37. Cheng-Hathaway, P.J., Reed-Geaghan, E.G., Jay, T.R., Casali, B.T., Bemiller, S.M., Puntambekar, S.S., von Saucken, V.E., Williams, R.Y., Karlo, J.C., Moutinho, M. et al. (2018) The Trem2 R47H variant confers loss-of-function-like phenotypes in Alzheimer’s disease. Mol Neurodegener, 13, 29.

38. Wang, Y., Cella, M., Mallinson, K., Ulrich, J.D., Young, K.L., Robinette, M.L., Gilfillan, S., Krishnan, G.M., Sudhakar, S., Zinselmeyer, B.H. et al. (2015) TREM2 lipid sensing sustains the microglial response in an Alzheimer’s disease model. Cell, 160, 1061–1071.

39. Ulland, T.K., Song, W.M., Huang, S.C., Ulrich, J.D., Sergushichev, A., Beatty, W.L., Loboda, A.A., Zhou, Y., Cairns, N.J., Kambal, A. et al. (2017) TREM2 Maintains Microglial Metabolic Fitness in Alzheimer’s Disease. Cell, 170, 649–663 e613.

40. Zheng, H., Jia, L., Liu, C.C., Rong, Z., Zhong, L., Yang, L., Chen, X.F., Fryer, J.D., Wang, X., Zhang, Y.W. et al. (2017) TREM2 Promotes Microglial Survival by Activating Wnt/beta-Catenin Pathway. J Neurosci, 37, 1772–1784.

41. Garcia-Reitboeck, P., Phillips, A., Piers, T.M., Villegas-Llerena, C., Butler, M.G., Mallach, A., Rodrigues, C., Arber, C.E., Heslegrave, A., Zetterberg, H. et al. (2018) Human induced pluripotent stem cell-derived microglia-like cells harboring TREM2 missense mutations show specific deficits in phagocytosis. Cell Rep, 24, 2300–2311.

42. Owens, R., Grabert, K., Davies, C.L., Alfieri, A., Antel, J.P., Healy, L.M. and McColl, B.W. (2017) Divergent Neuroinflammatory Regulation of Microglial TREM Expression and Involvement of NF-kappaB. Front Cell Neurosci, 11, 56.

43. Matarin, M., Salih, D.A., Yasvoina, M., Cummings, D.M., Guelfi, S., Liu, W., Nahaboo Solim, M.A., Moens, T.G., Paublete, R.M., Ali, S.S. et al. (2015) A genome-wide gene-expression analysis and database in transgenic mice during development of amyloid or tau pathology. Cell Rep, 10, 633–644.

44. Salih, D.A., Bayram, S., Guelfi, S., Reynolds, R.H., Shoai, M., Ryten, M., Brenton, J.W., Zhang, D., Matarin, M., Botia, J.A. et al. (2019) Genetic variability in response to amyloid beta deposition influences Alzheimer’s disease risk. Brain Commun, 1, fcz022.

45. Rossi, C., Cusimano, M., Zambito, M., Finardi, A., Capotondo, A., Garcia-Manteiga, J.M., Comi, G., Furlan, R., Martino, G. and Muzio, L. (2018) Interleukin 4 modulates microglia homeostasis and attenuates the early slowly progressive phase of amyotrophic lateral sclerosis. Cell Death Dis, 9, 250.

46. Langfelder, P. and Horvath, S. (2008) WGCNA: an R package for weighted correlation network analysis. BMC Bioinformatics, 9, 559.

47. Botia, J.A., Vandrovcova, J., Forabosco, P., Guelfi, S., D’Sa, K., United Kingdom Brain Expression, C., Hardy, J., Lewis, C.M., Ryten, M. and Weale, M.E. (2017) An additional k-means clustering step improves the biological features of WGCNA gene co-expression networks. BMC Syst Biol, 11, 47.

48. Imrichova, H., Hulselmans, G., Atak, Z.K., Potier, D. and Aerts, S. (2015) i-cisTarget 2015 update: generalized cis-regulatory enrichment analysis in human, mouse and fly. Nucleic Acids Res, 43, W57–64.

49. Luzina, I.G., Keegan, A.D., Heller, N.M., Rook, G.A., Shea-Donohue, T. and Atamas, S.P. (2012) Regulation of inflammation by interleukin-4: a review of “alternatives”. J Leukoc Biol, 92, 753–764.

50. Nelms, K., Keegan, A.D., Zamorano, J., Ryan, J.J. and Paul, W.E. (1999) The IL-4 receptor: signaling mechanisms and biologic functions. Annu Rev Immunol, 17, 701–738.

51. Gordon, S. and Martinez, F.O. (2010) Alternative activation of macrophages: mechanism and functions. Immunity, 32, 593–604.

52. Paradowska-Gorycka, A. and Jurkowska, M. (2013) Structure, expression pattern and biological activity of molecular complex TREM-2/DAP12. Hum Immunol, 74, 730–737.

53. Elmore, M.R., Najafi, A.R., Koike, M.A., Dagher, N.N., Spangenberg, E.E., Rice, R.A., Kitazawa, M., Matusow, B., Nguyen, H., West, B.L. et al. (2014) Colony-stimulating factor 1 receptor signaling is necessary for microglia viability, unmasking a microglia progenitor cell in the adult brain. Neuron, 82, 380–397.

54. Dagher, N.N., Najafi, A.R., Kayala, K.M., Elmore, M.R., White, T.E., Medeiros, R., West, B.L. and Green, K.N. (2015) Colony-stimulating factor 1 receptor inhibition prevents microglial plaque association and improves cognition in 3xTg-AD mice. J Neuroinflammation, 12, 139.

55. Olmos-Alonso, A., Schetters, S.T., Sri, S., Askew, K., Mancuso, R., Vargas-Caballero, M., Holscher, C., Perry, V.H. and Gomez-Nicola, D. (2016) Pharmacological targeting of CSF1R inhibits microglial proliferation and prevents the progression of Alzheimer’s-like pathology. Brain, 139(**Pt3**), 891–907.

56. Wu, K., Byers, D.E., Jin, X., Agapov, E., Alexander-Brett, J., Patel, A.C., Cella, M., Gilfilan, S., Colonna, M., Kober, D.L. et al. (2015) TREM-2 promotes macrophage survival and lung disease after respiratory viral infection. J Exp Med, 212, 681–697.

57. Piers, T.M., Cosker, K., Mallach, A., Johnson, G.T., Guerreiro, R., Hardy, J. and Pocock, J.M. (2020) A locked immunometabolic switch underlies TREM2 R47H loss of function in human iPSC-derived microglia. FASEB J, 34, 2436–2450.

58. Keren-Shaul, H., Spinrad, A., Weiner, A., Matcovitch-Natan, O., Dvir-Szternfeld, R., Ulland, T.K., David, E., Baruch, K., Lara-Astaiso, D., Toth, B. et al. (2017) A Unique Microglia Type Associated with Restricting Development of Alzheimer’s Disease. Cell, 169, 1276–1290 e1217.

59. Butovsky, O. and Weiner, H.L. (2018) Microglial signatures and their role in health and disease. Nat Rev Neurosci, 19, 622–635.

60. Sala Frigerio, C., Wolfs, L., Fattorelli, N., Thrupp, N., Voytyuk, I., Schmidt, I., Mancuso, R., Chen, W.T., Woodbury, M.E., Srivastava, G. et al. (2019) The Major Risk Factors for Alzheimer’s Disease: Age, Sex, and Genes Modulate the Microglia Response to Abeta Plaques. Cell Rep, 27, 1293–1306 e1296.

61. Zhou, Y., Song, W.M., Andhey, P.S., Swain, A., Levy, T., Miller, K.R., Poliani, P.L., Cominelli, M., Grover, S., Gilfillan, S. et al. (2020) Human and mouse single-nucleus transcriptomics reveal TREM2-dependent and TREM2-independent cellular responses in Alzheimer’s disease. Nat Med, 26, 131–142.

62. Hammond, T.R., Dufort, C., Dissing-Olesen, L., Giera, S., Young, A., Wysoker, A., Walker, A.J., Gergits, F., Segel, M., Nemesh, J. et al. (2019) Single-Cell RNA Sequencing of Microglia throughout the Mouse Lifespan and in the Injured Brain Reveals Complex Cell-State Changes. Immunity, 50, 253–271 e256.

63. Ueno, M., Fujita, Y., Tanaka, T., Nakamura, Y., Kikuta, J., Ishii, M. and Yamashita, T. (2013) Layer V cortical neurons require microglial support for survival during postnatal development. Nat Neurosci, 16, 543–551.

64. Fernandez, A.M. and Torres-Aleman, I. (2012) The many faces of insulin-like peptide signalling in the brain. Nat Rev Neurosci, 13, 225–239.

65. Kohman, R.A., DeYoung, E.K., Bhattacharya, T.K., Peterson, L.N. and Rhodes, J.S. (2012) Wheel running attenuates microglia proliferation and increases expression of a proneurogenic phenotype in the hippocampus of aged mice. Brain Behav Immun, 26, 803–810.

66. Thored, P., Heldmann, U., Gomes-Leal, W., Gisler, R., Darsalia, V., Taneera, J., Nygren, J.M., Jacobsen, S.E., Ekdahl, C.T., Kokaia, Z. et al. (2009) Long-term accumulation of microglia with proneurogenic phenotype concomitant with persistent neurogenesis in adult subventricular zone after stroke. Glia, 57, 835–849.

67. Broughton, S. and Partridge, L. (2009) Insulin/IGF-like signalling, the central nervous system and aging. Biochem J, 418, 1–12.

68. Arkins, S., Rebeiz, N., Brunke-Reese, D.L., Biragyn, A. and Kelley, K.W. (1995) Interferon-gamma inhibits macrophage insulin-like growth factor-I synthesis at the transcriptional level. Mol Endocrinol, 9, 350–360.

69. Suh, H.S., Zhao, M.L., Derico, L., Choi, N. and Lee, S.C. (2013) Insulin-like growth factor 1 and 2 (IGF1, IGF2) expression in human microglia: differential regulation by inflammatory mediators. J Neuroinflammation, 10, 37.

70. Wynes, M.W. and Riches, D.W. (2003) Induction of macrophage insulin-like growth factor-I expression by the Th2 cytokines IL-4 and IL-13. J Immunol, 171, 3550–3559.

71. Barrett, J.P., Minogue, A.M., Falvey, A. and Lynch, M.A. (2015) Involvement of IGF-1 and Akt in M1/M2 activation state in bone marrow-derived macrophages. Exp Cell Res, 335, 258–268.

72. Bosurgi, L., Cao, Y.G., Cabeza-Cabrerizo, M., Tucci, A., Hughes, L.D., Kong, Y., Weinstein, J.S., Licona-Limon, P., Schmid, E.T., Pelorosso, F. et al. (2017) Macrophage function in tissue repair and remodeling requires IL-4 or IL-13 with apoptotic cells. Science, 356, 1072–1076.

73. Kiyota, T., Okuyama, S., Swan, R.J., Jacobsen, M.T., Gendelman, H.E. and Ikezu, T. (2010) CNS expression of anti-inflammatory cytokine interleukin-4 attenuates Alzheimer’s disease-like pathogenesis in APP+PS1 bigenic mice. FASEB J, 24, 3093–3102.

74. Ohmori, Y. and Hamilton, T.A. (2000) Interleukin-4/STAT6 represses STAT1 and NF-kappa B-dependent transcription through distinct mechanisms. J Biol Chem, 275, 38095–38103.

75. Gray, M.J., Poljakovic, M., Kepka-Lenhart, D. and Morris, S.M., Jr. (2005) Induction of arginase I transcription by IL-4 requires a composite DNA response element for STAT6 and C/EBPbeta. Gene, 353, 98–106.

76. Yang, Z. and Ming, X.F. (2014) Functions of arginase isoforms in macrophage inflammatory responses: impact on cardiovascular diseases and metabolic disorders. Front Immunol, 5, 533.

77. Mills, C.D., Kincaid, K., Alt, J.M., Heilman, M.J. and Hill, A.M. (2000) M-1/M-2 macrophages and the Th1/Th2 paradigm. J Immunol, 164, 6166–6173.

78. Modolell, M., Corraliza, I.M., Link, F., Soler, G. and Eichmann, K. (1995) Reciprocal regulation of the nitric oxide synthase/arginase balance in mouse bone marrow-derived macrophages by TH1 and TH2 cytokines. Eur J Immunol, 25, 1101–1104.

79. Cherry, J.D., Olschowka, J.A. and O’Banion, M.K. (2015) Arginase 1+ microglia reduce Abeta plaque deposition during IL-1beta-dependent neuroinflammation. J Neuroinflammation, 12, 203.

80. Fu, A.K., Hung, K.W., Yuen, M.Y., Zhou, X., Mak, D.S., Chan, I.C., Cheung, T.H., Zhang, B., Fu, W.Y., Liew, F.Y. et al. (2016) IL-33 ameliorates Alzheimer’s disease-like pathology and cognitive decline. Proceedings of the National Academy of Sciences of the United States of America, 113, E2705–2713.

81. Li, Q., Cheng, Z., Zhou, L., Darmanis, S., Neff, N.F., Okamoto, J., Gulati, G., Bennett, M.L., Sun, L.O., Clarke, L.E. et al. (2019) Developmental Heterogeneity of Microglia and Brain Myeloid Cells Revealed by Deep Single-Cell RNA Sequencing. Neuron, 101, 207–223 e210.

82. Manzano-Leon, N., Delgado-Coello, B., Guaderrama-Diaz, M. and Mas-Oliva, J. (2006) Beta-adaptin: key molecule for microglial scavenger receptor function under oxidative stress. Biochemical and biophysical research communications, 351, 588–594.

83. Barbour, M., Plevin, R. and Jiang, H.R. (2016) MAP kinase phosphatase 2 deficient mice develop attenuated experimental autoimmune encephalomyelitis through regulating dendritic cells and T cells. Sci Rep, 6, 38999.

84. Krasemann, S., Madore, C., Cialic, R., Baufeld, C., Calcagno, N., El Fatimy, R., Beckers, L., O’Loughlin, E., Xu, Y., Fanek, Z. et al. (2017) The TREM2-APOE Pathway Drives the Transcriptional Phenotype of Dysfunctional Microglia in Neurodegenerative Diseases. Immunity, 47, 566–581 e569.

85. Song, W.M., Joshita, S., Zhou, Y., Ulland, T.K., Gilfillan, S. and Colonna, M. (2018) Humanized TREM2 mice reveal microglia-intrinsic and -extrinsic effects of R47H polymorphism. J Exp Med, 215, 745–760.

86. Griciuc, A., Patel, S., Federico, A.N., Choi, S.H., Innes, B.J., Oram, M.K., Cereghetti, G., McGinty, D., Anselmo, A., Sadreyev, R.I. et al. (2019) TREM2 Acts Downstream of CD33 in Modulating Microglial Pathology in Alzheimer’s Disease. Neuron, 103, 820–835 e827.

87. Linnartz-Gerlach, B., Bodea, L.G., Klaus, C., Ginolhac, A., Halder, R., Sinkkonen, L., Walter, J., Colonna, M. and Neumann, H. (2019) TREM2 triggers microglial density and age-related neuronal loss. Glia, 67, 539–550.

88. Zheng, H., Liu, C.C., Atagi, Y., Chen, X.F., Jia, L., Yang, L., He, W., Zhang, X., Kang, S.S., Rosenberry, T.L. et al. (2016) Opposing roles of the triggering receptor expressed on myeloid cells 2 and triggering receptor expressed on myeloid cells-like transcript 2 in microglia activation. Neurobiol Aging, 42, 132–141.

89. Zhou, J., Yu, W., Zhang, M., Tian, X., Li, Y. and Lu, Y. (2019) Imbalance of Microglial TLR4/TREM2 in LPS-Treated APP/PS1 Transgenic Mice: A Potential Link Between Alzheimer’s Disease and Systemic Inflammation. Neurochem Res, 44, 1138–1151.

90. Prolla, T.A. (2002) DNA microarray analysis of the aging brain. Chem Senses, 27, 299–306.

91. Godbout, J.P., Chen, J., Abraham, J., Richwine, A.F., Berg, B.M., Kelley, K.W. and Johnson, R.W. (2005) Exaggerated neuroinflammation and sickness behavior in aged mice following activation of the peripheral innate immune system. FASEB J, 19, 1329–1331.

92. Lee, C.K., Weindruch, R. and Prolla, T.A. (2000) Gene-expression profile of the ageing brain in mice. Nat Genet, 25, 294–297.

93. Verbitsky, M., Yonan, A.L., Malleret, G., Kandel, E.R., Gilliam, T.C. and Pavlidis, P. (2004) Altered hippocampal transcript profile accompanies an age-related spatial memory deficit in mice. Learn Mem, 11, 253–260.

94. Sierra, A., Gottfried-Blackmore, A.C., McEwen, B.S. and Bulloch, K. (2007) Microglia derived from aging mice exhibit an altered inflammatory profile. Glia, 55, 412–424.

95. Leyns, C.E.G., Gratuze, M., Narasimhan, S., Jain, N., Koscal, L.J., Jiang, H., Manis, M., Colonna, M., Lee, V.M.Y., Ulrich, J.D. et al. (2019) TREM2 function impedes tau seeding in neuritic plaques. Nat Neurosci, 22, 1217–1222.

96. Prokop, S., Miller, K.R., Labra, S.R., Pitkin, R.M., Hoxha, K., Narasimhan, S., Changolkar, L., Rosenbloom, A., Lee, V.M. and Trojanowski, J.Q. (2019) Impact of TREM2 risk variants on brain region-specific immune activation and plaque microenvironment in Alzheimer’s disease patient brain samples. Acta neuropathologica, 138, 613–630.

97. Sayed, F.A., Telpoukhovskaia, M., Kodama, L., Li, Y., Zhou, Y., Le, D., Hauduc, A., Ludwig, C., Gao, F., Clelland, C. et al. (2018) Differential effects of partial and complete loss of TREM2 on microglial injury response and tauopathy. Proc Natl Acad Sci U S A, 115, 10172–10177.

98. Meilandt, W.J., Ngu, H., Gogineni, A., Lalehzadeh, G., Lee, S.H., Srinivasan, K., Imperio, J., Wu, T., Weber, M., Kruse, A.J. et al. (2020) Trem2 Deletion Reduces Late-Stage Amyloid Plaque Accumulation, Elevates the Abeta42:Abeta40 Ratio, and Exacerbates Axonal Dystrophy and Dendritic Spine Loss in the PS2APP Alzheimer’s Mouse Model. J Neurosci, 40, 1956–1974.

99. Parhizkar, S., Arzberger, T., Brendel, M., Kleinberger, G., Deussing, M., Focke, C., Nuscher, B., Xiong, M., Ghasemigharagoz, A., Katzmarski, N. et al. (2019) Loss of TREM2 function increases amyloid seeding but reduces plaque-associated ApoE. Nat Neurosci, 22, 191–204.

100. Kim, C.C., Nakamura, M.C. and Hsieh, C.L. (2016) Brain trauma elicits non-canonical macrophage activation states. J Neuroinflammation, 13, 117.

101. Morganti, J.M., Riparip, L.K. and Rosi, S. (2016) Call Off the Dog(ma): M1/M2 Polarization Is Concurrent following Traumatic Brain Injury. PLoS One, 11, e0148001.

102. Chiu, I.M., Morimoto, E.T., Goodarzi, H., Liao, J.T., O’Keeffe, S., Phatnani, H.P., Muratet, M., Carroll, M.C., Levy, S., Tavazoie, S. et al. (2013) A neurodegeneration-specific gene-expression signature of acutely isolated microglia from an amyotrophic lateral sclerosis mouse model. Cell Rep, 4, 385–401.

103. Holtman, I.R., Raj, D.D., Miller, J.A., Schaafsma, W., Yin, Z., Brouwer, N., Wes, P.D., Moller, T., Orre, M., Kamphuis, W. et al. (2015) Induction of a common microglia gene expression signature by aging and neurodegenerative conditions: a co-expression meta-analysis. Acta Neuropathol Commun, 3, 31.

104. Edwards, F.A. (2019) A Unifying Hypothesis for Alzheimer’s Disease: From Plaques to Neurodegeneration. Trends Neurosci, 42, 310–322.

105. Zhong, L., Xu, Y., Zhuo, R., Wang, T., Wang, K., Huang, R., Wang, D., Gao, Y., Zhu, Y., Sheng, X. et al. (2019) Soluble TREM2 ameliorates pathological phenotypes by modulating microglial functions in an Alzheimer’s disease model. Nat Commun, 10, 1365.

106. Schlepckow, K., Monroe, K.M., Kleinberger, G., Cantuti-Castelvetri, L., Parhizkar, S., Xia, D., Willem, M., Werner, G., Pettkus, N., Brunner, B. et al. (2020) Enhancing protective microglial activities with a dual function TREM2 antibody to the stalk region. EMBO Mol Med, 12, e11227.

107. Lee, C.Y.D., Daggett, A., Gu, X., Jiang, L.L., Langfelder, P., Li, X., Wang, N., Zhao, Y., Park, C.S., Cooper, Y. et al. (2018) Elevated TREM2 Gene Dosage Reprograms Microglia Responsivity and Ameliorates Pathological Phenotypes in Alzheimer’s Disease Models. Neuron, 97, 1032–1048 e1035.

108. Schildge, S., Bohrer, C., Beck, K. and Schachtrup, C. (2013) Isolation and culture of mouse cortical astrocytes. J Vis Exp, 71, e50079.

109. Saura, J., Tusell, J.M. and Serratosa, J. (2003) High-yield isolation of murine microglia by mild trypsinization. Glia, 44, 183–189.

110. Patro, R., Duggal, G., Love, M.I., Irizarry, R.A. and Kingsford, C. (2017) Salmon provides fast and bias-aware quantification of transcript expression. Nat Methods, 14, 417–419.

111. Soneson, C., Love, M.I. and Robinson, M.D. (2015) Differential analyses for RNA-seq: transcript-level estimates improve gene-level inferences. F1000Res, 4, 1521.

112. Granger, B.R., Chang, Y.C., Wang, Y., DeLisi, C., Segre, D. and Hu, Z. (2016) Visualization of Metabolic Interaction Networks in Microbial Communities Using VisANT 5.0. PLoS Comput Biol, 12, e1004875.

113. Love, M.I., Huber, W. and Anders, S. (2014) Moderated estimation of fold change and dispersion for RNA-seq data with DESeq2. Genome biology, 15, 550.

114. Fabregat, A., Jupe, S., Matthews, L., Sidiropoulos, K., Gillespie, M., Garapati, P., Haw, R., Jassal, B., Korninger, F., May, B. et al. (2018) The Reactome Pathway Knowledgebase. Nucleic Acids Res, 46, D649–D655.

115. Kanehisa, M., Sato, Y., Kawashima, M., Furumichi, M. and Tanabe, M. (2016) KEGG as a reference resource for gene and protein annotation. Nucleic Acids Res, 44, D457–462.

116. Raudvere, U., Kolberg, L., Kuzmin, I., Arak, T., Adler, P., Peterson, H. and Vilo, J. (2019) g:Profiler: a web server for functional enrichment analysis and conversions of gene lists (2019 update). Nucleic Acids Res, 47, W191–W198.

