## Supplementary material for "*Trem2* promotes anti-inflammatory responses in microglia and is suppressed under pro-inflammatory conditions"

**Supplementary material Table S1. Genes showing decreased expression for microglia from HO-*Trem2* R47H KI versus WT mice by DESeq2 (FDR<0.05).**

**Supplementary material Table S2. Genes showing increased expression for microglia from HO-*Trem2* R47H KI versus WT mice by DESeq2 (FDR<0.05).**

**Supplementary material Table S3. Genes showing increased expression for microglia treated with IL-4 versus untreated by DESeq2 (FDR<0.05).**

**Supplementary material Table S4. Genes showing decreased expression for microglia treated with IL-4 versus untreated by DESeq2 (FDR<0.05).**

**Supplementary material Table S5. Cluster of genes showing the highest connectivity to *Arg1* from the co-expression genetic network associated with IL-4 (TOM>0.4).**

**Supplementary material Table S6. PCR primer sequences employed in this study.**

**Supplementary material Figure S1. Primary microglial culture purity is not affected by LPS or IL-4 treatment.** (A) Primary microglia were cultured from WT and HO-*Trem2* R47H KI mice, and treated with IL-4 for 48 hr, or LPS for 24 hr, according to the schedule given in Fig. 2. Representative images showing AIF1/IBA1 (green) and GFAP (red) double immuno-staining of microglia with DAPI. Note: GFAP-positive cells were rare, and some cells had nuclei that were condensed and homogenously stained with DAPI that were negative for both AIF1/IBA1 and GFAP, suggesting dying/dead cells. (B) The microglial purity was quantified as proportion of AIF1/IBA1-positive cells relative to the total number of DAPI nuclei. N=2 independent experiments. Data shown as mean ± SEM. Two-way ANOVA showed no significant main effects of genotype or treatment.

**Supplementary material Figure S2. Primary microglia cultured from HO-*Trem2* R47H KI mice show increased levels of apoptosis compared to WT cells.** (A) Representative images from flow cytometry of microglia cultured for 72 hr and stained with Annexin V-FITC and propidium iodide. Microglia were initially sorted using forward- and side-scatter to identify single cells of the standard size and granularity. Left panel: representative experiment from untreated microglia cultured in standard conditions. Middle panel: representative experiment from microglia treated with 50 nM staurosporine for 24 hr as a positive control to increase levels of apoptosis. Right panel: representative experiment from microglia cultured in standard conditions and not stained with Annexin V-FITC or propidium iodide to show specificity of signal. (B) Increased apoptotic microglia from HO-*Trem2* R47H KI mice. N=6-10 independent cell preparations. Data shown as mean ± SEM. Two-way ANOVA showed significant main effects of genotype (p=0.018) and cell status (p<0.0001) with a significant interaction (p<0.01) for cells cultured in standard conditions. Thus, Sidak's *post hoc* tests were performed to test pairwise significance between WT and HO-*Trem2* R47H KI, * p<0.05, *** p<0.001. Staurosporine treated cells acted a positive control and showed increased microglial apoptosis, two-way ANOVA showed only a significant main effect of cell status (p<0.0001).

**Supplementary material Figure S3. Trial and validation of *Trem2* knockdown using siRNA.** (A) Comparison of *Trem2* knockdown in the presence of 3 siRNA sequences versus non-targeting siRNA in BV-2 cell cultures (N=3 independent experiments). The siRNA No. 2 was chosen for the following *in vitro* acute *Trem2*-knockdown experiments due to its high knockdown levels. (B) *Trem2* siRNA efficiency tested with RT-qPCR at 72 hr after transfection in primary microglia. N=13 independent cell preparations. (C) Two independent siRNA sequences against *Trem2* gave similar results in primary microglia, the primary siRNA was No. 2, and the alternative siRNA sequence was No. 3 (from panel A). Both siRNAs produced stable *Trem2* knockdown compared to the negative control when analyzed at 72 hr after transfection. Expression of candidate genes were analyzed via RT-qPCR. Expression levels were first normalized to *Rps28*, and then the relative fold change in the knockdown was calculated versus the negative control from the same batch of cell preparation (shown with a dotted line at 1). N=3 independent experiments. Data shown as mean ± SEM. One-way ANOVA showed a significant main effect of siRNA treatment for expression of all genes tested, except for *C1qa* and *Csf1r*, where strong trends towards down-regulation were seen, indicated as horizontal lines. Sidak's *post hoc* tests were used to test pairwise significance between the different siRNA treatments, marked above individual groups. * p<0.05, ** p<0.01, *** p<0.001, **** p<0.0001

**Supplementary material Figure S4. *Trem2* knockdown impaired phagocytosis in primary microglia.** Phagocytosis assay was performed using pHrodo-conjugated *E. coli* and analyzed by FACS. As a negative control, phagocytosis was inhibited with 10 mM cytochalasin D. Left and middle panels: a representative experiment. Left panel: forward (FSC-H) versus side (SSC-H) scatter of light to allow identification of single microglial cells of the expected size and granularity (shown within the blue gate). Middle panel: frequency of microglia containing different levels of phagocytosed fluorescent bacteria. Right panel: phagocytosis in microglia with *Trem2* knockdown. N=3 independent experiments. Data shown as mean ± SEM. Paired *t*-test (paired within each culture preparation); * p<0.05, ** p<0.01, *** p<0.001, **** p<0.0001.

**Supplementary material Figure S5. Gene expression changes in primary microglia with acute *Trem2* knockdown.** The mRNA levels of individual genes at 72 hr post transfection were first normalized to *Rps28*, and then the relative fold change in the knockdown was calculated versus the negative control from the same batch of cell preparation (shown with a dotted line at 1). N=13 independent cell preparations. Data shown as mean ± SEM. One-sample *t*-test with Bonferroni correction; * p<0.05, ** p<0.01, *** p<0.001, ****p<0.0001.

**Supplementary material Figure S6. LPS suppression of *Trem2* expression in primary microglia results in little effect of *Trem2* knockdown on LPS-induced gene expression.** (A) The effect of LPS on *Trem2* expression with and without *Trem2* knockdown. Gene expression levels were normalized to *Rps28* and calculated as fold change relative to the negative control without LPS treatment in each individual culture preparation. (B) *Tnf* and *Il1b* expression, as examples of pro-inflammatory genes, were greatly up-regulated in primary microglia after LPS application. There was no significant difference between *Trem2* knockdown and negative controls. (C) ELISA analysis of the microglial supernatants showed that the secreted TNF-alpha levels in the medium were not significantly affected by *Trem2* knockdown. (D) Expression levels (RT-qPCR) of *C1qa, Cd68, Csf1r, Igf1, Pik3cg* and *Spi1* under the

LPS-stimulated conditions in *Trem2*-knockdown microglia compared to negative controls. N=3-6 (RT-qPCR) and 4 (ELISA) independent microglial preparations. Data shown as mean ± SEM. Two-way ANOVA with significant main effect of LPS incubation time and *Trem2* knockdown indicated as horizontal and vertical lines respectively. The only significant interactions between LPS treatment and *Trem2* knockdown were in *Trem2* expression (panel A) and *Igf1* expression (panel D), and so Sidak's *post hoc* tests were performed to test pairwise significance between the negative siRNA control and *Trem2* knockdown. * $p<0.05$, ** $p<0.01$, *** $p<0.001$, **** $p<0.0001$.

**Supplementary material Figure S7. Primary microglia from *Trem2* R47H KI mice show few changes in response to LPS.** (A) Expression of pro-inflammatory markers, *Tnf* and *Il1b.* (B) Gene expression of canonical genes. Gene expression was normalized to *Rps28* and calculated as fold change relative to the WT without LPS treatment. N=4-6 mice per genotype. Data shown as mean ± SEM. Two-way ANOVA with significant main effect of LPS-treatment and genotype indicated as horizontal and vertical lines respectively. A significant interaction between treatment and genotype was seen for *Il1b, Cd68, C1qa* and *Igf1* expression, and so Sidak's *post hoc* tests were performed to test pairwise significance between the genotypes within a treatment group, marked above individual groups. * $p<0.05$, ** $p<0.01$, *** $p<0.001$, **** $p<0.0001$.

**Supplementary material Figure S8. Expression profile of primary microglia from *Trem2* R47H KI and WT mice in response to LPS is similar whether normalized to *Rps28* or 3 housekeeping genes.** (A) Expression of *Trem2*. (B) Expression of pro-inflammatory markers, *Tnf* and *Il1b.* (C) Gene expression of canonical genes. Genes of interest were normalized to the geometric mean of 3 housekeeping genes: *H3f3b* (*H3 histone, family member 3B*), *Arf1* (*ADP-ribosylation factor 1*) and *Rps28,* and then calculated as fold change relative to the WT without LPS treatment. Compare to Supplementary material, Fig. S7. N=4-6 mice per genotype. Data shown as mean ± SEM. Two-way ANOVA with significant main effect of LPS-treatment and genotype indicated as horizontal and vertical lines respectively. A significant interaction between treatment and genotype was seen for *Trem2, Il1b, Cd68, C1qa* and *Igf1* expression, and so Sidak's *post hoc* tests were performed to test pairwise significance between the genotypes within a treatment group, marked above individual groups. * $p<0.05$, ** $p<0.01$, *** $p<0.001$, **** $p<0.0001$.

**Supplementary material Figure S9. The *Rps28* housekeeping gene showed stable expression in primary microglia treated with *Trem2* siRNA.** Primary microglia treated with *Trem2* siRNA or negative siRNA for 72 hr. (A) Total RNA assessed using capillary electrophoresis. Clear peaks for ribosomal RNA indicate RNA Integrity values (RIN) typically around 9.0 from 10.0 obtained regardless of siRNA treatment type. The RNA prepared from the *Trem2* siRNA-treated microglia showed increased levels of low molecular weight RNA species, consistent with reduced survival (see Supplementary material, Fig. S2), and altered metabolism of microglia with reduced *Trem2* activity, which is in agreement with previous studies with *Trem2* mutant mice and cells (35, 38, 39, 57). (B) *Rps28* levels (RT-qPCR) normalized to high molecular weight total RNA. Using the capillary electrophoresis plots we calculated the concentration of high molecular weight RNA using the area under the curve, and found that absolute *Rps28* expression levels were similar between the *Trem2* siRNA and negative siRNA treated cells. Hence many RT-qPCR experiments were plotted relative to *Rps28* expression. N=6 independent cell preparations for siRNA treatment. Data shown as mean ± SEM.

**Supplementary material Figure S10. The three housekeeping genes used in this study, *Rps28*, *H3f3b* and *Arf1* showed stable expression in primary microglia from *Trem2* R47H KI and WT mice treated with LPS and IL-4.** Primary microglia treated with LPS and IL-4 for 24 or 48 hr respectively as shown in Fig. 2. (A) Total RNA was assessed using capillary electrophoresis, and showed RIN values typically around 9.0 from 10.0 obtained regardless of genotype and treatment type. The RNA prepared from microglia from the *Trem2* R47H KI mice showed increased levels of low molecular weight RNA species, consistent with reduced survival (see Supplementary material, Fig. S2), and altered metabolism of microglia with reduced *Trem2* activity, which is in agreement with previous studies with *Trem2* mutant mice and cells (35, 38, 39, 57). Using the capillary electrophoresis plots we calculated the concentration of high molecular weight RNA using the area under the curve, and found that absolute *Rps28*, *H3f3b* and *Arf1* levels by RT-qPCR normalized to high molecular weight RNA were similar between the three treatment conditions and also between *Trem2* R47H KI and WT treated cells. Hence RT-qPCR results were very similar whether plotted relative to *Rps28* or all three housekeeping genes expression. N=6 independent cell preparations per genotype. Data shown as mean ± SEM. (B) RNA-seq gene expression levels for housekeeping genes *Rps28*, *H3f3b* and *Arf1* were similar between the three treatment conditions and also between *Trem2* R47H KI and WT treated cells, and thus similar to the RT-qPCR expression data. N=3 independent cell preparations per genotype. Data shown as mean ± SEM.

**Supplementary material Figure S11. Gene expression changes shown by microglia from *Trem2* R47H KI and WT mice in response to IL-4 via RNA-seq are similar to those seen by RT-qPCR.** The genes studied by RT-qPCR in Fig. 4-5 relative to *Rps28* show similar changes in gene expression by RNA-seq. N=3 independent microglial preparations. Data shown as mean ± SEM. Two-way ANOVA with significant main effect of IL-4-treatment and genotype indicated as horizontal and vertical lines respectively. A significant interaction between treatment and genotype was seen for only *Arg1* expression, and so Sidak's *post hoc* tests were performed to test pairwise significance between the genotypes within a treatment group, marked above individual groups. * $p<0.05$, ** $p<0.01$, *** $p<0.001$, **** $p<0.0001$.

**Supplementary material Figure S12. Biological annotations of co-expression genetic networks associated with *Trem2* expression and IL-4-treatment.** (A) Biological annotations associated with *Trem2* expression levels of primary microglia from HO-*Trem2* R47H KI and WT mice, with enrichment p-values corrected for multiple testing. Vertical dashed line signifies corrected enrichment p=0.05. (B) Biological annotations associated with IL-4-treatment using RNA-seq data from primary microglia treated with IL-4 versus untreated, with enrichment p-values corrected for multiple testing. Vertical dashed line signifies corrected enrichment p=0.05.
