## Supplementary figures for "*Trem2* promotes anti-inflammatory responses in microglia and is suppressed under pro-inflammatory conditions"

### Figure S1

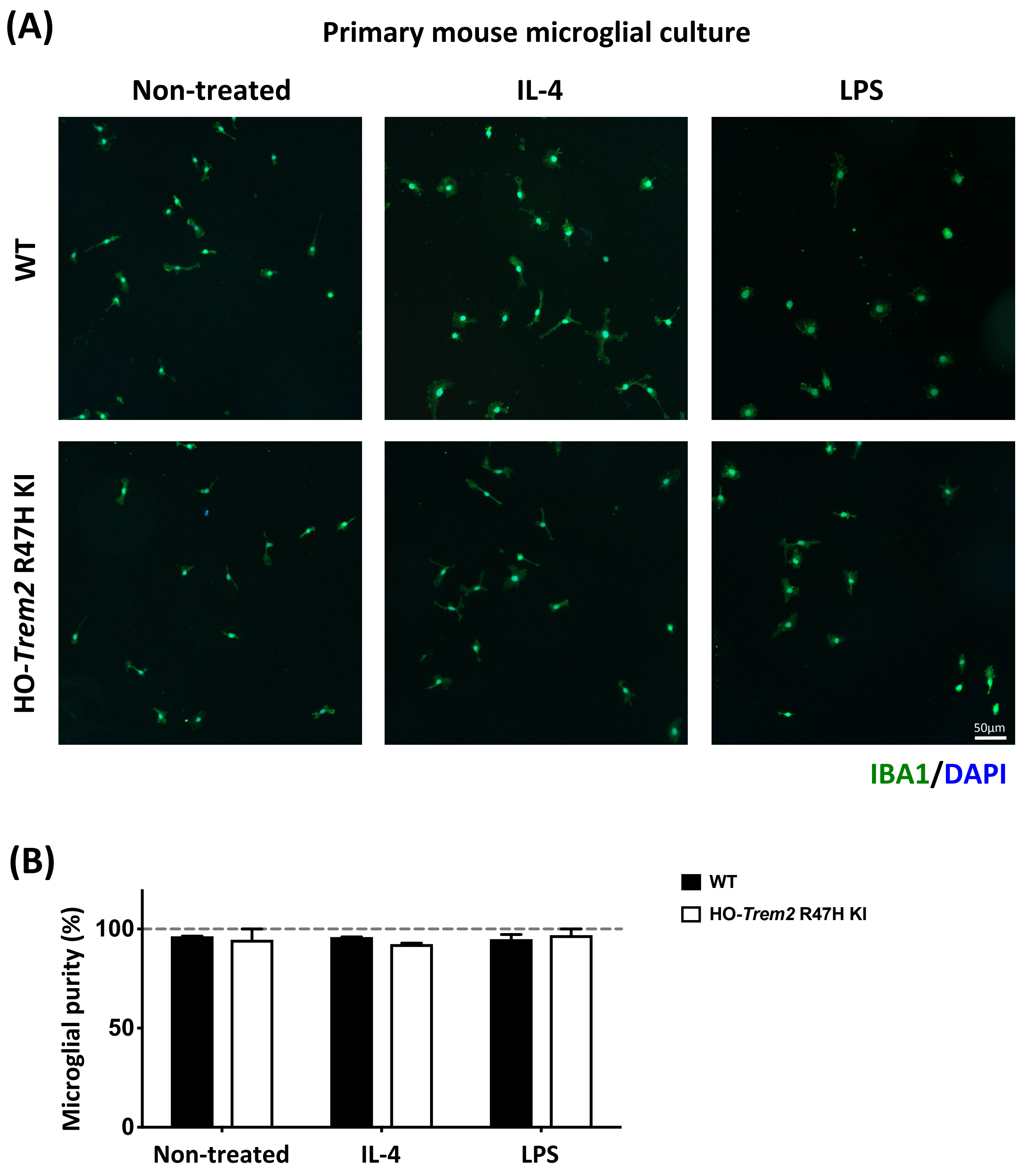
(A)
Primary mouse microglial culture
Non-treated
IL-4
LPS
WT
HO-Trem2 R47H KI
50µm
IBA1/DAPI
(B)
WT
HO-Trem2 R47H KI
Microglial purity (%)
100
50
0
Non-treated
IL-4
LPS

### Figure S2

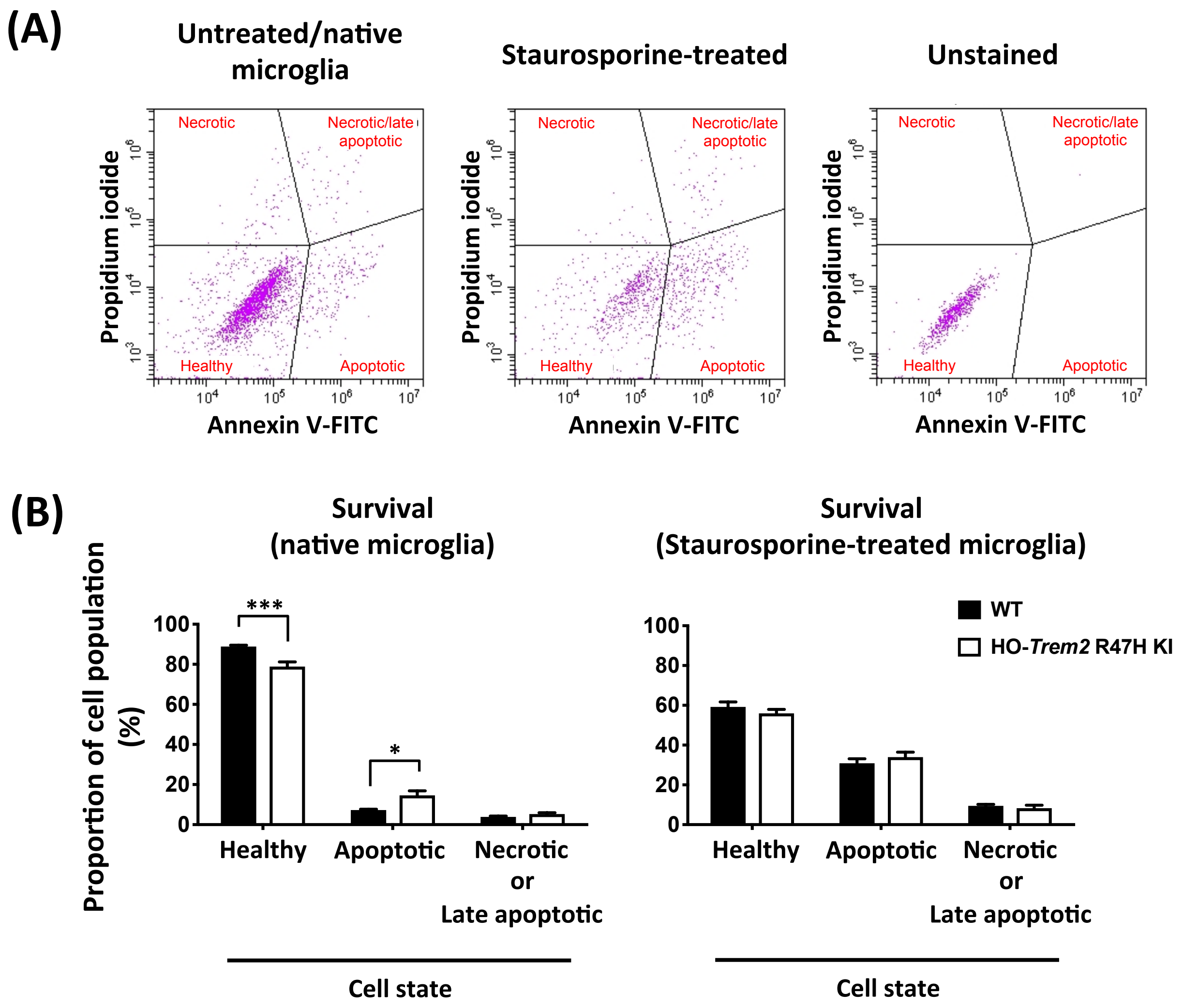
(A)
Untreated/native microglia
Staurosporine-treated
Unstained
Necrotic
Necrotic/late apoptotic
Healthy
Apoptotic
Propidium iodide
Annexin V-FITC
(B)
Survival (native microglia)
Survival (Staurosporine-treated microglia)
Proportion of cell population (%)
WT
HO-Trem2 R47H KI
Healthy
Apoptotic
Necrotic or Late apoptotic
Cell state
***
*

### Figure S3

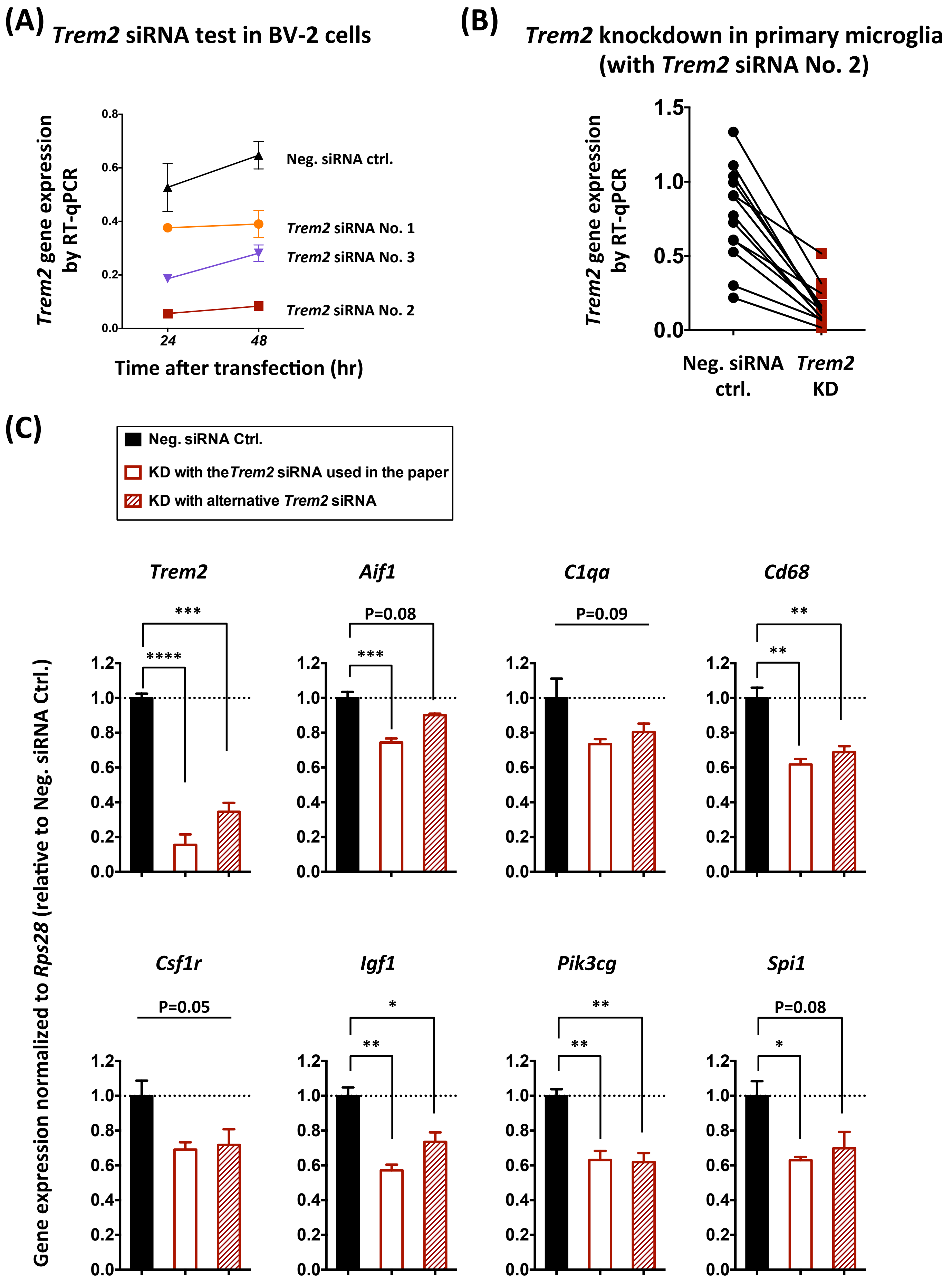

(A)
Trem2 siRNA test in BV-2 cells
Trem2 gene expression by RT-qPCR
0.0
0.2
0.4
0.6
0.8
24
48
Time after transfection (hr)
Neg. siRNA ctrl.
Trem2 siRNA No. 1
Trem2 siRNA No. 3
Trem2 siRNA No. 2
(B)
Trem2 knockdown in primary microglia (with Trem2 siRNA No. 2)
Trem2 gene expression by RT-qPCR
0.0
0.5
1.0
1.5
Neg. siRNA ctrl.
Trem2 KD
(C)
Neg. siRNA Ctrl.
KD with the Trem2 siRNA used in the paper
KD with alternative Trem2 siRNA
Gene expression normalized to Rps28 (relative to Neg. siRNA Ctrl.)
Trem2
***
****
Aif1
P=0.08
***
C1qa
P=0.09
Cd68
**
**
Csf1r
P=0.05
Igf1
*
**
Pik3cg
**
**
Spi1
P=0.08
*
0.0
0.2
0.4
0.6
0.8
1.0
1.2

### Figure S4

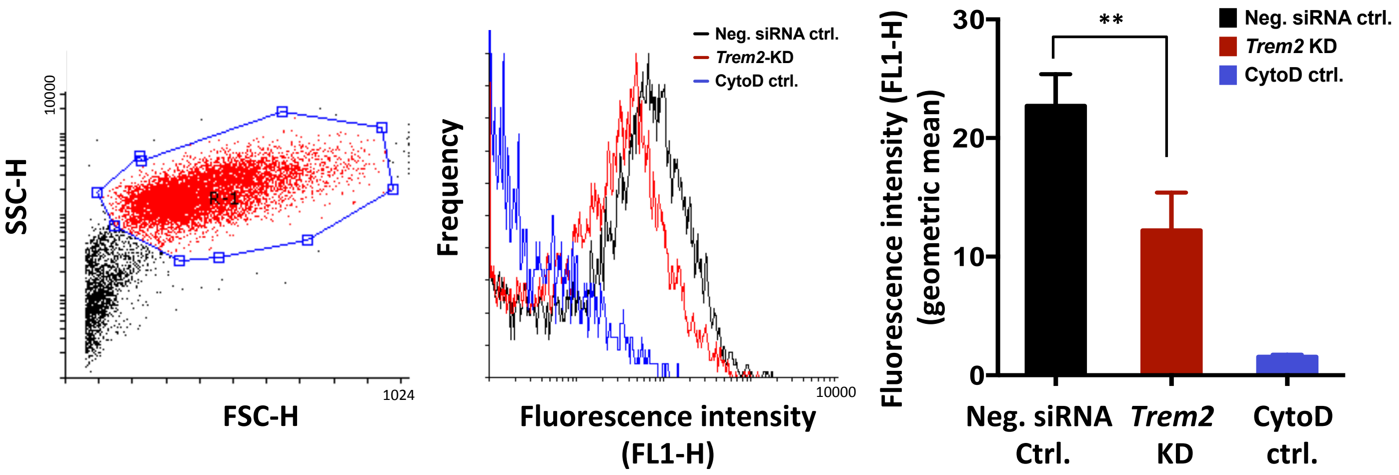
SSC-H
10000
FSC-H
1024
R-1
Frequency
Neg. siRNA ctrl.
Trem2-KD
CytoD ctrl.
Fluorescence intensity (FL1-H)
10000
Fluorescence intensity (FL1-H) (geometric mean)
30
20
10
0
**
Neg. siRNA ctrl.
Trem2 KD
CytoD ctrl.
Neg. siRNA Ctrl.
Trem2 KD
CytoD ctrl.

### Figure S5

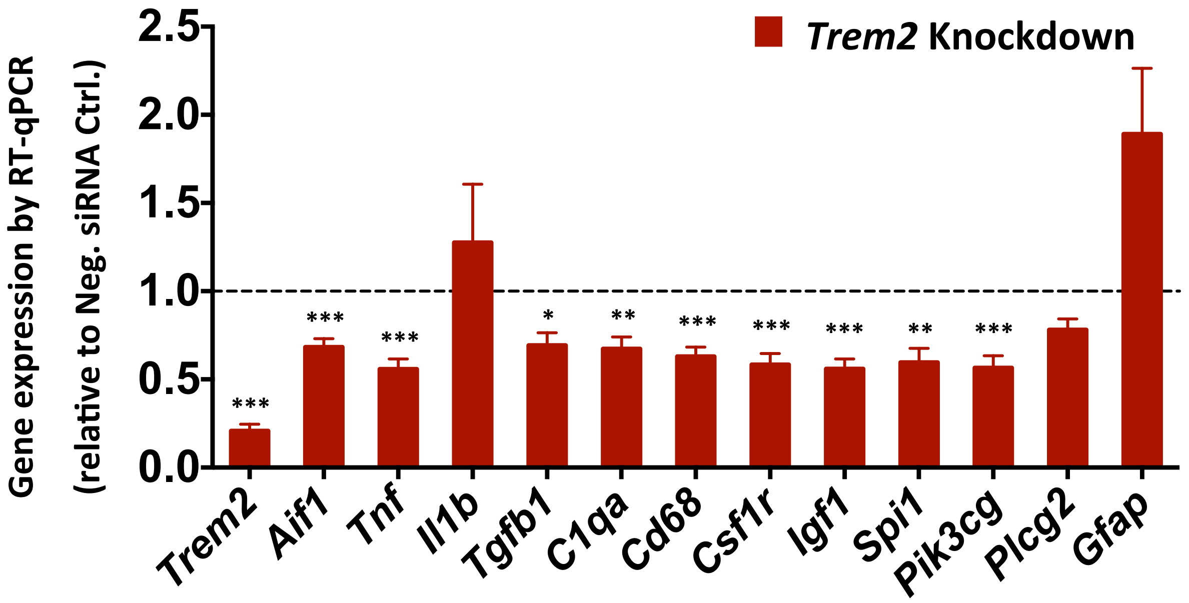

Trem2 Knockdown
Gene expression by RT-qPCR
(relative to Neg. siRNA Ctrl.)
2.5
2.0
1.5
1.0
0.5
0.0
***
***
***
*
**
***
***
***
**
***
Trem2
Aif1
Tnf
Il1b
Tgfb1
C1qa
Cd68
Csf1r
Igf1
Spi1
Pik3cg
Plcg2
Gfap

### Figure S6

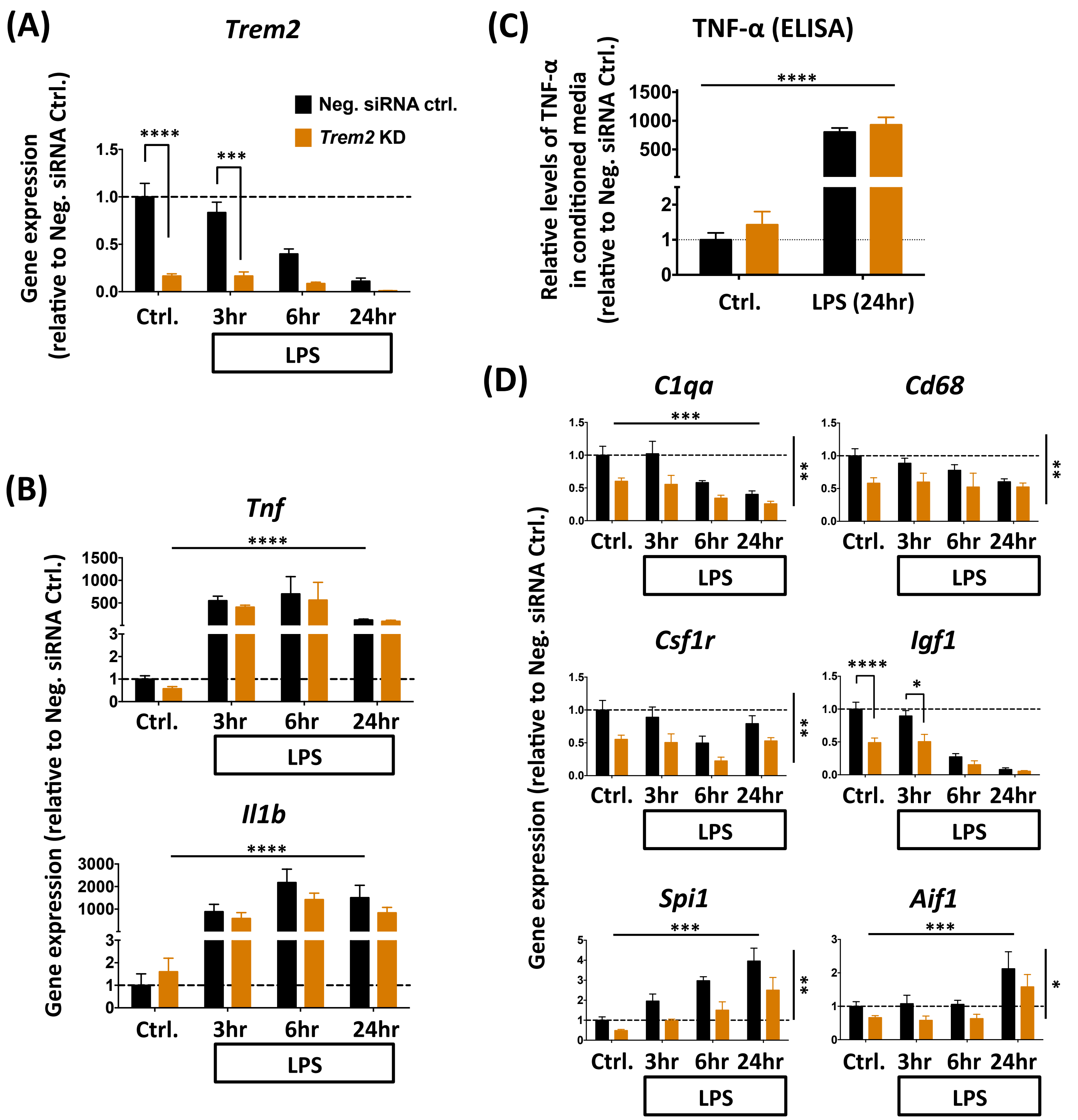

(A)
Trem2
Neg. siRNA ctrl.
Trem2 KD
Gene expression (relative to Neg. siRNA Ctrl.)
****
***
Ctrl.
3hr
6hr
24hr
LPS
(B)
Tnf
Il1b
Gene expression (relative to Neg. siRNA Ctrl.)
****
(C)
TNF-α (ELISA)
Relative levels of TNF-α in conditioned media (relative to Neg. siRNA Ctrl.)
****
Ctrl.
LPS (24hr)
(D)
C1qa
Cd68
Csf1r
Igf1
Spi1
Aif1
Gene expression (relative to Neg. siRNA Ctrl.)

### Figure S7

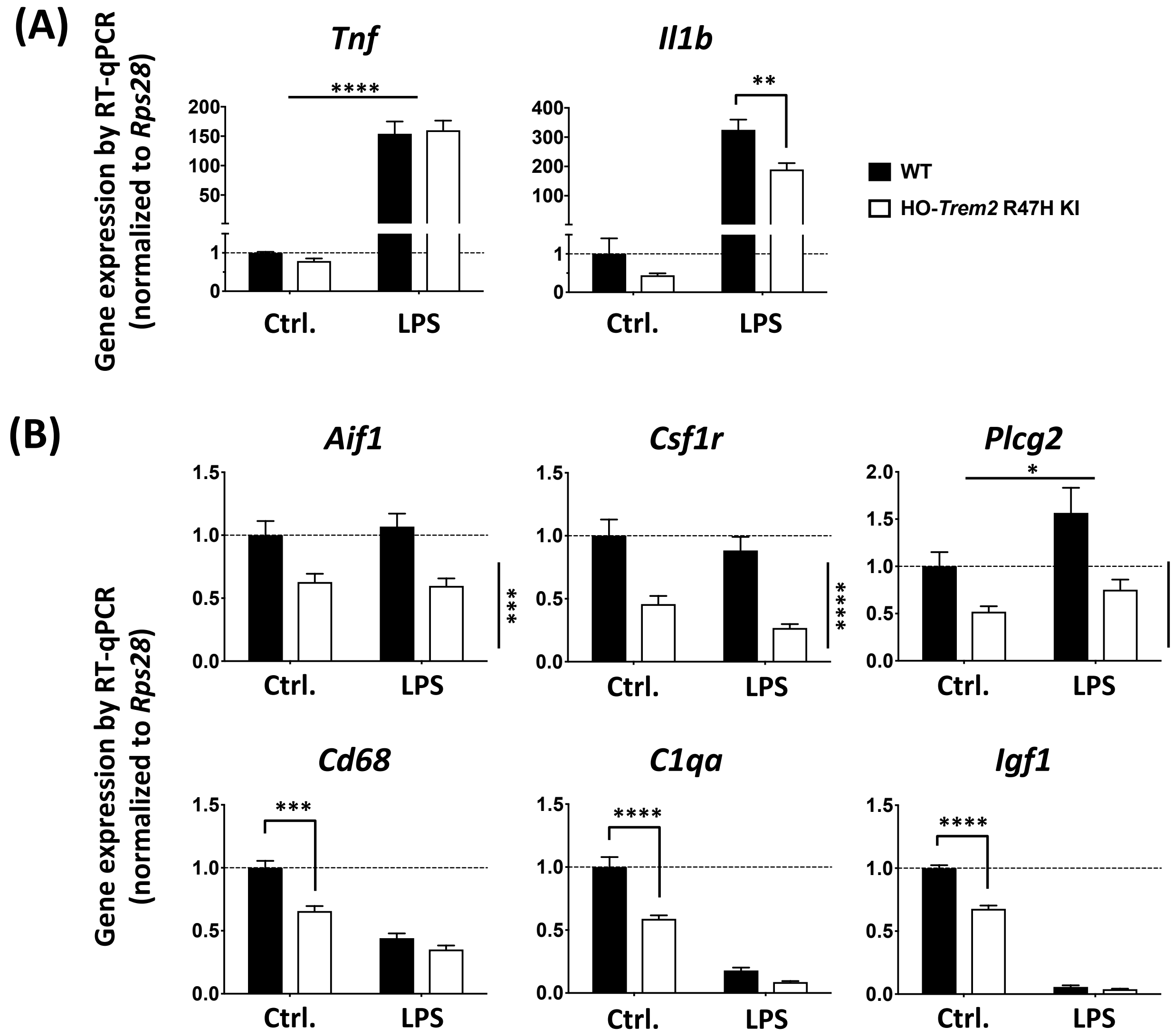

(A)
Tnf
Il1b
Gene expression by RT-qPCR (normalized to Rps28)
****
**
200
150
100
50
1
0
400
300
200
100
Ctrl.
LPS
WT
HO-Trem2 R47H KI
(B)
Aif1
Csf1r
Plcg2
Cd68
C1qa
Igf1
1.5
1.0
0.5
0.0
2.0
***
****
*
Ctrl.
LPS

### Figure S8

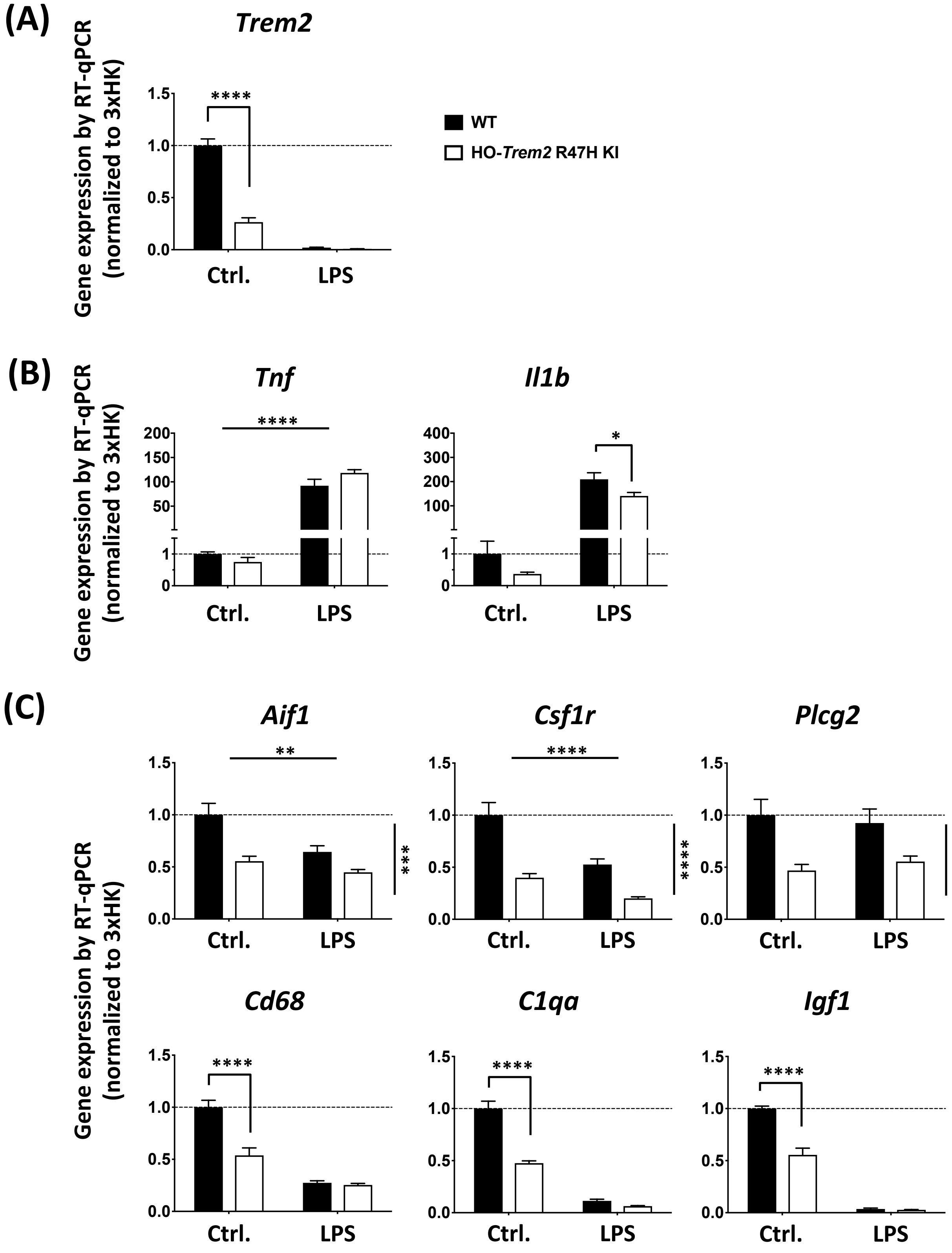

(A)
Trem2
Gene expression by RT-qPCR (normalized to 3xHK)
WT
HO-Trem2 R47H KI
Ctrl.
LPS
(B)
Tnf
Il1b
(C)
Aif1
Csf1r
Plcg2
Cd68
C1qa
Igf1

### Figure S9

(A)

**Neg. siRNA treated**

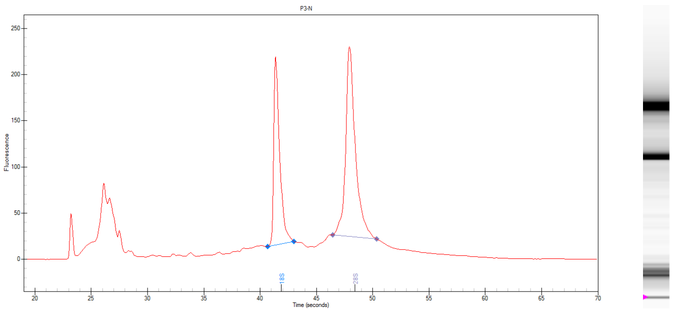

***Trem2* KD**

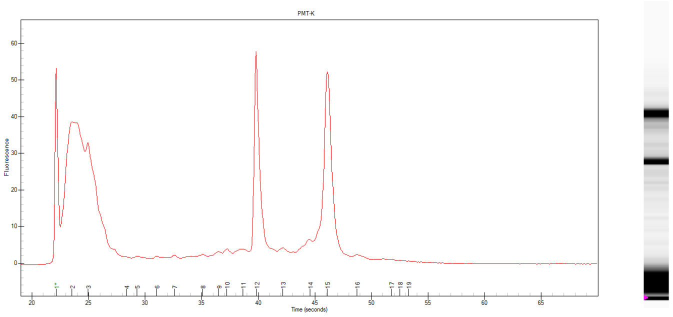

(B)

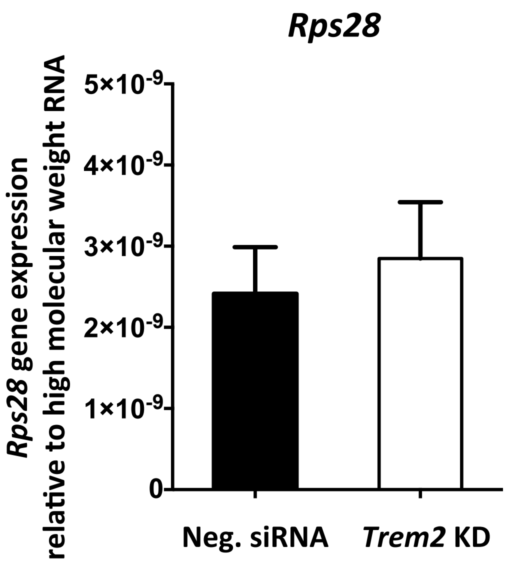

### Figure S10

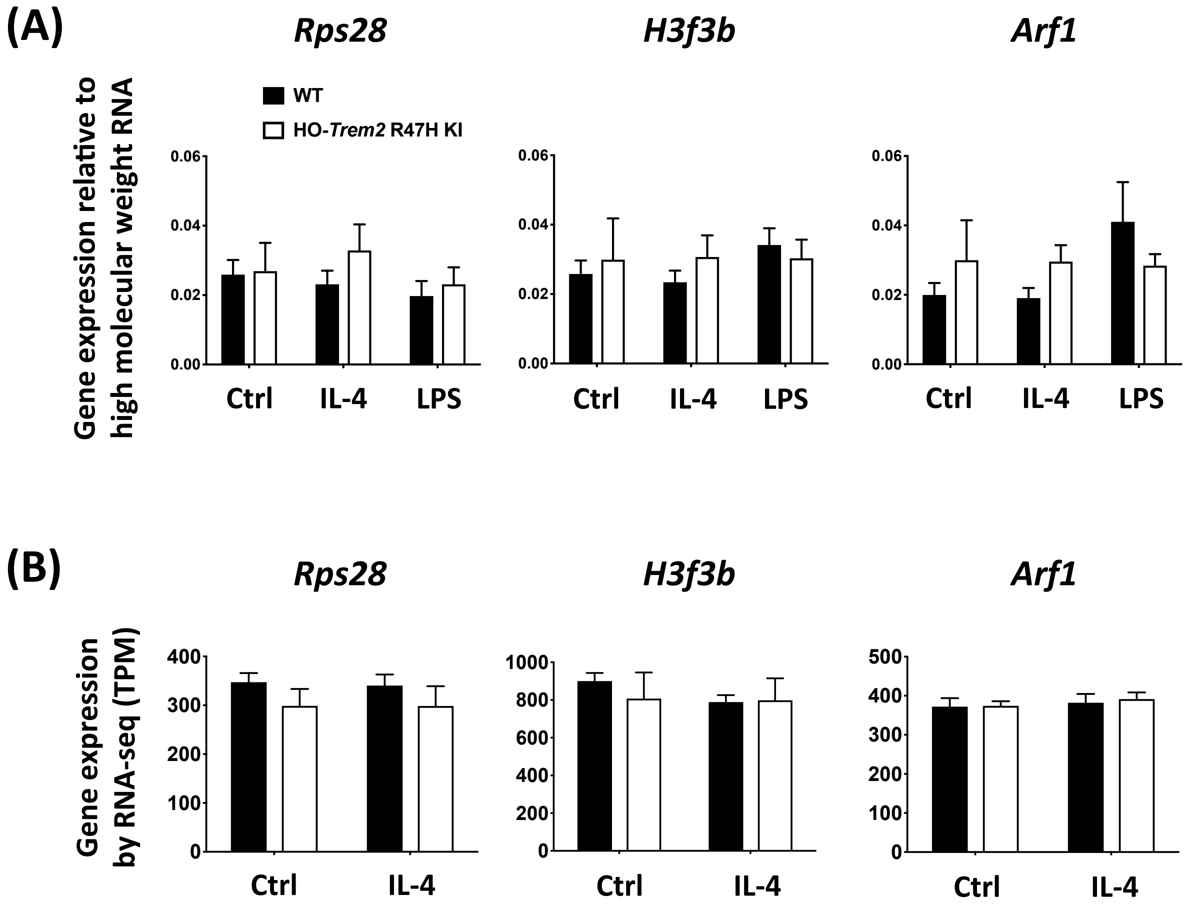
(A)
Rps28
H3f3b
Arf1
WT
HO-Trem2 R47H KI
Gene expression relative to high molecular weight RNA
0.06
0.04
0.02
0.00
Ctrl
IL-4
LPS
(B)
Rps28
H3f3b
Arf1
Gene expression by RNA-seq (TPM)
400
300
200
100
0
1000
800
600
400
200
0
500
400
300
200
100
0
Ctrl
IL-4

### Figure S11

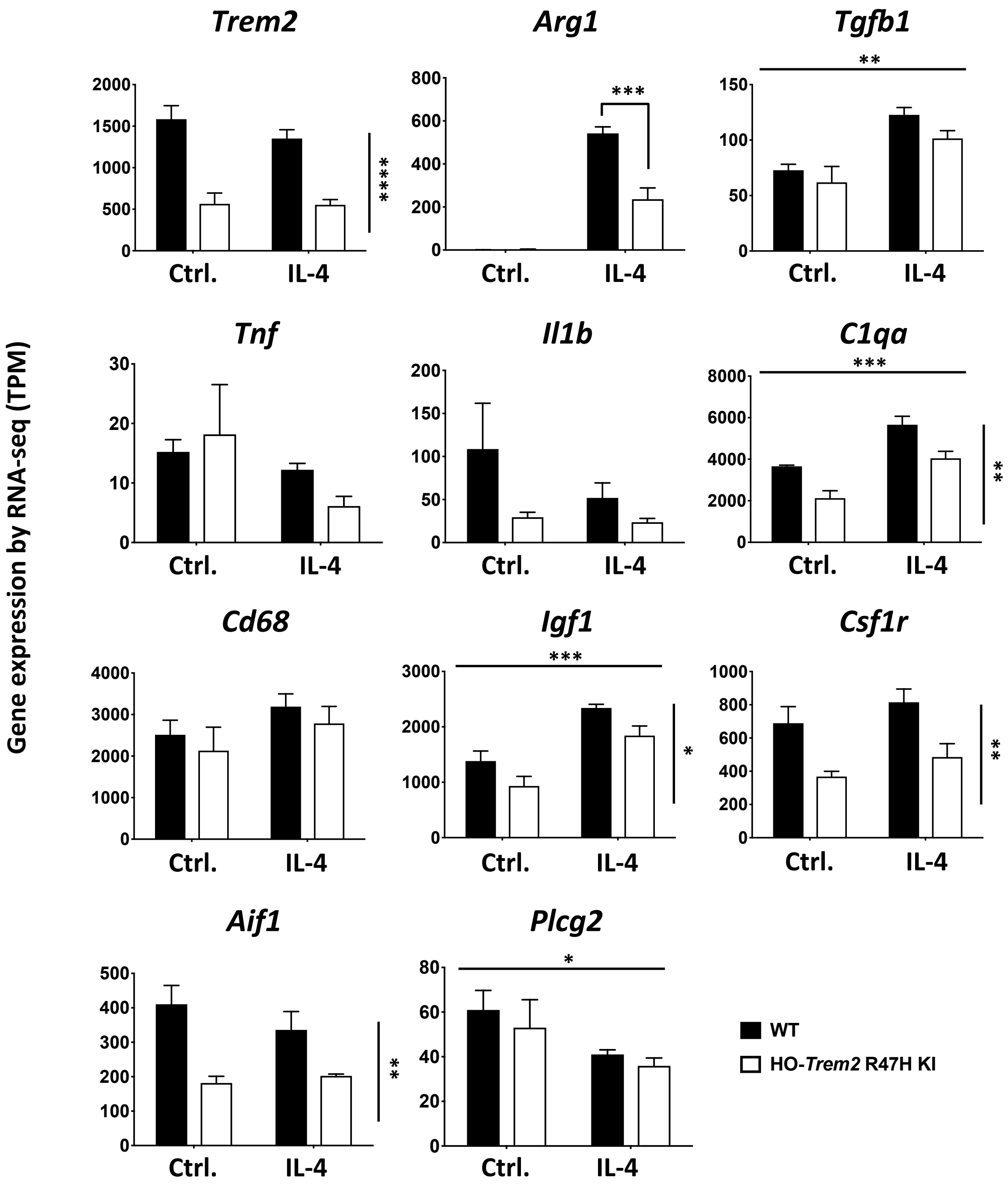
Gene expression by RNA-seq (TPM)
Trem2
2000
1500
1000
500
0
****
Ctrl.
IL-4
Arg1
800
600
400
200
0
***
Ctrl.
IL-4
Tgfb1
**
150
100
50
0
Ctrl.
IL-4
Tnf
30
20
10
0
Ctrl.
IL-4
Il1b
200
150
100
50
0
Ctrl.
IL-4
C1qa
***
8000
6000
4000
2000
0
**
Ctrl.
IL-4
Cd68
4000
3000
2000
1000
0
Ctrl.
IL-4
Igf1
***
3000
2000
1000
0
*
Ctrl.
IL-4
Csf1r
1000
800
600
400
200
0
**
Ctrl.
IL-4
Aif1
500
400
300
200
100
0
**
Ctrl.
IL-4
Plcg2
*
80
60
40
20
0
Ctrl.
IL-4
WT
HO-Trem2 R47H KI

### Figure S12

(A)

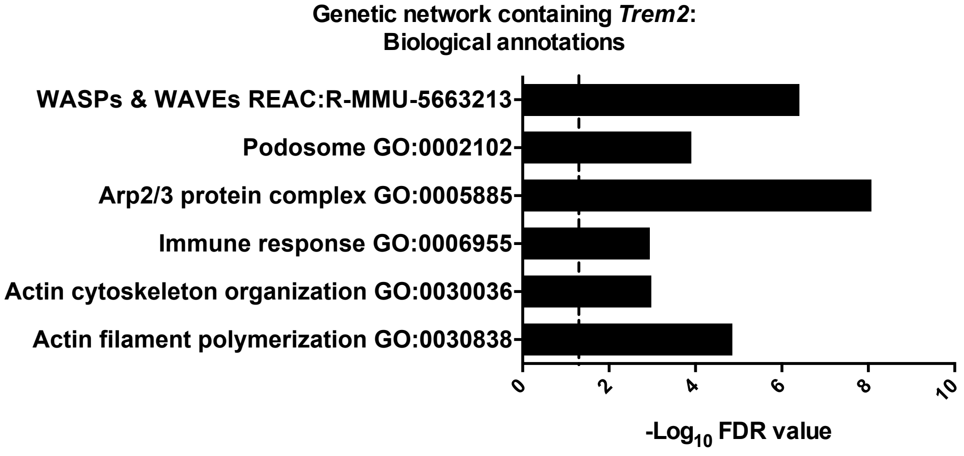


(B)

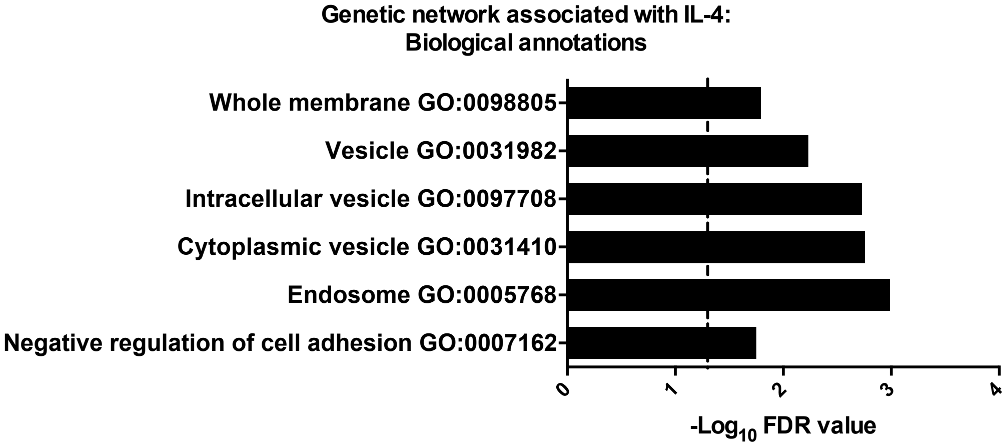
