## Supplementary material for "*Trem2* promotes anti-inflammatory responses in microglia and is suppressed under pro-inflammatory conditions": Table S6

**Table S6.** PCR primer sequences used in the study

| Gene | Forward primer (5’-> 3’) | Reverse primer (5’->3’) |
| --- | --- | --- |
| ***Aif1*** | GGAGACGTTCAGCTACTCTGAC | CATCCACCTCCAATCAGGGC |
| ***Arf1*** | TCCGAGATGCTGTTCTCTTGG | CAGTTCCTGTGGCGTAGAGAG |
| ***Arg1*** | ACCTGGCCTTTGTTGATGTCC | CTGGTTGTCAGGGGAGTGTTG |
| ***C1qa*** | GTCTCAAAGGAGAGAGAGGGGAG | TATGGACTCTCCTGGTTGGTG |
| ***Cd68*** | TTCACCTTGACCTGCTCTCTC | GTAGGTTGATTGTCGTCTGCG |
| ***Csf1r*** | GGATGGATACCAAATGGCCC | ACTGGTAGTTGTTAGGCTGC |
| ***H3f3b*** | CGTTCGTTGGTGGAGTATCTG | GTGGACTTCCTAGCGGTCTG |
| ***Igf1*** | GGACCGAGGGGCTTTTACTTC | CCAGTCTCCTCAGATCACAGC |
| ***Il1b*** | TCCTGTGTAATGAAAGACGGC | GGTGCTGATGTACCAGTTGGG |
| ***Pik3cg*** | CGGTCTATGCTGTACTGGGAG | GGAAGGGACATTCTCTGGAGG |
| ***Plcg2*** | AACTCTATTCCTCCTGTCGCC | GAACCTGCTGTTGCTTACTCTC |
| ***Rps28*** | ATCAAGCTGGCTAGGGTAACC | GGCCTTTGACATTTCGGATGA |
| ***Spi1*** | TATCAAACCTTGTCCCCAGCC | TGGTAGGTCATCTTCTTGCGG |
| ***Stat6*** | GCCAGATAACCATGCCCTTTG | TTGGTGAGGTCCTGTTCAGTG |
| ***Tgfb1*** | ATACCAACTATTGCTTCAGCTCC | CAGAAGTTGGCATGGTAGCCC |
| ***Tnf*** | ACCACGCTCTTCTGTCTACTG | CAGGCTTGTCACTCGAATTTTG |
| ***Trem2*** | GACCTCTCCACCAGTTTCTCC | TCAGAGTGATGGTGACGGTTC |
